# Modulation of T helper 1 and T helper 2 immune balance in a stress mouse model during *Chlamydia muridarum* genital infection

**DOI:** 10.1101/863753

**Authors:** Tesfaye Belay, Elisha Martin, Gezelle Brown, Raenel Crenshaw, Julia Street, Ashleigh Freeman, Shane Musick, Tyler Kinder

## Abstract

A mouse model to study the effect of cold-induced stress on *Chlamydia muridarum* genital infection and immune response has been developed in our laboratory. Our previous results show that cold-induced stress increases the intensity of chlamydia genital infection, but little is known about the effect of cold-induced stress on differentiations and activities of T cell subpopulations and bone marrow derived dendritic cells (BMDCs). The factors that regulate the production of T helper 1 (Th1) or T helper 2 (Th2) cytokines is not clear. The objective of this study was to examine whether cold-induced stress modulates the expression of transcription factors and hallmark cytokines of Th1 and Th2 or differentiation of BMDCs during *C. muridarum* genital infection in mice. Our results show that mRNA level of beta2-adrenergic receptor (β2-AR) compared to β1-AR and β3-AR was high in mixed population of CD4+ T cells and BMDCs. Further, decreased expression of T-bet and low level of interferon-gamma (IFN-γ) production and increased expression of GATA-3 and interleukin-4 (IL-4) production in CD4+ T of stressed mice was observed. Exposure of BMDCs to feroterol (β2-AR agonist) or ICI,118551 (*β*2-AR antagonist), respectively, revealed significant stimulation or inhibition of β2-AR in stressed mice. Moreover, co-culturing of mature BMDC and naïve CD4+ T cells resulted in increased production of IL-4, IL-10, and IL-17 in culture supernatants, suggesting that stimulation of β2-AR leads to the increased production of Th2 cytokines. Overall, our results show for the first time that cold-induced stress is able to modulate the pattern of Th1 and Th2 cytokine environment, suggesting that it promotes the differentiation to Th2 rather than Th1 by the overexpression of GATA-3 correlated with elevated production of IL-4, IL-10, IL-23, and IL-17 in contrast to a low expression of T-bet correlated with less IFN-γ secretion in the mouse model.

## INTRODUCTION

Chlamydia genital infection caused by *Chlamydia trachomatis*, the most common bacterial sexually transmitted disease (STD) worldwide [1] This infection, if left untreated, leads to the development of pelvic inflammatory disease (PID), fallopian tube scarring, ectopic pregnancy, infertility, and neonatal conjunctivitis [2, 3]. Epidemiologic data from the Centers for Disease Control and Prevention and World Health Organization indicate that more than 90 and 4 million new cases occur annually worldwide and, in the US, respectively [4]. In the United States, chlamydia genital infection disproportionately affects populations of low socio-economic status, particularly African-Americans [5, 6]. The reasons are not well known, but stress may have a major role in the persistent high rate of the disease associated with low socioeconomic conditions [7].

Several studies in animal models have demonstrated that anti-chlamydial T cell responses in the local genital mucosa play a role in the clearance of *C. trachomatis* from the genital tract [8–12]. It is known that immunity to *C. trachomatis* is mediated by T-helper (Th), specifically T-helper 1 (Th1), cells through interferon-gamma (IFN-*γ*) -dependent and -independent pathways [13–16].Moreover, other studies in human subjects have identified and characterized factors that may be involved in the development of immunopathogenesis after chlamydia genital infection caused by the human strain *C. trachomatis* [17, 18]. Those murine model studies have exhibited vital aspects of human genital chlamydial infection, including similar course of active infection, pathologic consequences or sequelae of an infection, tubal inflammation, hydrosalpinx formation and infertility [19–20]. It is well recognized that genital infection with *C. trachomatis* leads both to the recruitment of protective immune responses to the site of infection and infiltration of inflammatory products that may also contribute to pathological damage in the host [2, 21–23]. Repeated chlamydia genital infection in animal models [23–25] and human subjects [26, 27] is known to have a much greater risk of tubal obstruction than those with less exposure to *C. trachomatis*.

*Chlamydia muridarum*, previously known as the mouse pneumonitis strain of *C. trachomatis*, is commonly used for the study of chlamydia immunity, pathogenesis and vaccine development [28, 29]. However, significant differences in the biology and pathogenesis of infections caused *C. trachomatis* and *C. muridarum* have been reported [30]. Mouse strain-dependent studies show that *C. muridarum* produces shorter courses of infection in C57BL/6 and BALB/c mice than in C3H/HeN mice, even though more resistance to oviduct pathology was observed in C57BL/6 mice [31]. A study has demonstrated the successful infectivity and elicitation of pathologies with both human and murine strains of *Chlamydia* in mice [32]. It is mostly agreed that no model is perfect, extensive literature reports suggest that immunological findings made using *C. muridarum* infection in mice is not vastly different from those that occur with *C. trachomatis* infections in humans [33].

Psychological or physical stress resulting from hardship of life in society has major impacts on public health [34–38]. Glucocorticoids and catecholamines are stress hormones that serve as the major mediators of stress responses by modulating signaling events, which result in either immunosuppression or immunostimulation in the host [39–41]. Norepinephrine (NE) is one of the catecholamines of the sympathetic nervous system released during stressful conditions including cold-water application [39, 42, 43]. Studies document the existence of at least eight different alpha or beta subtypes of NE receptors [44]. Norepinephrine binds and stimulates the beta2 adrenergic receptor (β2-AR), which is predominantly expressed on CD4^+^ and B cells [45–47]. Studies have further shown that production and direct binding of NE to β2-AR modulates immune responses, including altering cytokine production, lymphocyte proliferation, and antibody secretion [47–51].

More studies have shown that dendritic cells (DCs) have an essential role in the induction of T cell responses to clear *C. muridarum* during genital infection [29, 52]. Furthermore, DCs pulsed with a recombinant chlamydial major outer membrane protein antigen elicit a CD4(+) type T-helper 2 (Th2) rather than type Th1 immune response that is not protective [53]. Other studies have shown that transcription factors T-box expressed in T cells (T-bet) and GATA-binding protein-3 (GATA-3) and play important a regulation role in the differentiation of naive Th cells towards Th1 or Th2 cells in human subjects and animal models [18, 54]. Specifically, modulation of cytokines and transcription factors (T-bet and GATA-3 in CD4+ T cells of chlamydia patients was reported [18], but the expression patterns of T-bet and GATA-3 in CD4+ T cells in our stress model during chlamydia genital infection yet is unknown.

Application of cold water as a stressor in animal models, including mice, has resulted in changes in immune responses that correlate with the activity of the neuroendocrine system of corticosteroids and catecholamines [55–60]. Although stress is implicated as a risk factor for various infections, its effect on chlamydia genital infection is unknown. We have previously shown that cold-water stress induces the production of catecholamines, which may play a critical role in the modulation of the immune system leading to increased intensity of *C. muridarum* genital infection [59, 60]. Furthermore, we have shown that supplement of NE to *in vitro* exerts an immunosuppressive effect on splenic T cell production of cytokines, decreased C. *muridarum* shedding in the genital tract of *β*1Adr/*β*2Adr KO mice, and severe pathology with a higher rate of infertility in *C. muridarum* infected mice [60]. However, our understanding of the expression of and function of Th1 and Th2 during cold-induced stress remains limited.

Here we sought to determine whether cold-stress treatment to mice could lead to imbalanced Th subsets during *C. muridarum* genital infection. Our overall hypothesis was that cold-stress treatment leads to the skewing CD4+ Th1 cell to Th2 cell function as determined by mRNA levels of T-bet and GATA-3 transcriptions factors and the level of signature cytokines produced during *in vitro* proliferation. We evaluated the differentiation and cytokine production ability of bone marrow derived dendritic cells (BMDCs) and its influence on the induction of Th cells upon co-culturing *in vitro*.

## MATERIALS AND METHODS

### Chlamydia stock culture and McCoy cell line

McCoy mouse cell line and stock culture of *Chlamydia muridarum (*previously known as *Chlamydia trachomatis* agent of mouse pneumonitis (MoPn) were kindly provided by Dr. Joseph Igietseme, then at Morehouse School of Medicine and currently at CDC, Atlanta, GA.

### Animals

Six- to seven-week-old female BALB/c strain mice were purchased from Hilltop Lab Animals, Inc. (Scottsdale, PA). Mice were housed in a vivarium at Bluefield State College (BSC) located in the Basic Science Building of the School of Arts and Sciences. Mice were given food and water *ad libitum* in an environmentally controlled room with equal daylight and night hours. all animal experimental protocols. This study was carried out in strict accordance with the recommendations in the Guide for the Care and Use of Laboratory Animals of the National Institutes of Health. The protocol was approved by Bluefield State College Institutional Animal Care and Use Committee (IACUC) (Assurance # A521-01) prior to beginning of the research. *Chlamydia muridarum* inclusion bodies was deposited into genital tract of each mouse under Ketamine-Xylazine induced anesthesia and all efforts were made to minimize animal suffering.

### Cold-water stressing protocol

Before the start of actual experiments, mice were acclimated for 7 days to relieve the stress of transport. Mice were then divided into three groups: stressed, handled/non-stressed and not handled/non-stressed. To induce chronic stress, five mice at a time were placed in a packet filled with four cm of cold water (4 to 5^°^ C) for five minutes daily for 24 days, as previously described [56]. The water level was deep enough to cover the backs of the mice and force them to swim in the cold water. At the end of each stressing period, mice were dried with towels to avoid hypothermia. Hereafter this group is referred to as the “test” or “stressed” group. To ensure that the stress was caused by the water treatment, and not simply by handling, the same protocol was applied, with the exception that the water was room temperature (22°C). A group of non-stressed mice were kept at room temperature without water treatment. Initial data on stress hormone production and *C. muridarum* course of infection showed no significant difference between the handled/non-stressed and the not-handled/non-stressed groups. For simplicity, the not-handled/non-stressed group was chosen as the control, and hereafter it is this group that is referred to by the terms “non-stressed” and “control “Throughout the study.

### Inoculation and isolation of Chlamydia muridarum

To regulate estrous cycles, all mice were injected subcutaneously with 2.5 mg of progesterone in 100 µL of phosphate buffered saline (PBS) on day 17 of the 24-day stressing period. On day 24, stressed and non-stressed mice were inoculated intravaginally with 10^5^ IFU of *C. muridarum* in a volume of 30 μL of PBS while under Ketamine-Xylazine induced anesthesia.

The course of infection was monitored by cervico-vaginal swabbing at 3-day intervals for the first 40 days of the primary course of infection. *C. muridarum* was recovered from swabs by staining infected monolayers of McCoy tissue culture with fluorescein isothiocyanate-labeled, genus-specific anti-chlamydial antibodies. Inclusion bodies in 10 fields/well were visualized, imaged, and counted under fluorescent microscope (Motic Inverted Microscope AE31E) and inclusion-forming units per mL (IFU/mL) was determined as previously descried (4).

### Purification of T cells from spleen and genital tract of mice

Spleens were removed aseptically then minced and teased with forceps in RPMI 1640 complete medium supplemented with 10% fetal bovine serum 1% penicillin/streptomycin, and 0.1% gentamicin (Sigma, St. Louis, MO). Spleen cell suspensions were pressed through a 70-μm cell strainer (Becton Dickinson, Franklin Lakes, NJ) to remove tissue debris. Splenocyte CD4+ T cells were isolated using EasyStep mouse T cell enrichment kits based on immunomagnetic negative selection, (STEMCELL Technologies, Vancouver, Canada) following the manufacturer’s instructions. Briefly, suspension of spleen cells was incubated with the EasySep Ab mixtures to non-CD4+ T cells followed by biotin selection mixture and magnetic beads. Labeled non-CD4+ T cells were removed using EasySep magnets. Genital tract lysates were produced by homogenization and lysates were pressed through a 70-μm cell strainer to purify CD4+ T cells as above.

### Antigen presenting cell preparation

Antigen Presenting Cells (APCs) from splenocytes of non-stressed mice were prepared by treatment with Mytomycin C. Briefly, splenocyte cells were washed by centrifugation at 300 x g for 10 minutes and freshly prepared Mytomycin C at the concentration of 25 μg per mL was added to 10^7^ splenocyte cells per mL and incubated at 37 °C for 30 minutes. Cells were washed four times as above and a final cell count of 5 x 10^5^ Mytomycin C-treated cells was prepared for proliferation assay.

### T cell proliferation assay

T cell proliferation *in vitro* was performed as described previously (7) with some modifications. Briefly, cells were seeded in triplicate at a density of 5 × 10^5^ cells mixed with 5 x 10^5^ Mytomycin C treated cells per well in a 96-well culture plate (Becton Dickinson). The cells were stimulated with 2.5 μg/mL of concanavalin A (Con A) cultured for 72 h in a water-jacketed incubator at 37°C and 5% CO_2_ (NuAire, Plymouth, MN). Culture supernatants were collected after 72 h of proliferation then stored at −86°C in preparation of measuring cytokine production using ELISA.

#### Testing the action of synthetic norepinephrine, beta 2-adrenergic receptor (β2-AR) agonist, β2-AR antagonists on T cell proliferation

We investigated the influence of synthetic norepinephrine bitartrate salt (NE), β2-AR agonist or antagonist on T cell cytokine production. Spleens were harvested as above from stressed/infected (SI), non-stressed/infected (NSI), and non-stressed/non-infected (NSNI) mice and isolated using Dynabeads® Magnetic Separation following the manufacturer’s instructions. Purified T cells were seeded at a density of 5 x 10^5^ per well in 96-well plates along with the same number of antigen presenting cells (APCs were exposed to NE (1 μM), fenoterol (β2-AR antagonist) (1 μM) and ICI 118,551 (β2-AR antagonist) (1 μM) for 1 h. Cells were proliferated for 72 h at 37° C in the presence/absence of Con A (5 μgmL-1), (0.3 μgmL-1) irradiated chlamydia antigen. Culture supernatants were collected after 72 h of proliferation and immediately stored at −20°C to be tested for the production of selected cytokines by ELISA.

#### Isolation and generation of bone marrow-derived dendritic cells (BMDCs) and bone marrow-derived macrophages (BMDMs)

To collect bone marrow cells, the femur and tibia bones were crushed using a mortar and pestle and debris was removed by passing the suspension through a 70 μm mesh nylon strainer. Cells were pipetted vigorously up and down to obtain single-cell suspension. Cells were washed once with complete RPMI 1640 and red blood cells were lysed using red cell lysis buffer from Sigma. Purified cells were cultured in a tissue culture flask containing 20 mL RPMI 1640 supplemented with recombinant murine Granulocyte-Macrophage Colony Stimulating Factor (GM-CSF) (10 ng/mL) (BioSource, Invitrogen cytokines & signaling, Camarillo, CA). At day 3 of the culture, half of the old culture was replaced with 10 mL of complete RPMI 1640 media and GM-CSF (10 ng/mL). At day 7, all floating cells and loosely adherent cells were collected as immature BMDCs, whereas the adherent cells were collected as BMDMs. All cells were counted in Trypan blue and their viability was over 80%.

#### Cytokine production and dendritic cell (DC) maturation upon lipopolysaccharide (LPS) stimulation

To examine the effect of β2-AR activation, immature dendritic cell (iDC) culture was pre-exposed to NE (1 μM), fenoterol (β2-AR antagonist) (1 μM) and ICI 118,551 (β2-AR antagonist) (1 μM) for 1 h before stimulation with LPS (5 μg/mL) for 24 h. Supernatants were collected, and the production of different cytokines was assessed using an ELISA kit.

#### Co-culturing of Naïve CD4+ T cells and LPS-stimulated BMDCs *in vitro*

To examine the effect of BMDCs on T cell proliferation, freshly isolated CD4+ T cells isolated from stressed and non-stressed *Chlamydia muridarum*-infected mice were co-cultured with matured BMDCs for four days in 96-well plates at BMDC: T cell ratio of 1:10. Culture supernatant was collected at day four of co-culturing and cytokine production was determined by ELISA.

#### Priming Th1 or Th2 *in vitro* under polarizing conditions

Naïve CD4 + T cells from stressed and non-stressed groups of mice were isolated using the mouse naïve CD4+ T cell isolation kit for activation with polarizing milieu. Briefly, splenic naïve CD4+ T cells were seeded at 1 x 10^5^ cells/well of a plate coated with 1 μg/mL anti-CD3 and anti-CD28 and incubated for 72 h in the presence of recombinant murine interferon-gamma (IFN-γ) (10 ng/mL) recombinant murine IL-12 (10 ng/mL) and anti-mouse IL-4 (100 ng/mL) for Th1 polarization or recombinant IL-4 (100 ng/mL) anti-mouse IFN-γ (10 ng/mL) and anti-mouse IL-12 (10 ng/mL) for Th2 polarization. The CD4+ T cells were stimulated again for another four days in coated plates and fresh medium. Culture supernatants were collected to determine the polarization of IL-4 and IFN-γ in stressed and non-stressed CD4+ T cell samples as determined by ELISA

### Cytokine measurement using ELISA

The level of cytokines in supernatants collected from in vitro proliferating CD4+ T cells and bone-marrow derived dendritic cells (BMDCs) of different treatment groups was measured using an ELISA kit (Invitrogen, Camarillo, CA) following the manufacturer’s instructions as previously described (59). Briefly, standard samples and test samples were diluted and added to wells in duplicate and incubated as directed. Optical densities were measured at 450 nm with an ELISA plate reader from Biotech (Winooski, VT). The concentration of the cytokine in each sample was obtained by extrapolation from the standard calibration curve generated in each assay.

#### RNA isolation, cDNA synthesis and polymerase chain reaction (PCR) analysis

Total RNA was isolated from cultured splenocyte and genital tract T cells, BMDCs using the FastPrep24™ or FastPrep® FP120 Instrument and Fast RNA® Pro Green Kit (MP Biomedicals) following the manufacturer’s instructions. Briefly, total RNA was released into the protective RNApro™ solution by homogenizing samples in 2 mL tubes using Lysing Matrix D. Total RNA Extraction was completed using chloroform and precipitation. Then the isolated RNA was quantified using a Theme Scientific Nanodrop (Thermofisher). RNA sample values ranging from 1.9 to 2.1 at 260/280 and > 2.0 at 260/230 ratio were used for cDNA synthesis.

#### cDNA synthesis and quantitative real time PCR (qPCR) analysis

Oligo primers of target molecules for mRNA determination were purchased from Integrated DNA Technologies (Skokie, IL). First-strand cDNA for gene expression analysis using real-time qPCR was carried out using the iScript cDNA synthesis kit from Bio-Rad following the manufacturer’s instructions. Briefly, a volume of 4 µL of 5x iScript reaction mix and 1 µL of iScript reverse transcriptase and 1 µg of total RNA was mixed in clean Eppendorf tubes. Each tube was then brought to a total of 20 µL using RNase/DNase-free water. The tubes were then incubated at 25°C for 5 minutes, for 30 minutes at 42°C, and for 5 minutes 85°C. The tubes were placed at −20°C until usage in qPCR, which was conducted using 2x iTaq SYBR Green Supermix of Bio-Rad following the manufacturer’s instructions. Briefly, 10 µL of the iTaq Universal SYBR Green Supermix, 4 µL of the forward and reverse primer mix for each gene, 4 µL of RNase/DNase free water and 2 µL of the cDNA samples were added to PCR strips containing 8 wells with two replicates for each sample per plate. The strips were then placed in a Bio-Rad CFX96 PCR machine set to run a 3-step cycle.

#### Sequences of oligo primers used in the study

Mouse T-bet forward: 5’-TCAACCAGCACCAGACAGAG-3’ reverse 5’-AACATCCTGTAATGGCTTGTG-3’

Mouse GATA-3 forward: 5’-CTTATCAAGCCCAAGCGAAG-3’, reverse: 5-CCCATTAGCGTTCCTCCTC-3’

Mouse β1AR forward: 5’-AAACTCTGGTAGCGAAAGGGGAC-3’ reverse 5’ TCTGCTCATCGTGGTGGGTAAC-3’

Mouse β2AR forward: 5’-AGCCGTTCCCATAGGTTTCG-3’ reverse 5’-CGTCCTCGATTGTGTCTTTCTACG-3’

Mouse β3AR forward: 5’-CGAAGAGCATCACAAGGAGGG-3’ reverse 5’-CGAAACTGGTTGCGGAACTGTGT-3’

Mouse IFN-gamma forward: 5’-GCCATCAGCAACAACATAAGC-3’, reverse: 5’-TGAGCTCATTGAATGCTTGG-3’

Mouse IL-4 forward: 5’-CATCGGCATTTTGAACGAG-3’ reverse 5’-CGAGCTCACTCTCTGTGGTG-3’.

#### qPCR data analysis

Quantitative PCR was performed using BIO RAD iTaq™ Universal SYBR® Green Supermix following the manufacturer’s instructions on a CFX96 Real-Time System. Analysis was performed using BioRad CFX Manager Software with GAPDH as a housekeeping gene for reference. Briefly, ITaq™ Universal SYBR®. We compared the cycle threshold (Ct) required for the fluorescent signal to cross the threshold that exceeds background level. Threshold level and Ct value on a real-time PCR amplification curve are shown in results section. The Ct values from each sample were also used to determine the fold change using the Livak method. To determine the fold change using this method, the following equations were used:

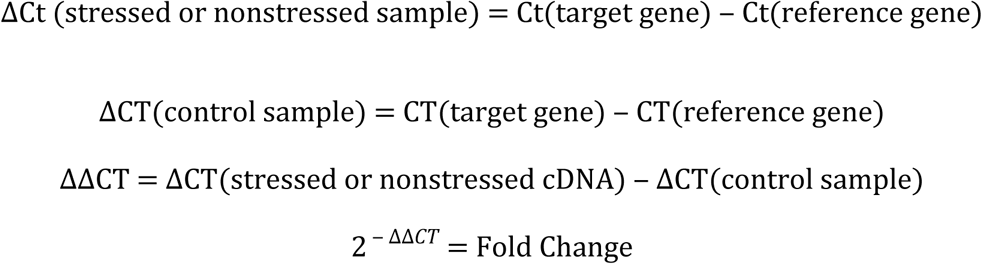

The target genes in the sample refer to genes of interest, such as cytokines. The reference gene refers to the GAPDH gene used to normalize the CT value of the sample. Using this equation, the fold change of the mRNA of test and control samples were determined. Some of qPCR products were further analyzed by running on gel electrophoresis.

The ΔΔCT method was used to calculate relative changes in gene expression of target genes determined from real-time quantitative PCR experiments. Expression in stressed and non-stressed was normalized to the internal control gene, GAPDH and relative to the non-stressed sample as a control. One representative of two independent experiments is shown. Comparison was at the level of *P <0.05*.

#### Western blotting of analysis

Protein from mouse genital tract lysate, CD4+ T cells or dendritic cells were isolated using the FastPrep-24™ or FastPrep® FP120 Instrument and FastRNA® Pro Green Kit (MP Biomedicals) following the manufacturer’s instructions. Purified protein was denatured and separated by sodium dodecyl sulfate polyacrylamide gel electrophoresis (SDS-PAGE) in a Criterion™ Cell and transferred to a nitrocellulose membrane in a Criterion™ Blotter (Bio-Rad) following manufacturer’s instructions. Band formation was detected via an Amplified Opti-4CN Substrate Kit following the manufacturer’s instructions. Relative expression was calculated by the ΔΔCt method to compare expression levels of the same transcript in different samples or by the ΔCt method to compare expression levels of several transcripts in the same sample using ChemiDOC molecular imager and software (BioRad) following the manufacturer’s instructions. Equivalent protein loading and transfer efficiency were verified by using β−actin as a control.

#### Establishing interaction networks of gene and proteins

Cytoscape is a social network of open source software that is used to visualize molecular interaction networks and biological pathways. Data of gene expression and cytokine production in stressed and non-stressed mice with or without *Chlamydia muridarum* genital infection were collected for interaction with other important genes or proteins extracted from data base. Related protein networks of up-regulated or down-regulated genes or proteins were visualized using the cystoscope software.

### Statistical analysis

All data are expressed as a mean+/- standard errors of the mean (SEM) unless otherwise stated. Statistical significance between any two groups were tested using Student’s t-test and ANOVA was used to test a statistical difference between more than two groups. Statistical significance was at the level of *P < 0.05*.

## RESULTS

### Cold water-induced stress increases the intensity of *Chlamydia muridarum* genital infection during primary infection

To determine the level of infection in stressed and non-stressed mice along with immune response analysis, we measured the number of viable *C. muridarum* present on vaginal swabs collected every 3 to 6 days post infection. As shown in **Table 1**, there are greater numbers of chlamydia inclusion counts in the swabs of stressed mice compared to non-stressed mice, indicating that cold-induced stress increases the intensity of *C. muridarum* genital infection in mice. Our results are consistent with our previous findings of increased *C. muridarum* shedding in regions of the genital tract of stressed mice compared to non-stressed mice [59, 60].

**Table 1:** Course of *Chlamydia muridaruim* genital infection in stressed and non-stressed mice as measured by quantitation of viable organisms shed from the genital tract.

| **Days post infection** | **Stressed mice** | ***Non-stressed*** |
| --- | --- | --- |
| 3 | $5.1 \times 10^4 \pm 25^a$ | $3.0 \times 10^3 \pm 21$ |
| 9<sup>b</sup> | $6.4 \times 10^5 \pm 12$ | $1.9 \times 10^4 \pm 43$ |
| 12<sup>b</sup> | $4.1 \times 10^3 \pm 20$ | $4.1 \times 10^2 \pm 20$ |
| 15 | $4.1 \times 10^3 \pm 20$ | $2.2 \times 10^1 \pm 20$ |
| 21 | $2.5 \times 10^2 \pm 20$ | $3.1 \times 10^1 \pm 20$ |
| 28 | $1.1 \times 10^1 \pm 15$ | Non-detectable |
| 34 | Non-detectable<sup>c</sup> | Non-detectable |
| 42 | Non-detectable | Non-detectable |

<sup>a</sup> Each data is mean and standard deviation of inclusion forming units/milliliter determined from McCoy cell culture representing combined results of two separate experiments (n=5 mice per group). Data are a mean ± SEM of two or more independent experiments ran on different dates using the method previously described (60).
<sup>b</sup>A statistical difference in counts was observed between stressed and non-stressed mice on day 9 and 15 after infection ( $P < 0.05$ ).
<sup>c</sup>No *Chlamydia muridarum* count in non-stress mice and stressed mice thereafter indicating the resolution of infection.

### Cold-induced stress results in differential gene expression of beta-adrenergic receptor (β-AR) subtypes in mouse genital tract T cells during *Chlamydia muridarum* infection

This experiment was to determine the gene expression levels of β-adrenergic receptor (β-AR) subtypes in splenic CD4+ T cells of stressed and non-stressed mice during *C. muridarum* genital infection. We hypothesized that differential gene expression of β-AR subsets may play an important role in modulating the immune function against chlamydia genital infection in our stress mouse model. Real-time quantitative PCR based on fluorescence detection and threshold cycle (Ct) values comparison was used in measuring mRNA level of β-AR subsets. The basis for analysis was that Ct is inversely proportional to the initial copy number of mRNA or a higher initial copy number of mRNA correlates to a lower Ct value. As shown in the amplification curve (**Figure 1A**), Ct values of β2-AR in splenic CD4+T cells of stressed mice was about five times lower than that of non-stressed mice. In contrast, the amplification curve of β1-AR and β3-AR of splenic CD4+T cells showed no difference between stressed and non-stressed mice (**Figure 1B, 1C**). Calculation of fold-changes and *p-*values on the expression of β-AR subsets was performed. The relative expression of β2-AR in splenic CD4+T cells compared to that of non-stressed mice showed the highest fold-changes (p <0.05). On the other hand, the relative expressions of β1-AR and β3-AR in splenic CD4+ T cells showed no significant difference between stressed and non-stressed mice (data not shown).

**Figure 1:**
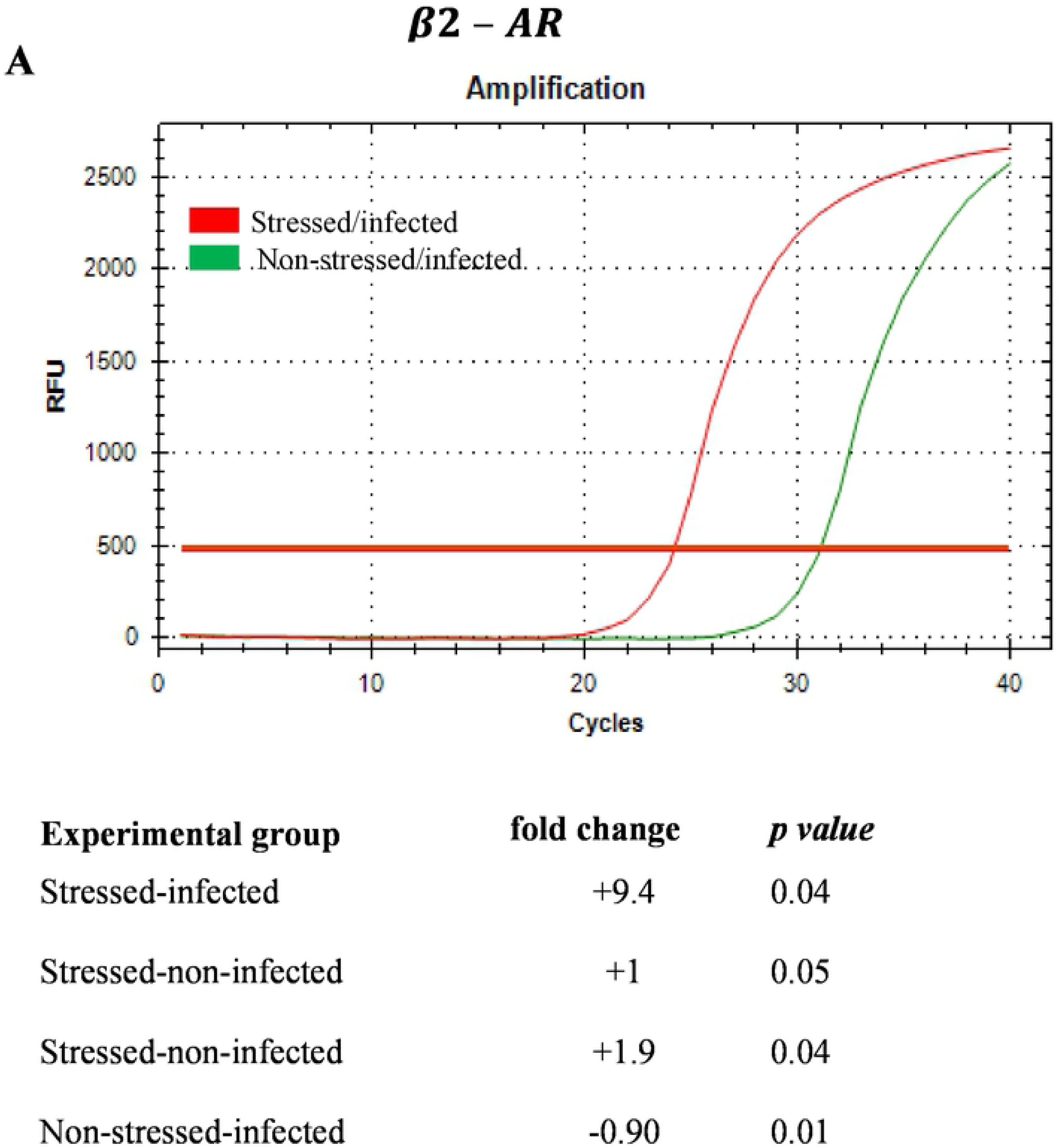
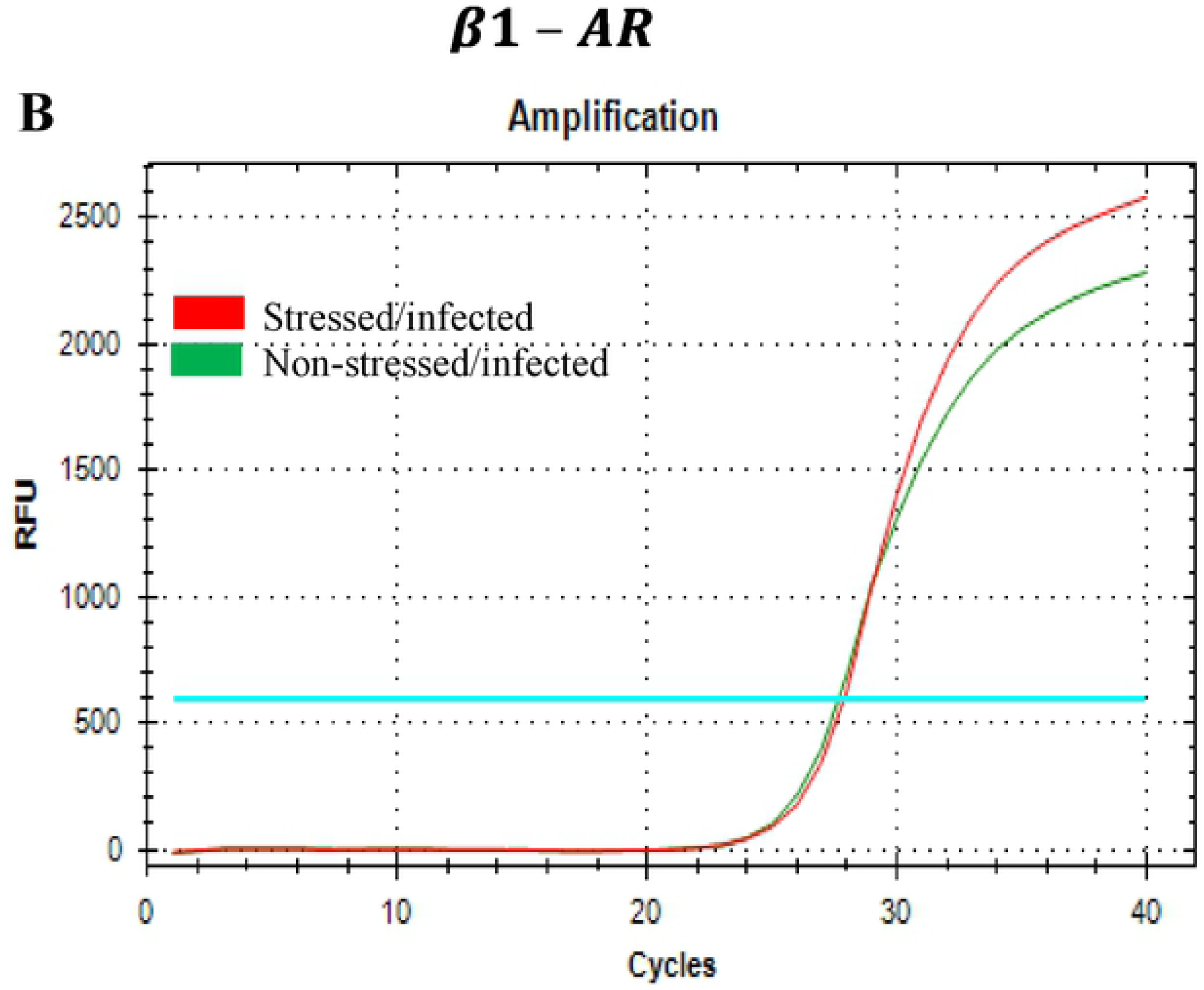
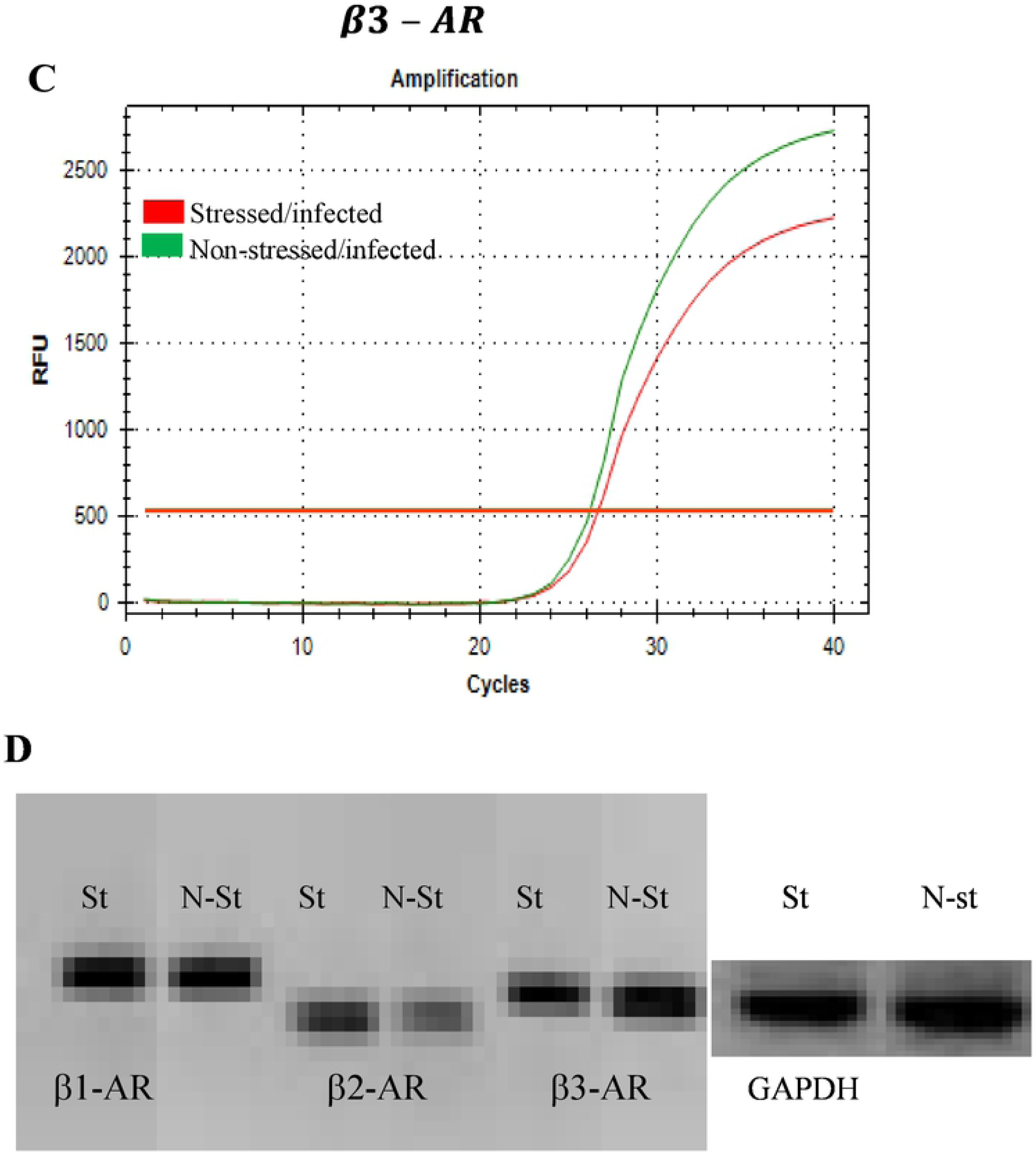
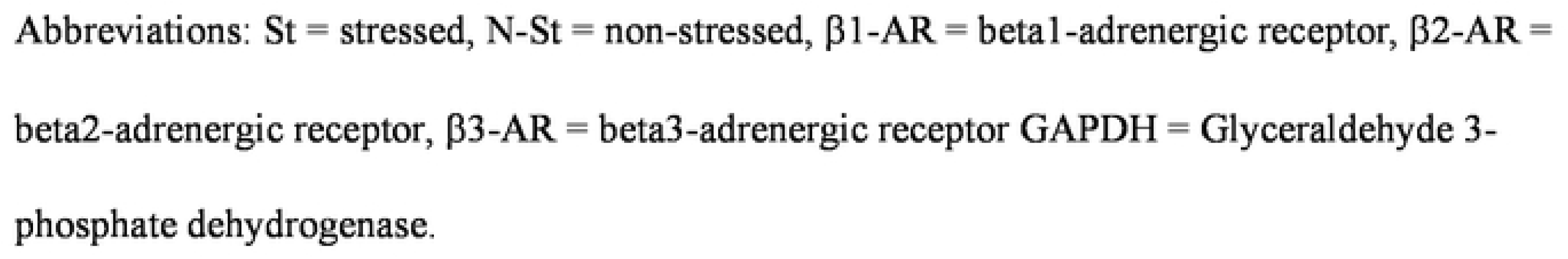
Gene expression profiles of beta-adrenergic receptor (β-AR) subsets in splenic T cells of stressed and non-stressed mice during *Chlamydia muridarum* genital infection. Amplification cures along with fold-changes of mRNA levels of β2-AR subsets normalized to the housekeeping gene, GADPH are shown. (**A**) β2-AR, (**B)** β1 –AR, (**C**) β3-AR and (**D**) PCR-products of β2-AR subsets are displayed by gel electrophoresis. Fold-changes mRNA levels of β1 –AR and β3-AR showed no significant difference between stressed and non-stressed group (data not shown). ***Abbs***: **St** = stressed, **Non-St** = non-stressed. Data shown are a representative of two or more independent experiments ran on different dates.

In another set of experiments, CD4+ T cell populations were enriched from the genital tract of stressed and non-stressed mice to examine gene expression profiles of the β-AR subsets. Similar to splenic CD4+ T cells, there was an increased mRNA level of β2-AR in the genital tract CD4+ T cells stressed mice compared to that of non-stressed mice (**Figure 2A**). The patterns of gene expressions of β1-AR and β3-AR of CD4+ T cells showed no statistical differences in stressed and non-stressed mice (**Figure 2B, 2C**). DNA intensity of β2-AR PCR end product was slightly higher in stressed mice than non-stressed mice, whereas of β1-AR and β3-AR PCR products show no difference in DNA intensity between stressed and no-stressed groups (**Figure 2D**).

**Figure 2:**
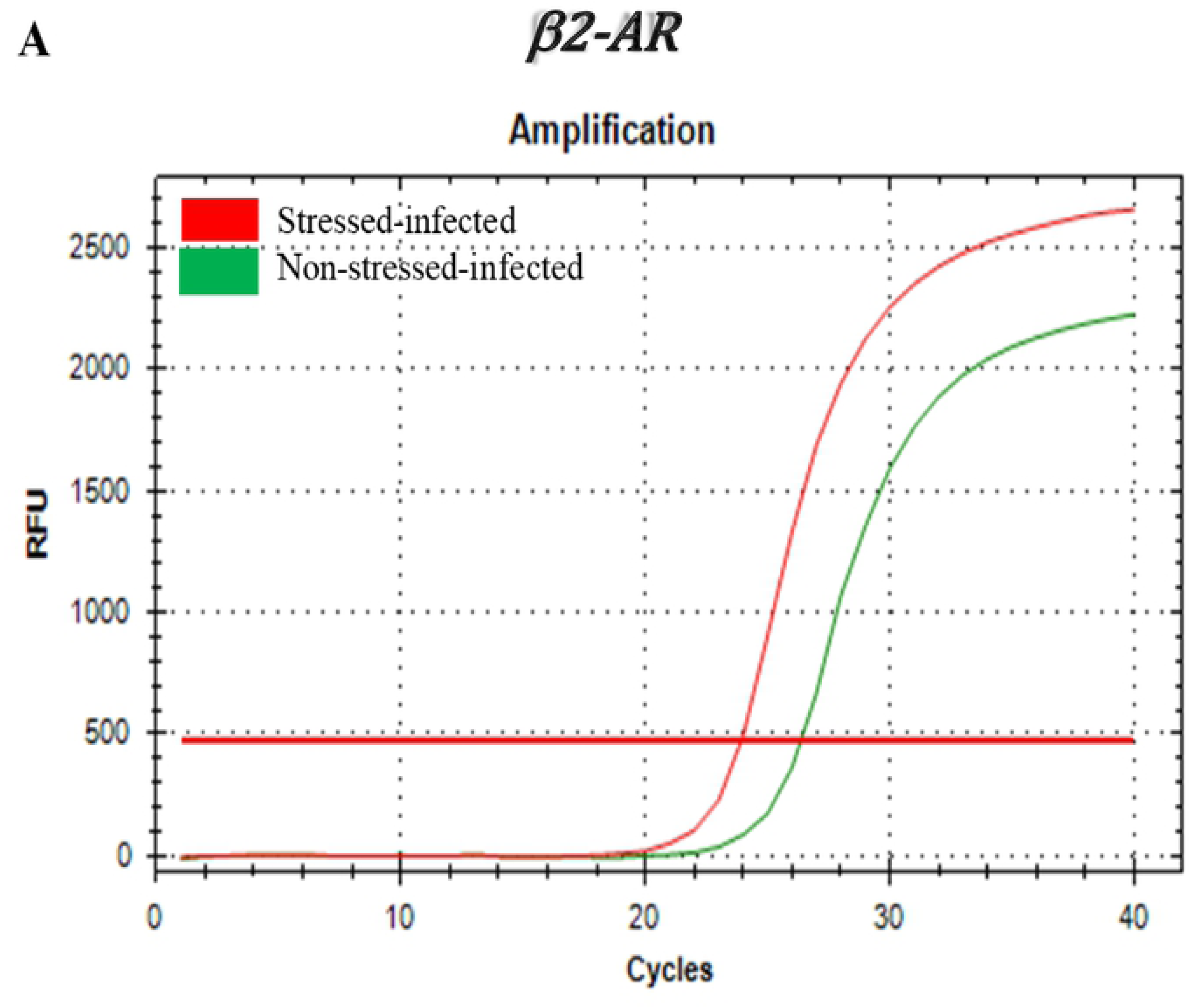
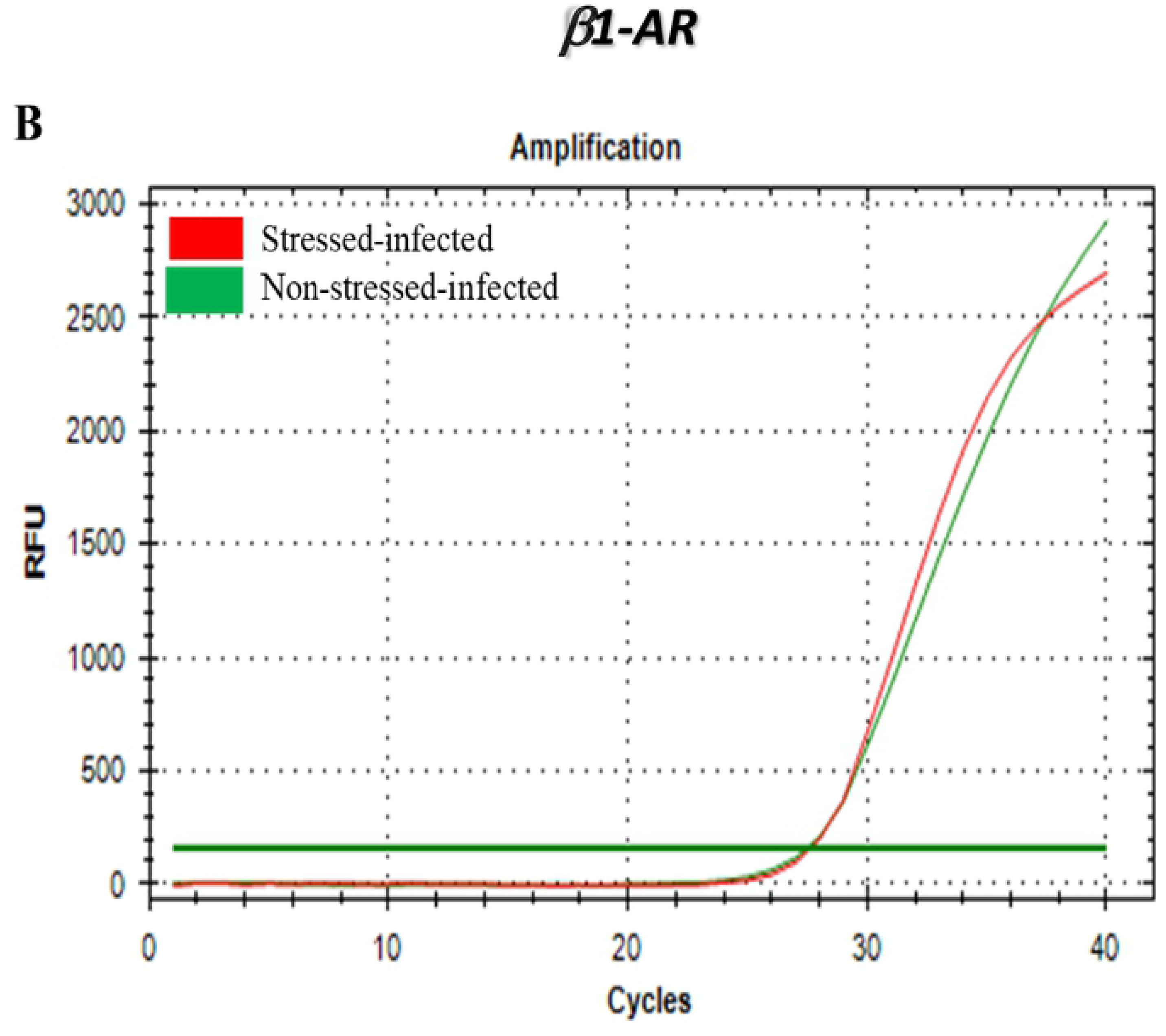
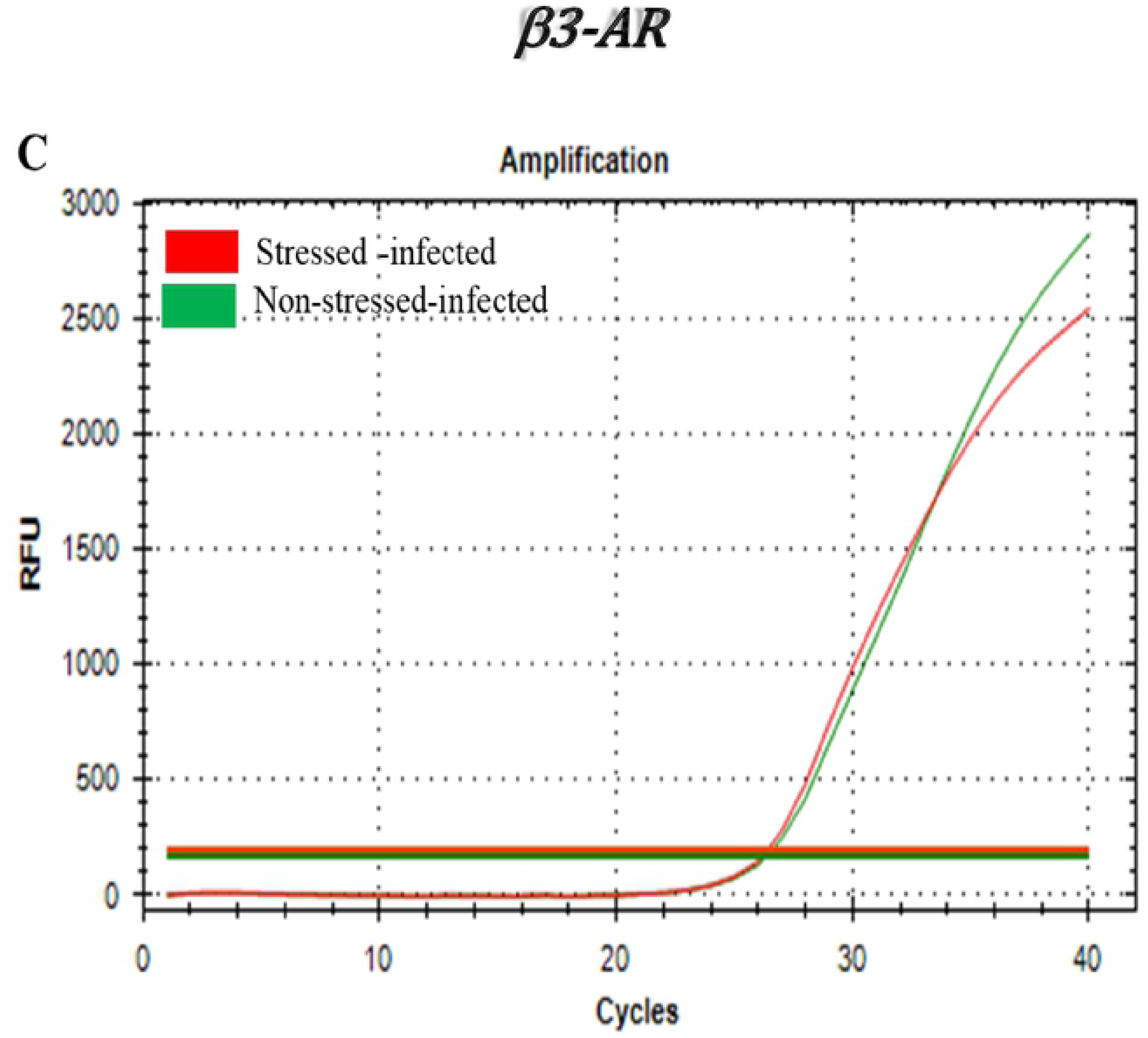
Gene expression profiles of beta-adrenergic receptor (β-AR) subsets in T cells isolated from the genital tract of stressed and non-stressed mice during *Chlamydia muridarum* genital infection. Amplification cures along with fold-changes of mRNA levels of β2-AR subsets normalized to the housekeeping gene, GADPH are shown. (**A**) β2-AR, (**B**) β1 –AR, (**C**) β3-AR. Fold-changes of mRNA levels of β1 –AR and β3-AR showed no significant difference between stressed and non-stressed group (data not shown). ***Abbs***: **St** = stressed, **Non-St** = none-stressed. Data shown are a representative of two or more independent experiments ran on different dates.

### Cold-induced stress leads to increased gene expression of transcription factor of GATA-3 in CD4+ T cells of genital tract during *Chlamydia muridarum* infection

Our laboratory showed previously that cold-induced stress treatment to mice results in differential gene expression and secretion of cytokines during *C. muridarum* genital infection. However, little is known about the effect cold-induced stress on the differentiation of the two major T helper (Th) Th1 and Th2 subsets. This study was to determine the gene expression profiles of transcription factors (T-bet and GATA-3) in all populations of CD4+ T cells isolated from the spleen and genital tract of stressed and non-stressed mice with or without *C. muridarum* genital infection. We hypothesized that cold-induced stress alters the gene expression patterns of T-bet and GATA-3 that determine the production of Th1 and Th2 type cytokines. As shown in **Figure 3A**, a marked increase of mRNA level of GATA-3 in CD4+T cells isolated from the genital tract of stressed mice was obtained and compared to that of non-stressed mice. Furthermore, a significant increase in gene expression of GATA-3 in CD4+ T cells isolated from the uterus and cervix of stressed mice is shown compared to that of non-stressed mice (**Figure 3C**). In contrast, T-bet mRNA expression was not elevated in either the whole genital tract or specific regions of the genital tract of both stressed and non-stressed mice (**Figure 3D, 3E**). Furthermore, T bet gene expression was down-regulated in oviduct CD4+ T cells of stressed mice compared to that of non-stressed mice (**Figure 3F**). DNA intensity of GATA-3 PCR end-product was significantly higher in stressed mice than non-stressed mice (**Figure 3G**). These results suggest that transcription factor GATA-3 may play a big role in the switching of Th1 to Th2, arming the immune system during stressful conditions of chlamydia genital infection.

**Figure 3:**
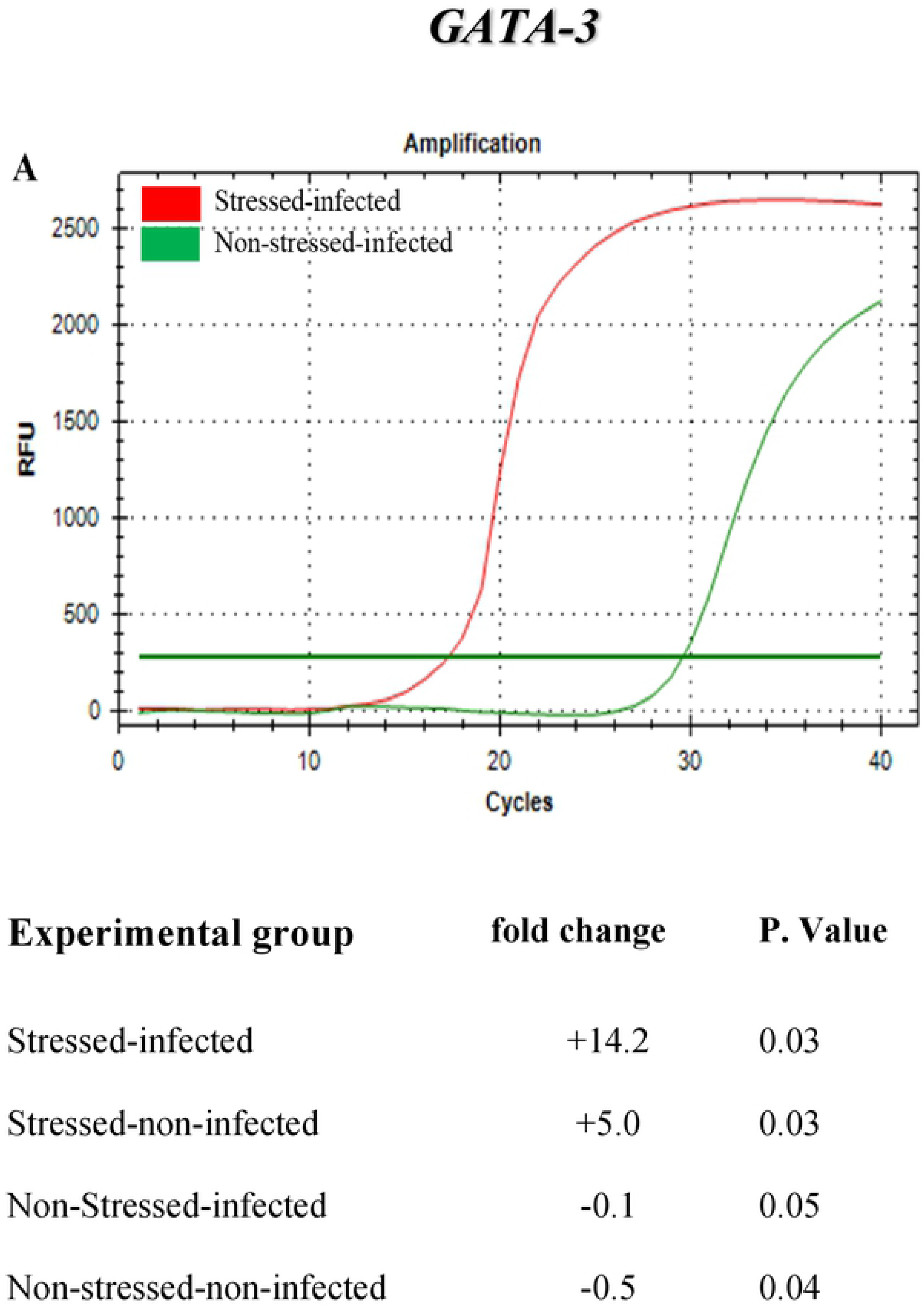
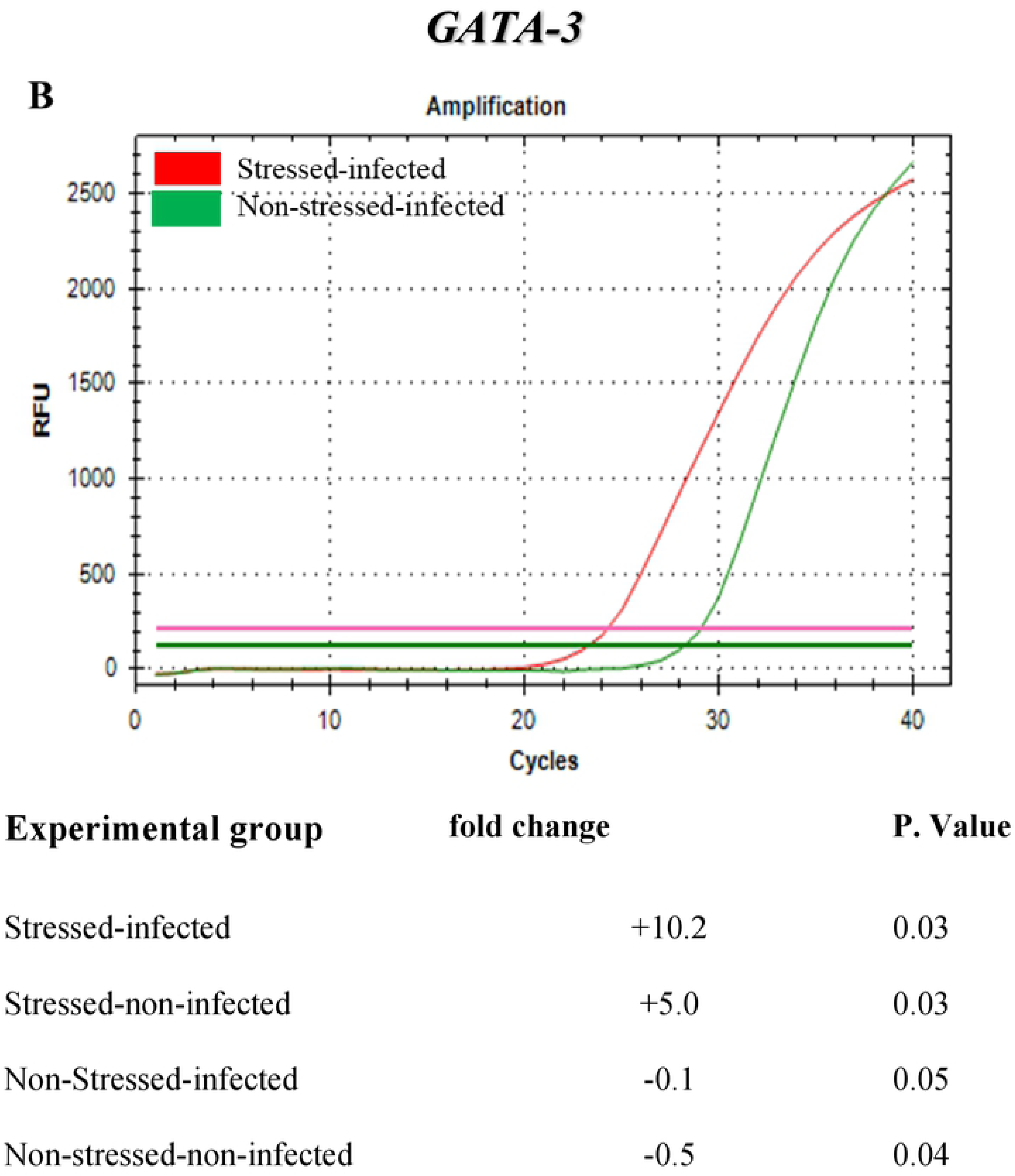
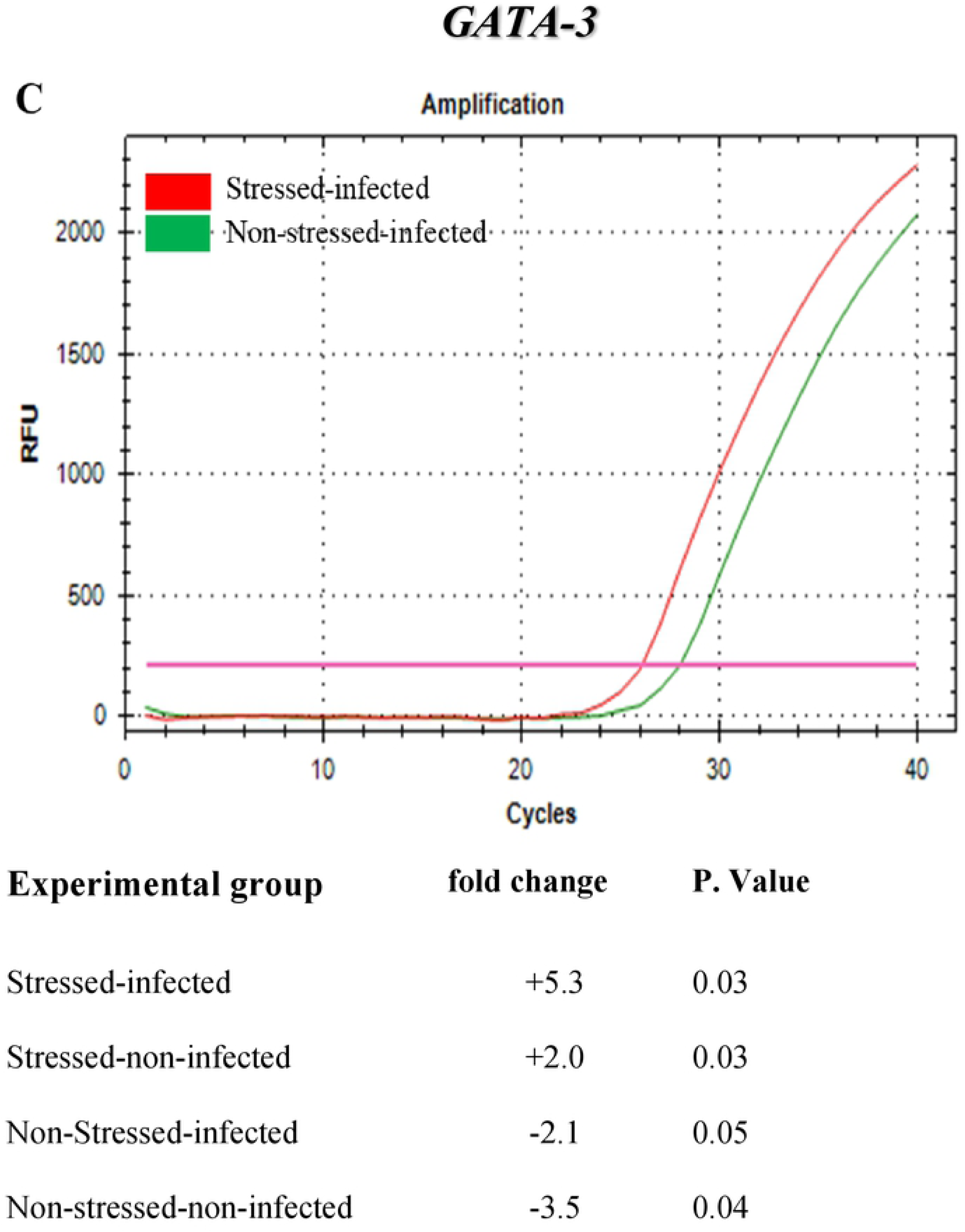
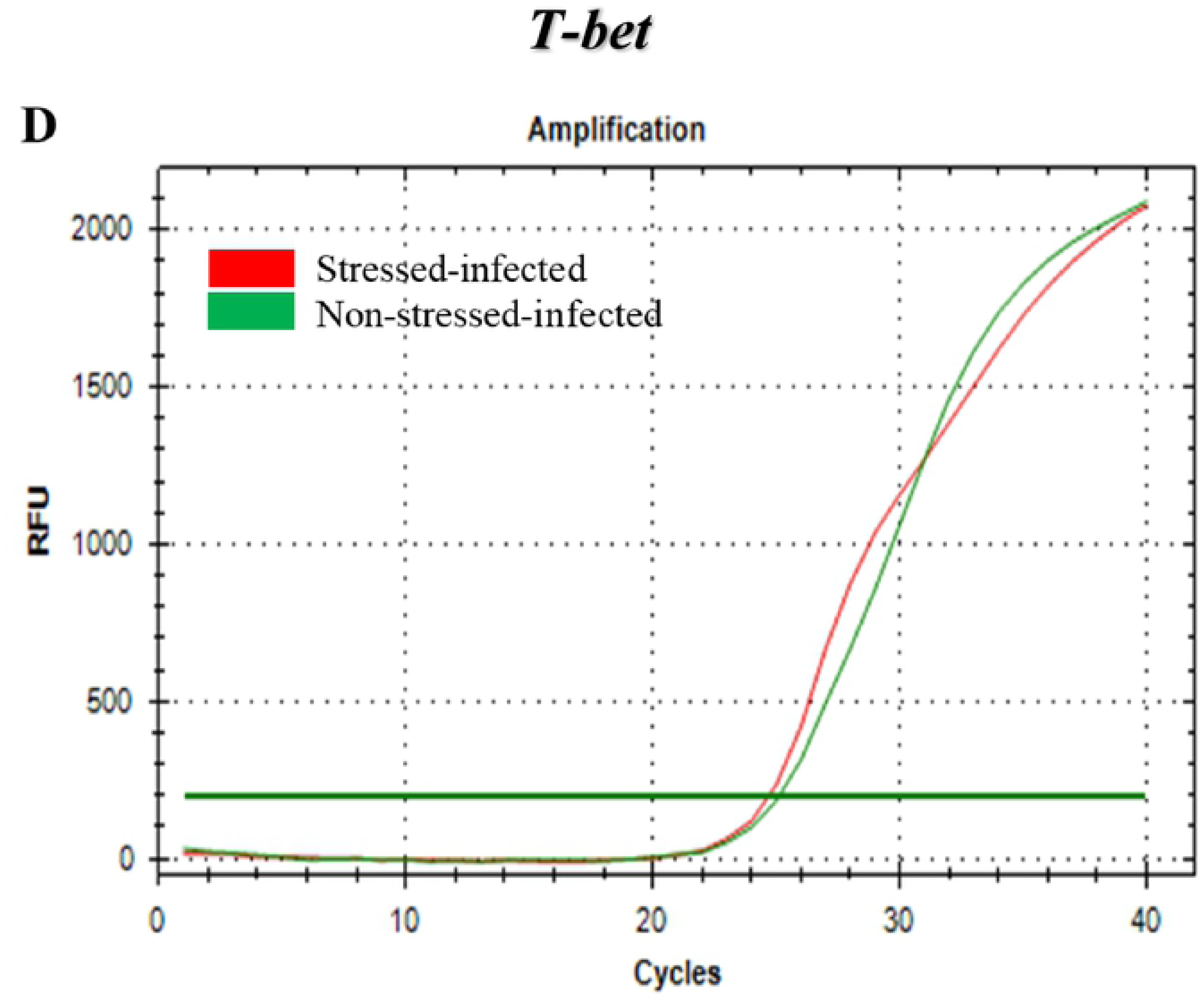
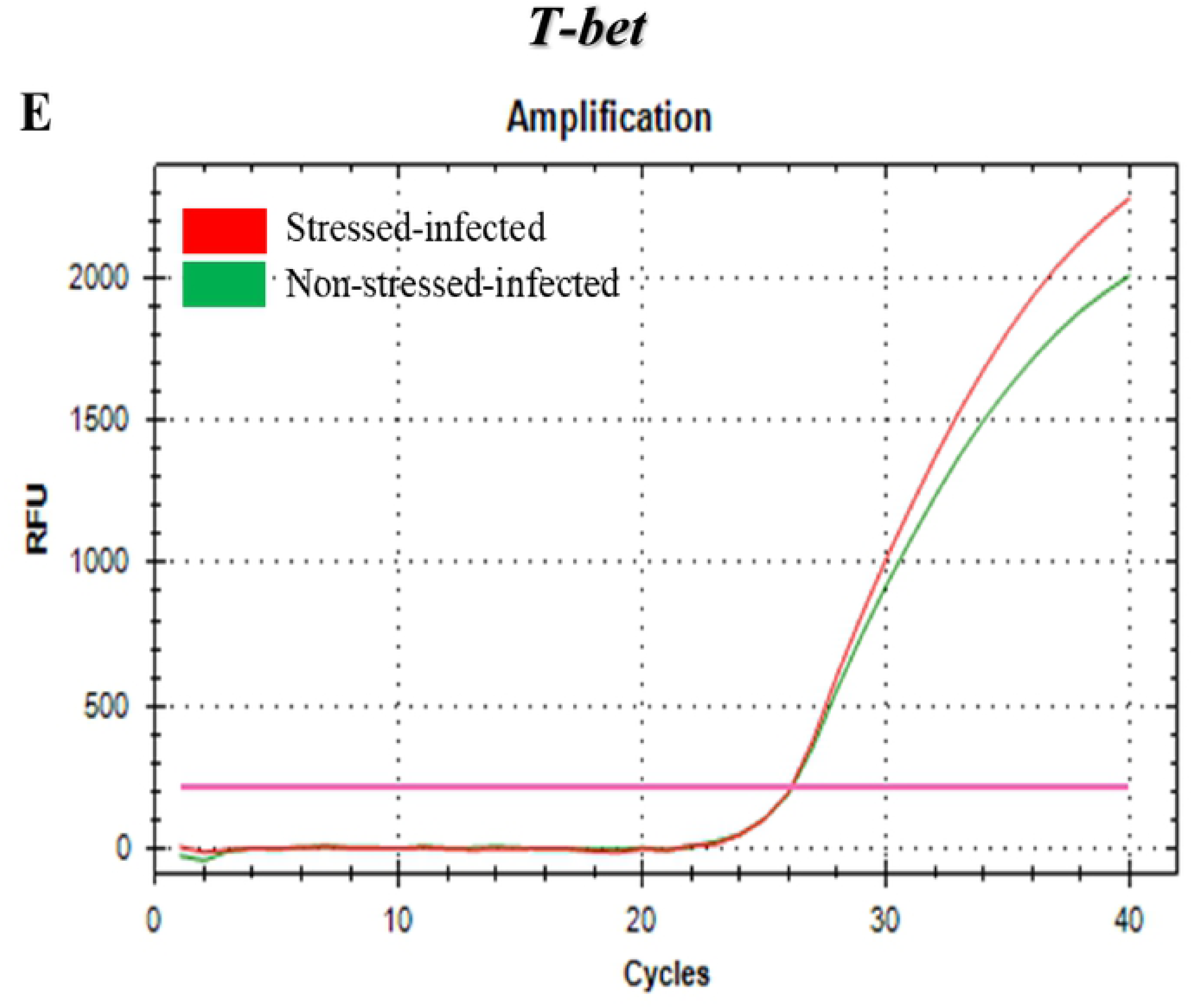
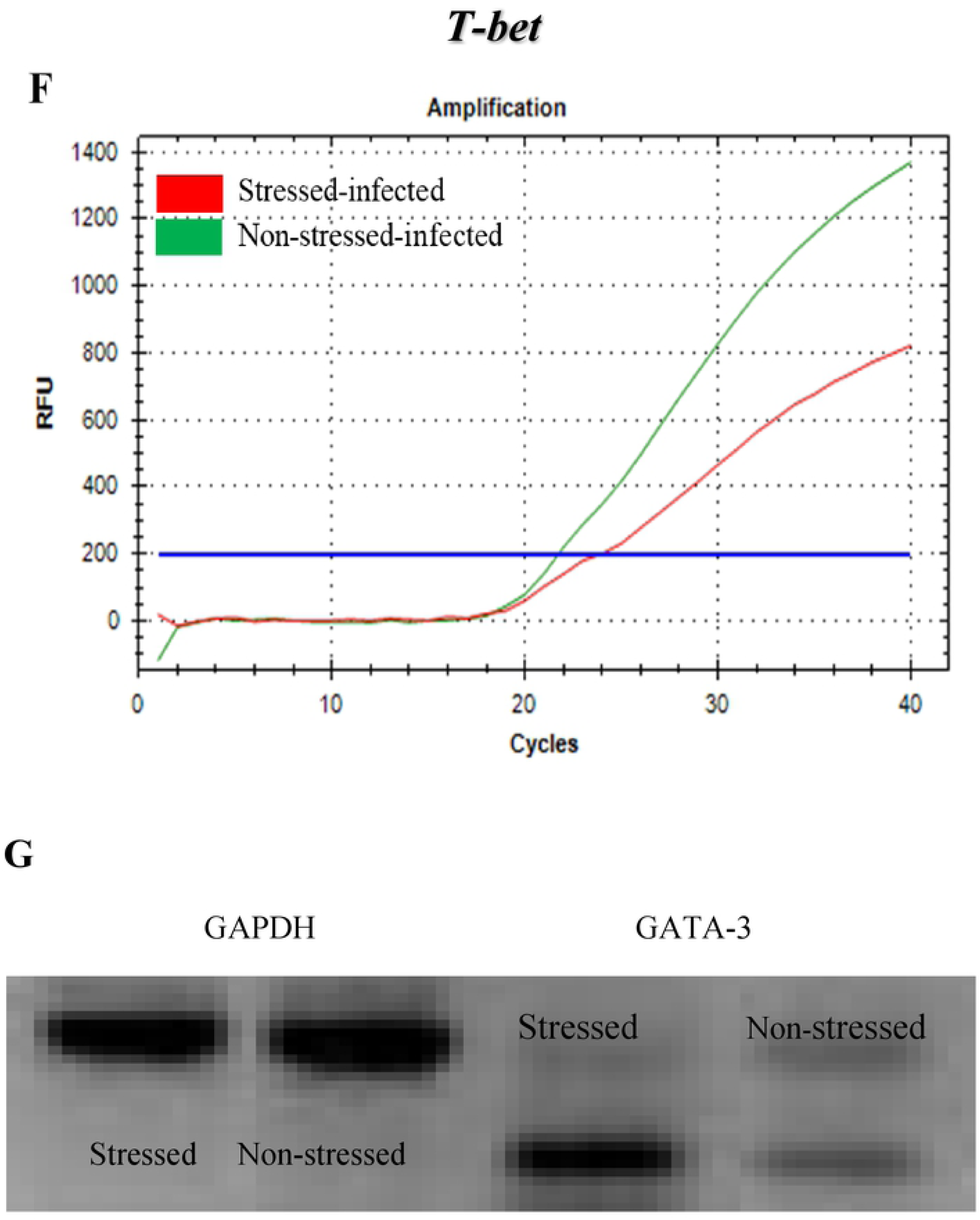
Gene expression profiles of GATA-3 and T-bet in T cells isolated from the whole or the parts of the genital tract of stressed and non-stressed mice during *Chlamydia muridarum* genital infection. Amplification cures along with fold-changes of mRNA levels of transcription factors of T cells isolated (**A**) whole genital tract, (**B**) uterus horns, (**C**) oviduct, (**D**) T-bet in T cells of whole genital tract, (**E**) uterus horns (**F** oviduct), and (**G**) gel electrophoresis of GATA-3 are shown. Fold-changes of T-bet showed no significant difference between stressed and non-stressed group (data not shown). ***Abbs***: **St** = stressed, **Non-St** = none-stressed. Shown data are representatives of two or more independent experiments.

### Cold-induced stress leads to increased gene expression of transcription factor GATA-3 in lysate of whole and regions of the genital tract of stress mice during *Chlamydia muridarum* infection

The purpose of this study was to determine if there is a differential gene expression in the different regions of the genital tract during *Chlamydia muridarum* genital infection. The mRNA profiles of GATA-3 and T-bet in the lysates of whole genital tract, cervix, uterine horns, and oviducts of stressed and non-stressed mice with/without *C. muridarum* were assessed. Our finding is that, after 48 hours of infection cold-induced stress results in up-regulated expression of GATA-3 in oviduct (**Figure 4A**), uterine horns (**Figure 4B**), and cervix (**Figure 4C**). In contrast, cold-induced stress resulted in down-regulated gene expression of T-bet in the cervix (**Figure 4D**) and uterine horns (**Figure 4E**) of stressed mice compared to that of non-stressed mice. However, the mRNA level of T-bet in the cervix was undetectable. Gene expression of transcription factor of GATA-3 was up-regulated in the different regions of the genital tract of stressed mice during *C. muridarum* infection, while gene expression of T-bet was reduced compared to non-stressed mice. Our results indicate a positive correlation in gene expression patterns of transcription factors in CD4+ T cells and genital tract lysates of stressed and non-stressed mice demonstrating differences in the magnitude of gene expression of GATA-3 and T-bet in differing regions of the genital tract, reflecting the dynamic nature of the transcription factors during stress infection.

**Figure 4:**
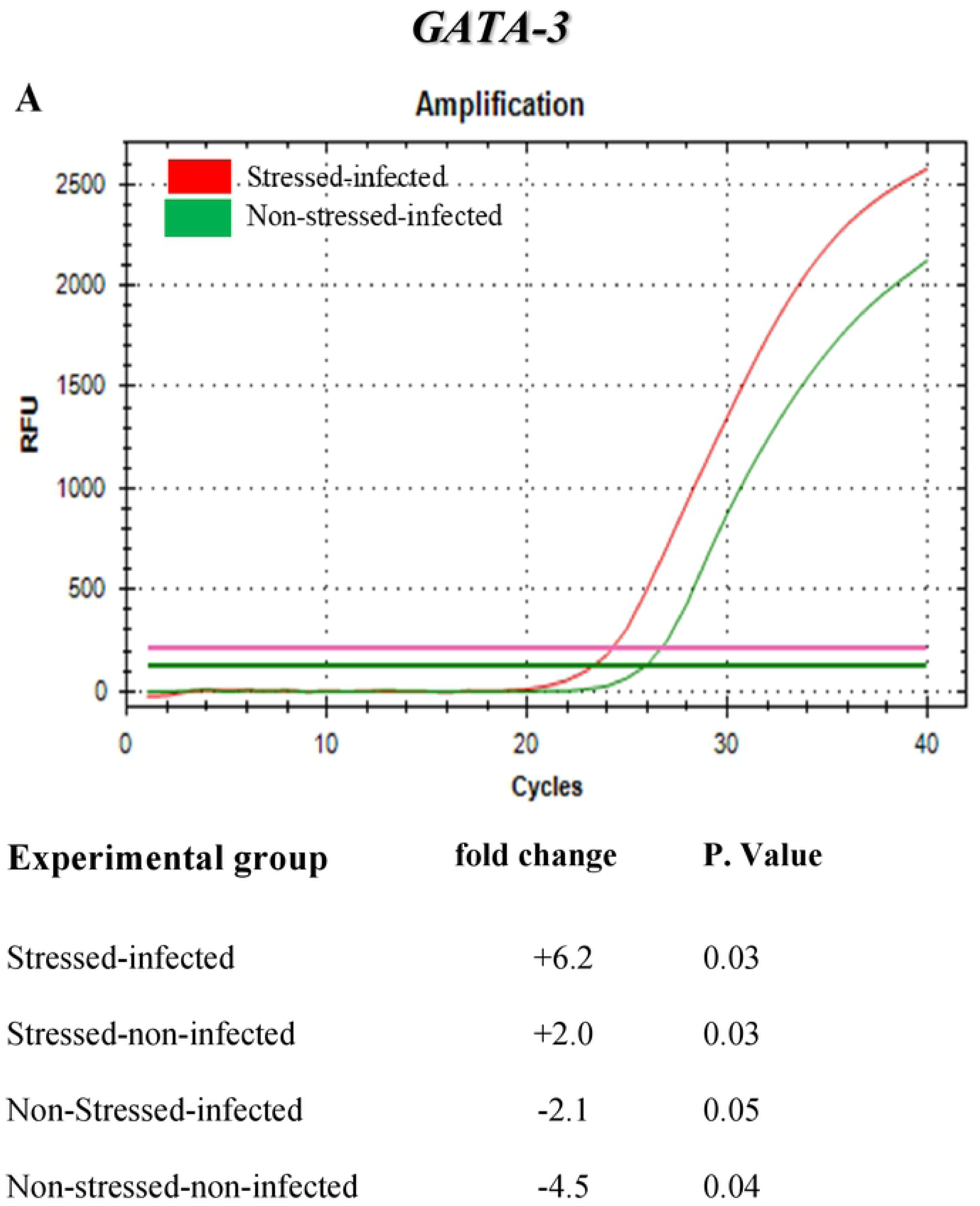
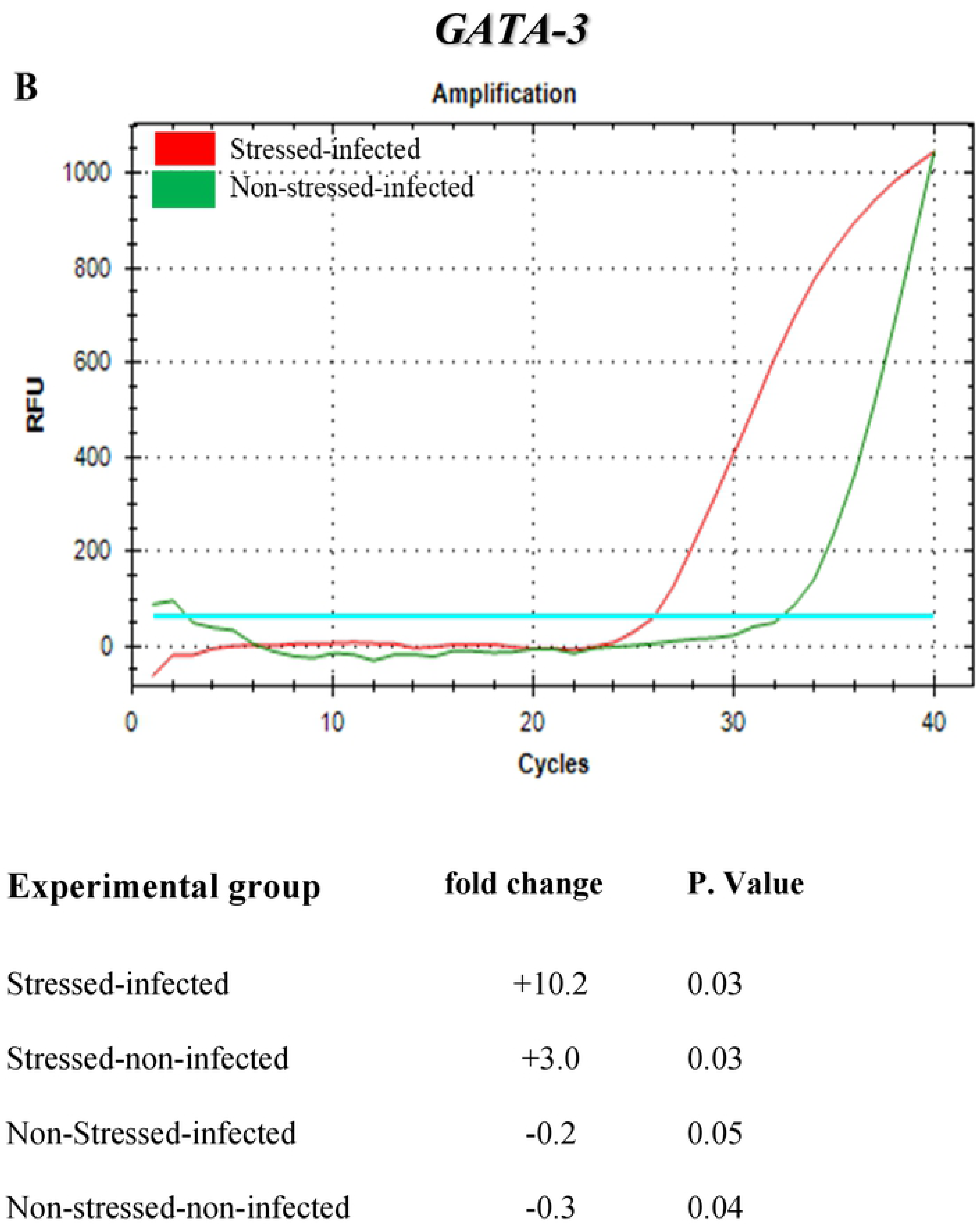
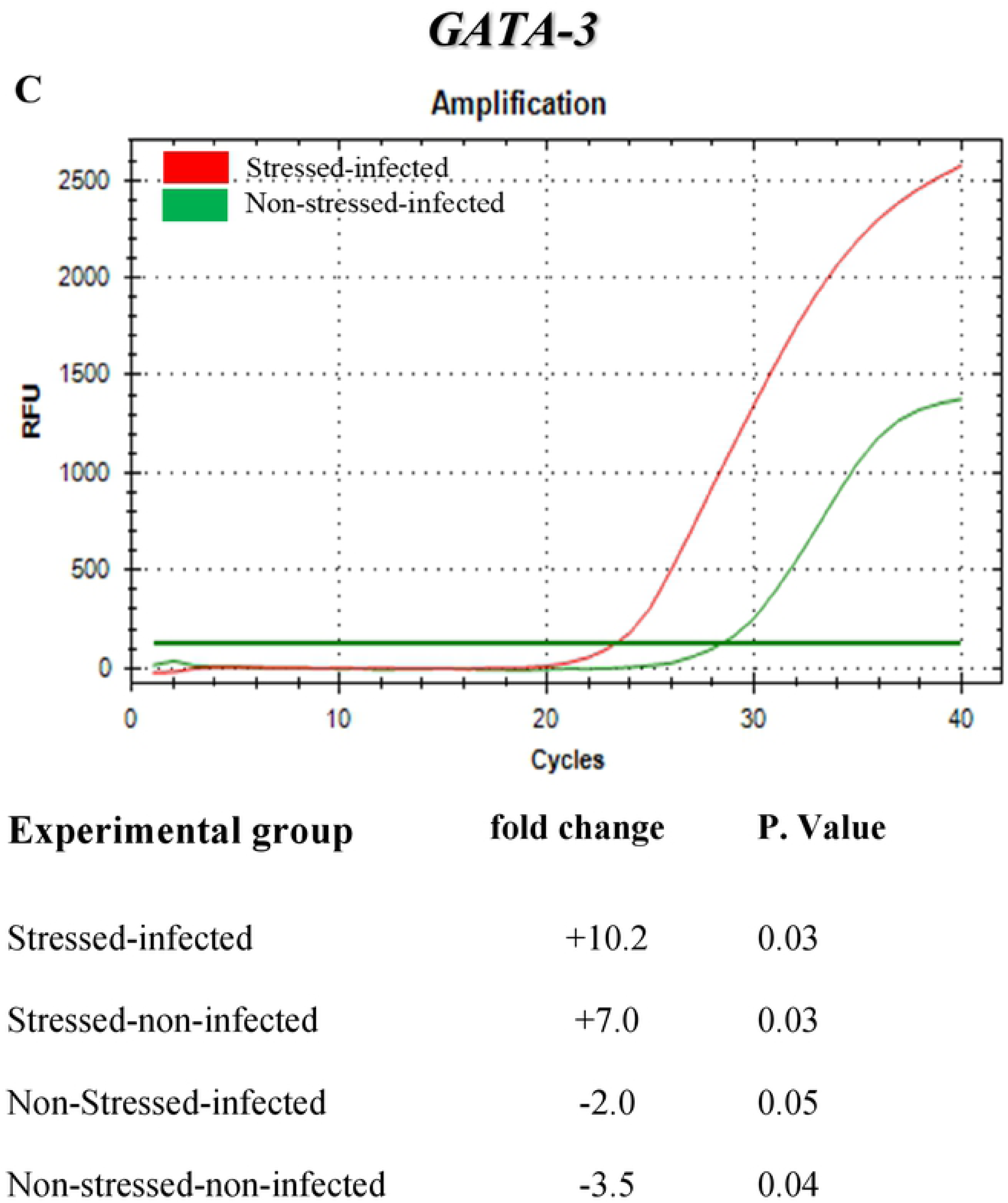
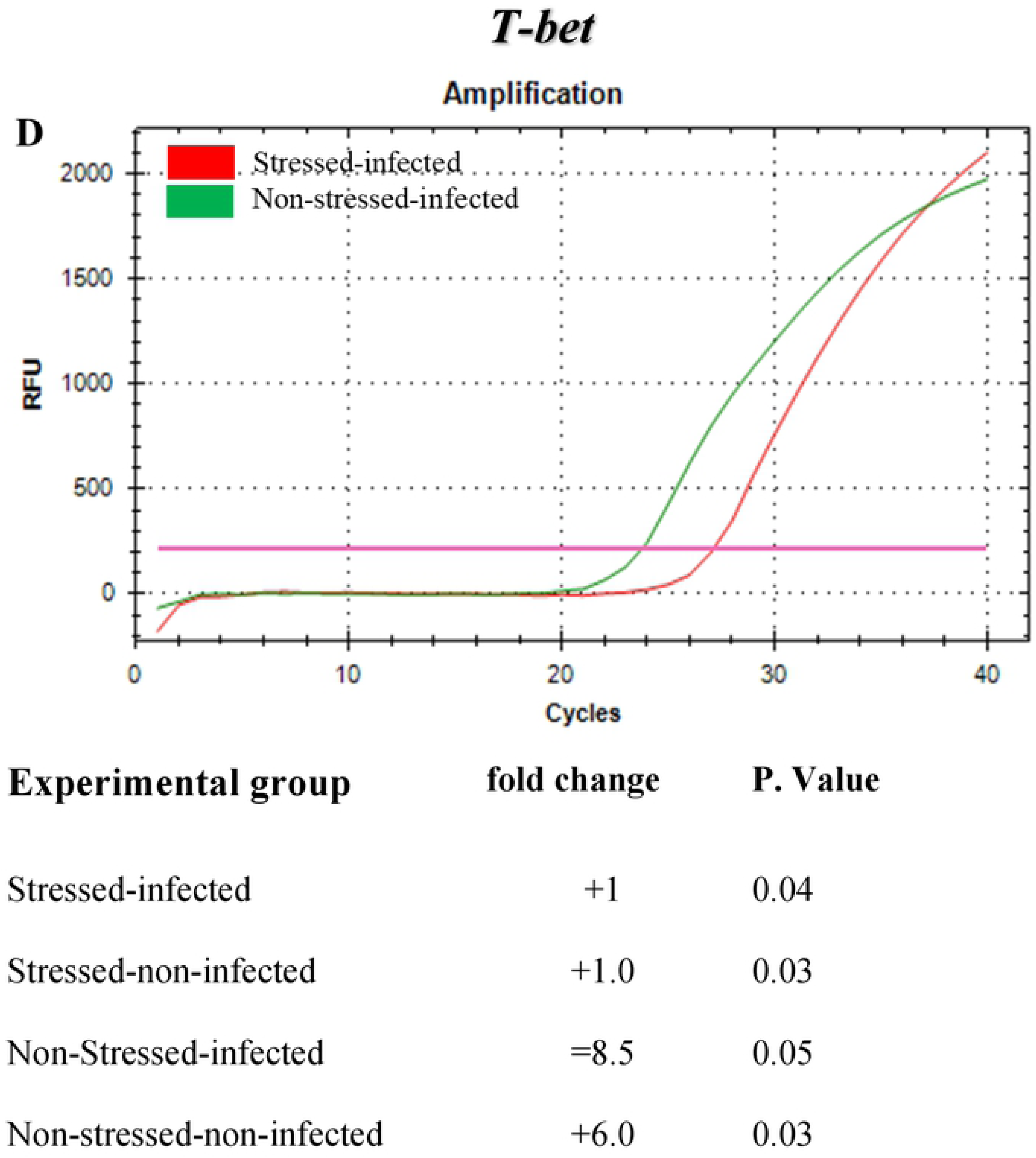
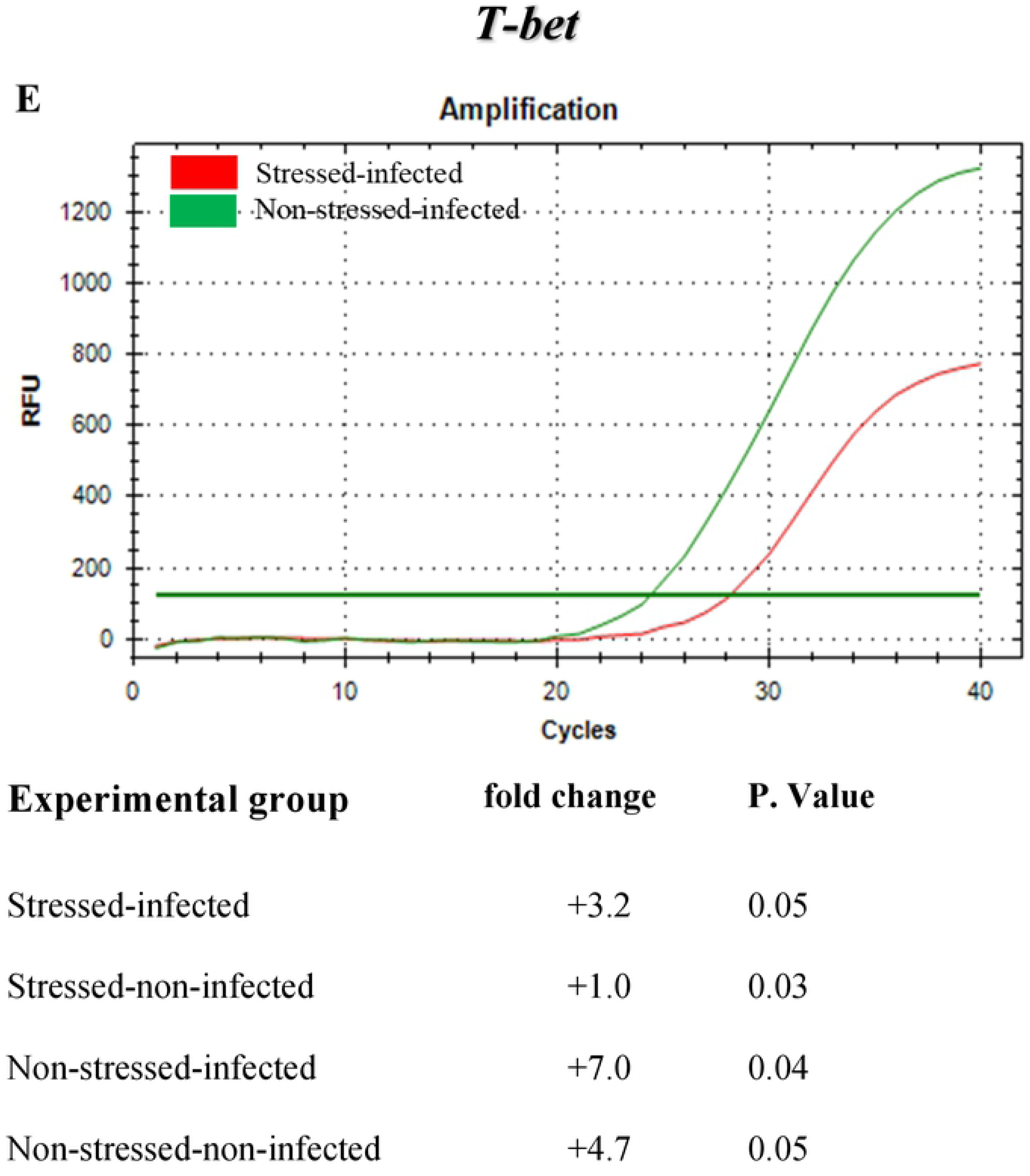
Amplification cures along with fold-changes of mRNA levels of transcription factors GATA-3 and T-bet in lysates of the whole or parts of genital tract of stressed or non-stressed mice during *Chlamydia muridarum* genital infection. (**A**) GATA-3 in oviduct (**B**) uterus horns, (**C**) cervix, (**D**) T-bet in oviduct (**E**) uterus horns. No gene expression was detected T-bet in the cervix of stressed or non-stress mice. Data shown are a representative of two or more independent experiments ran on different dates.

### Cold-induced stress alters gene expression patterns of transcription factors and signature cytokines in mouse splenic naïve CD4+ T cells during *Chlamydia muridarum* genital infection

Our aim was to further evaluate the gene expression of transcription factors (GATA3 and T-bet) and IL-4 and γ-IFN in splenic naïve CD4+ T cells isolated from stressed and non-stressed mice with/without *C. muridarum* genital infection. As shown **Figure 5A**, the lowest Ct value and highest fold-change (p < 0.05) of GATA-3 gene expression was observed in stressed/infected mice compared to the other treatment groups. In contrast, the highest Ct value and negative fold-change of T-bet gene expression (p <0.05) was obtained in stressed/infected mice (**Figure 5B**). Similarly, the Ct value and the highest fold-changes of IL-4 was observed in stressed/infected mice (**Figure 5C**), whereas the highest Ct value and negative value in fold-changes of IFN-γ was observed in the stressed group (**Figure 5D**).

**Figure 5:**
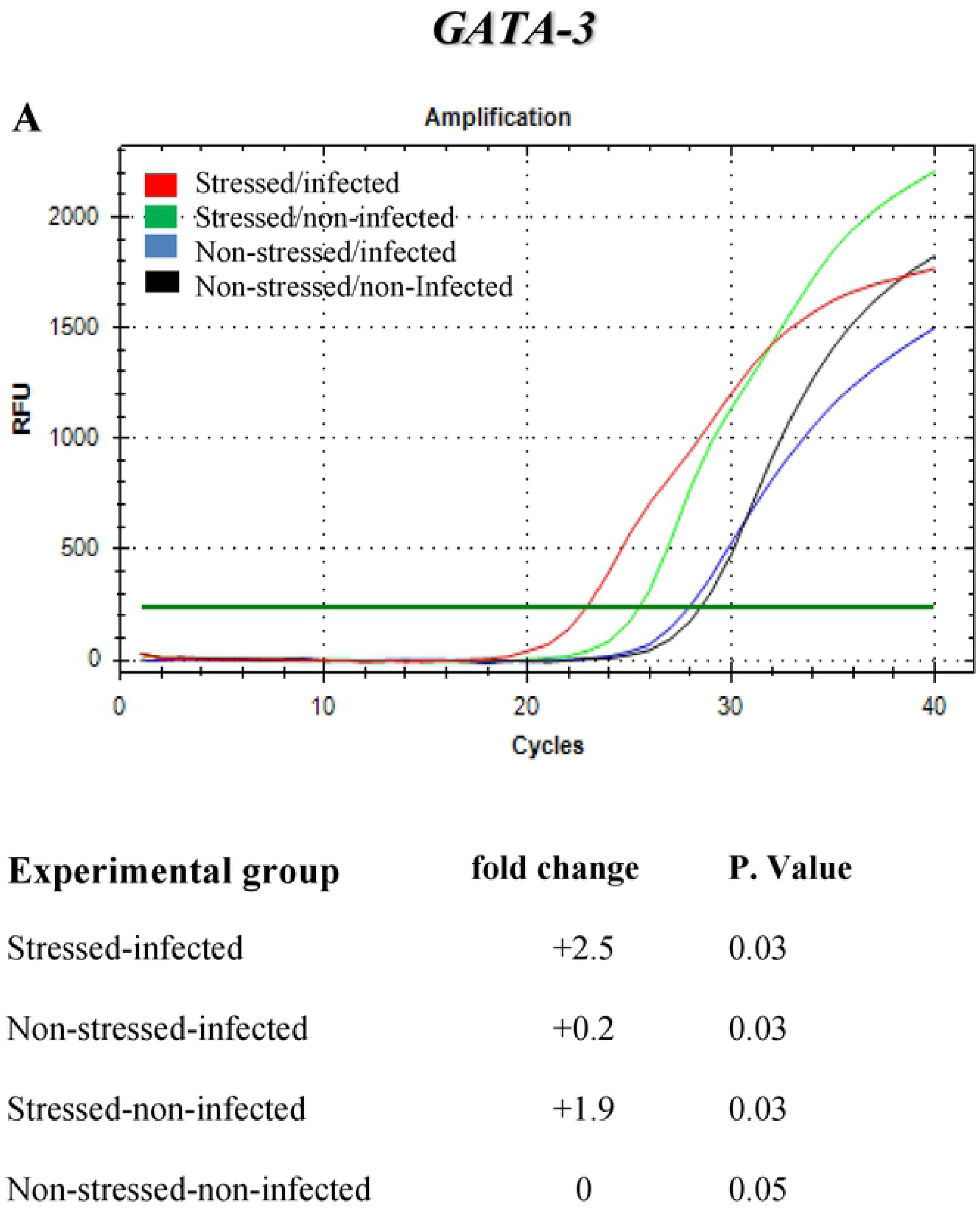
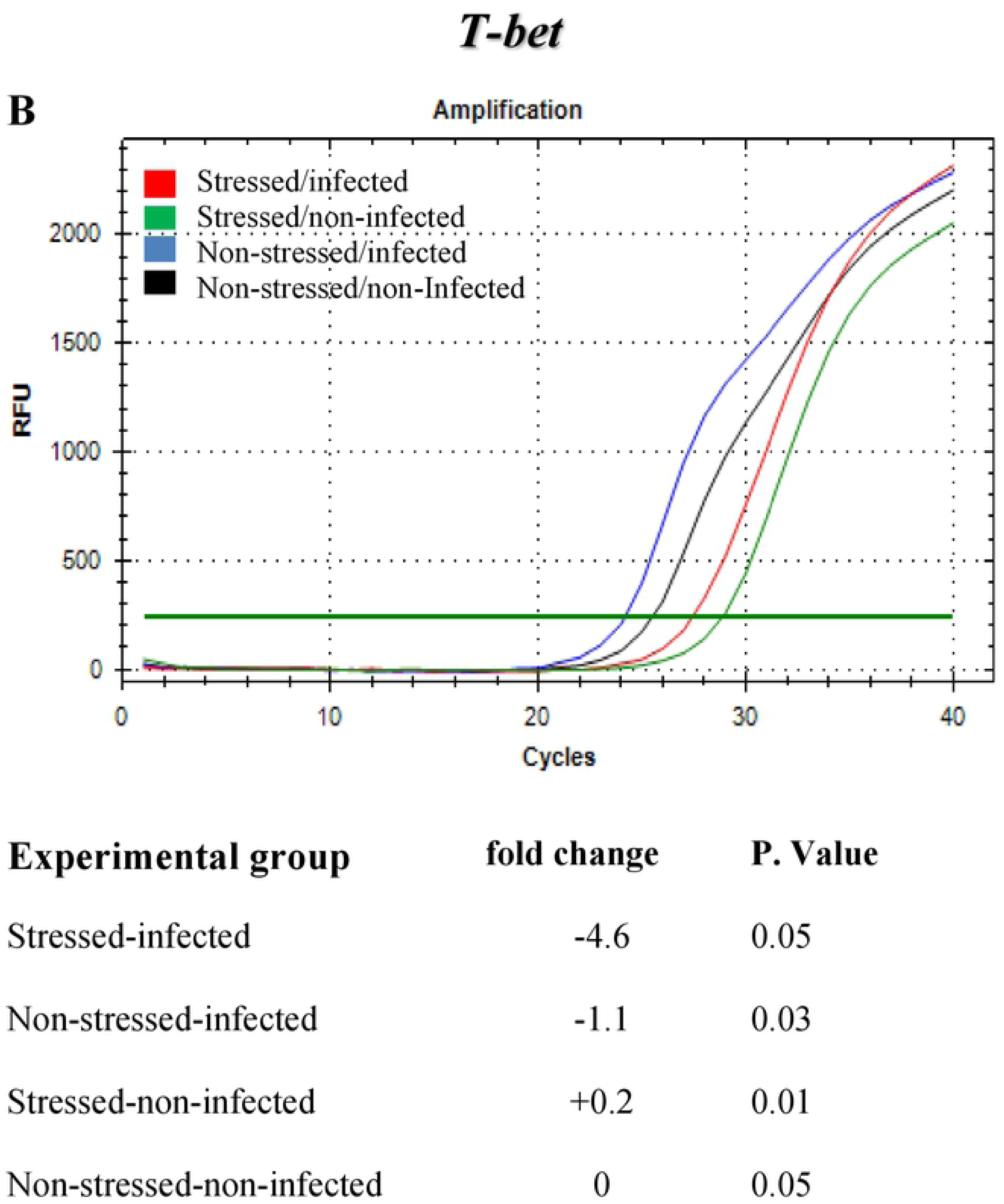
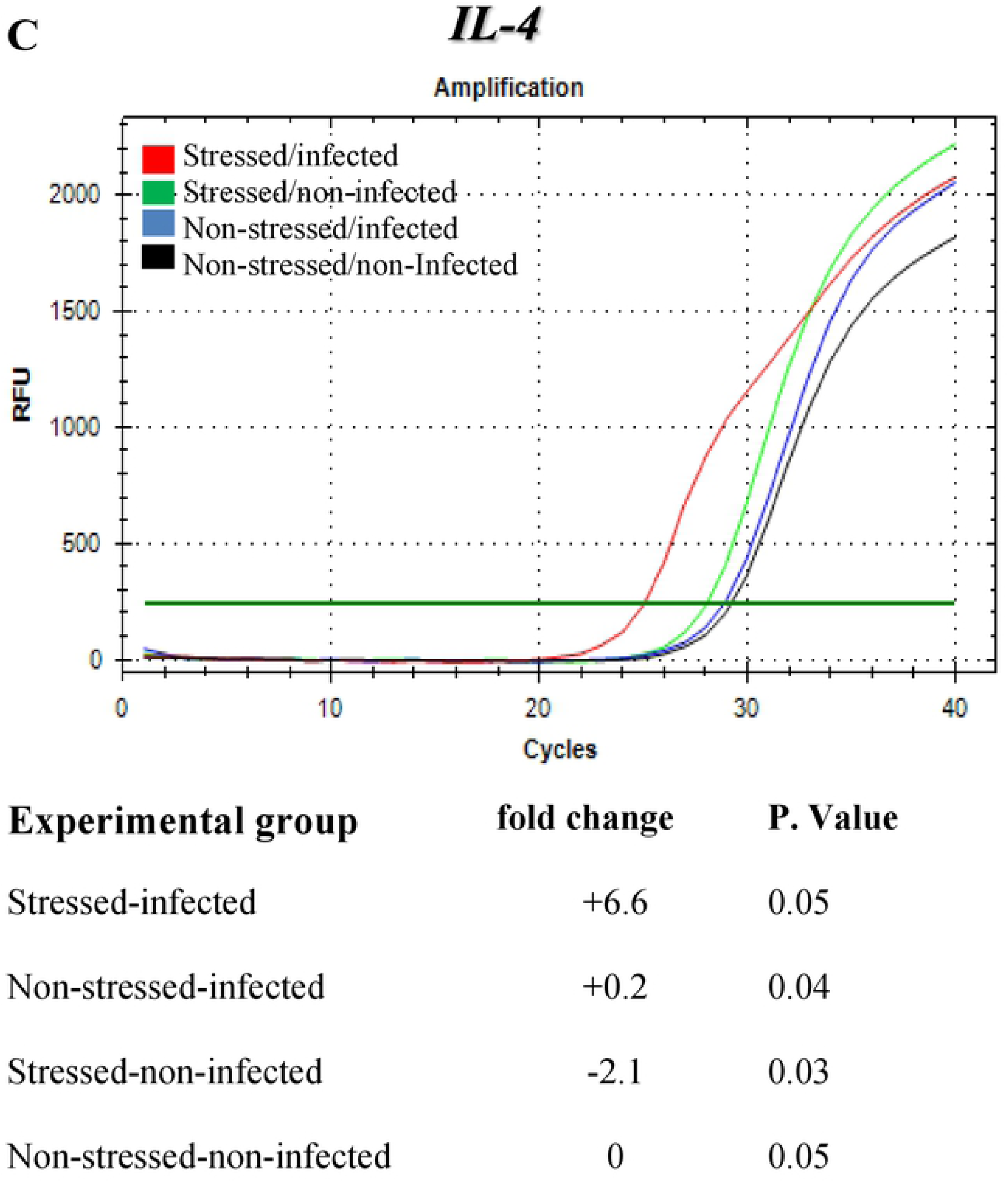
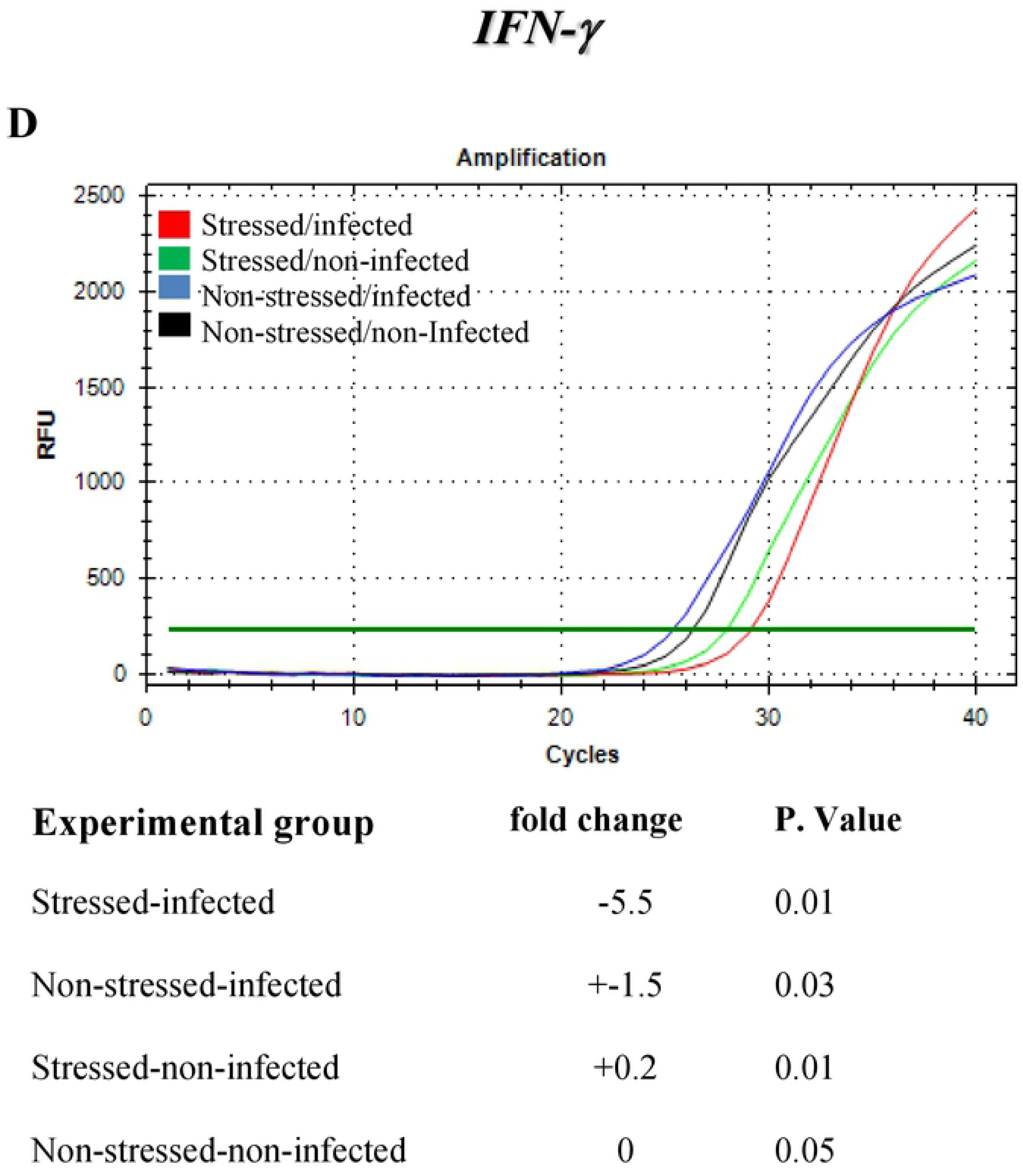
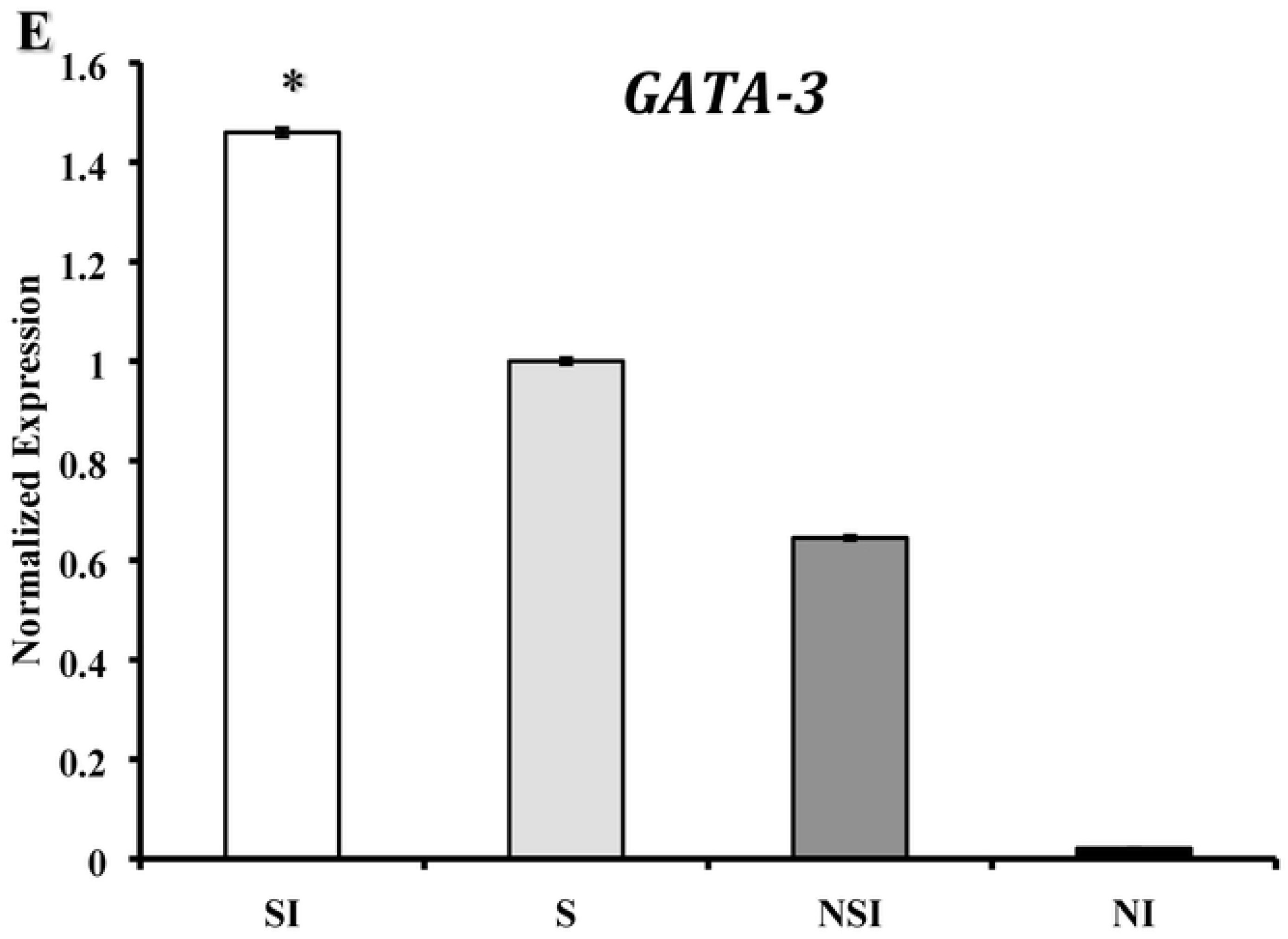
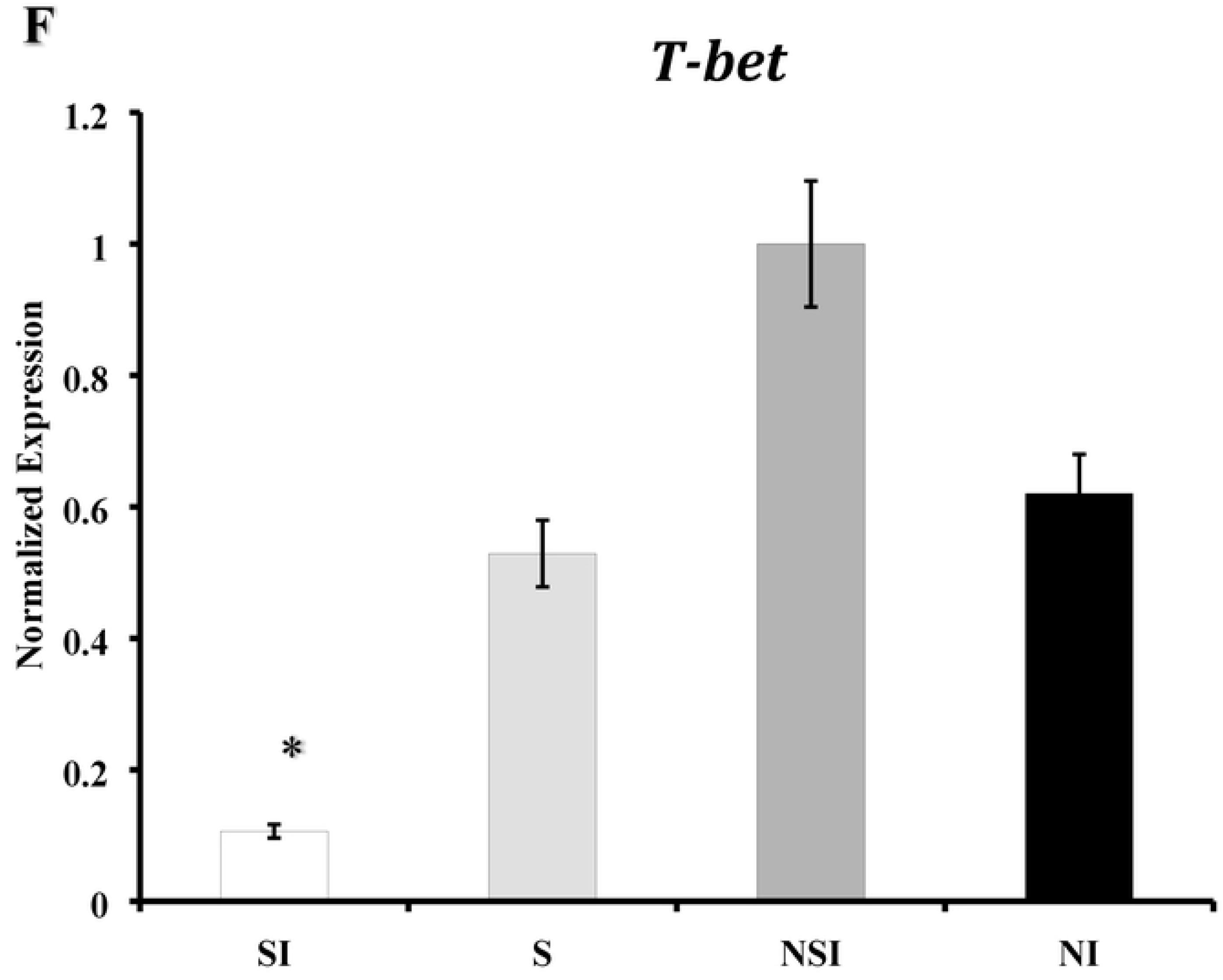
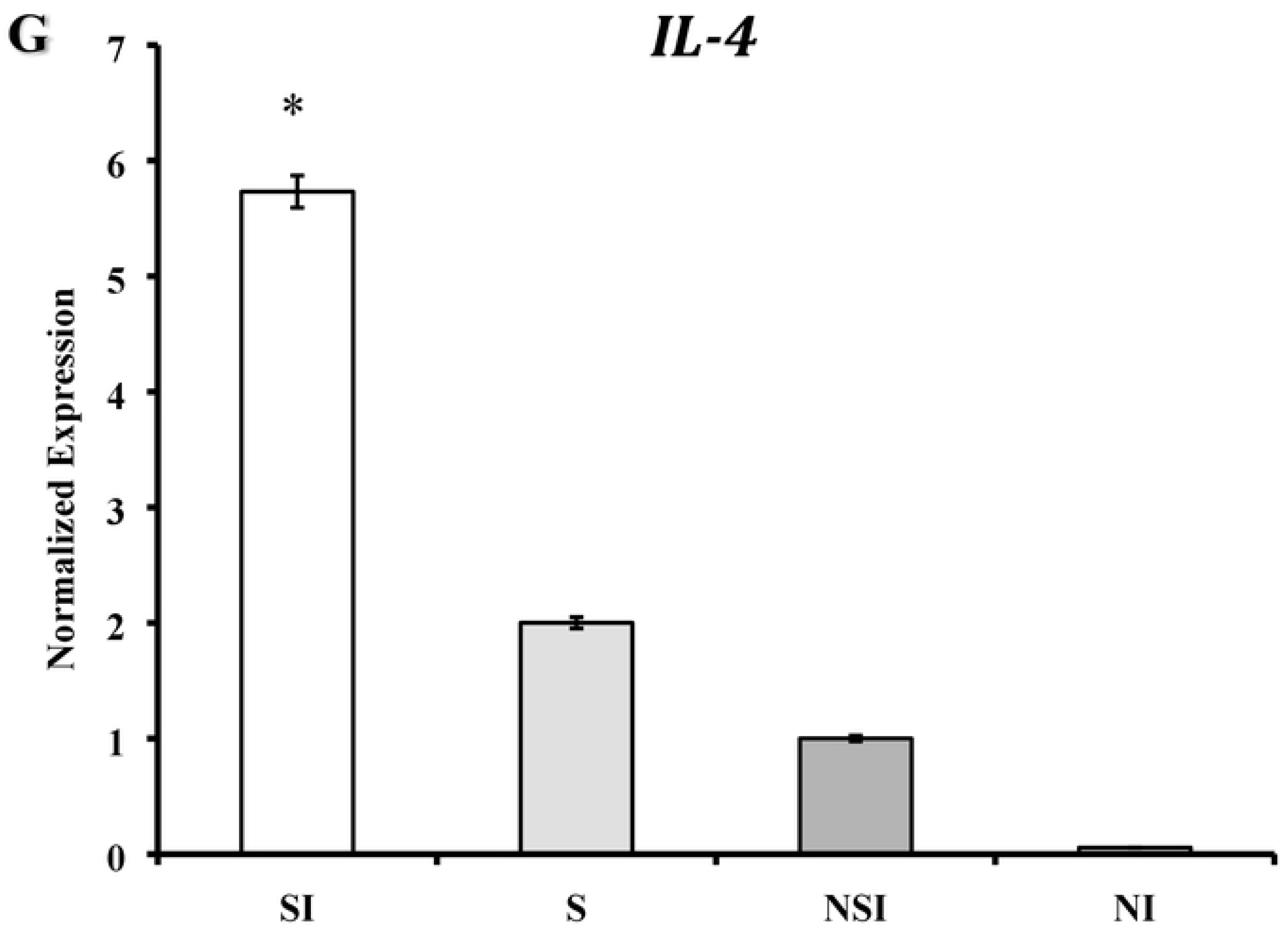
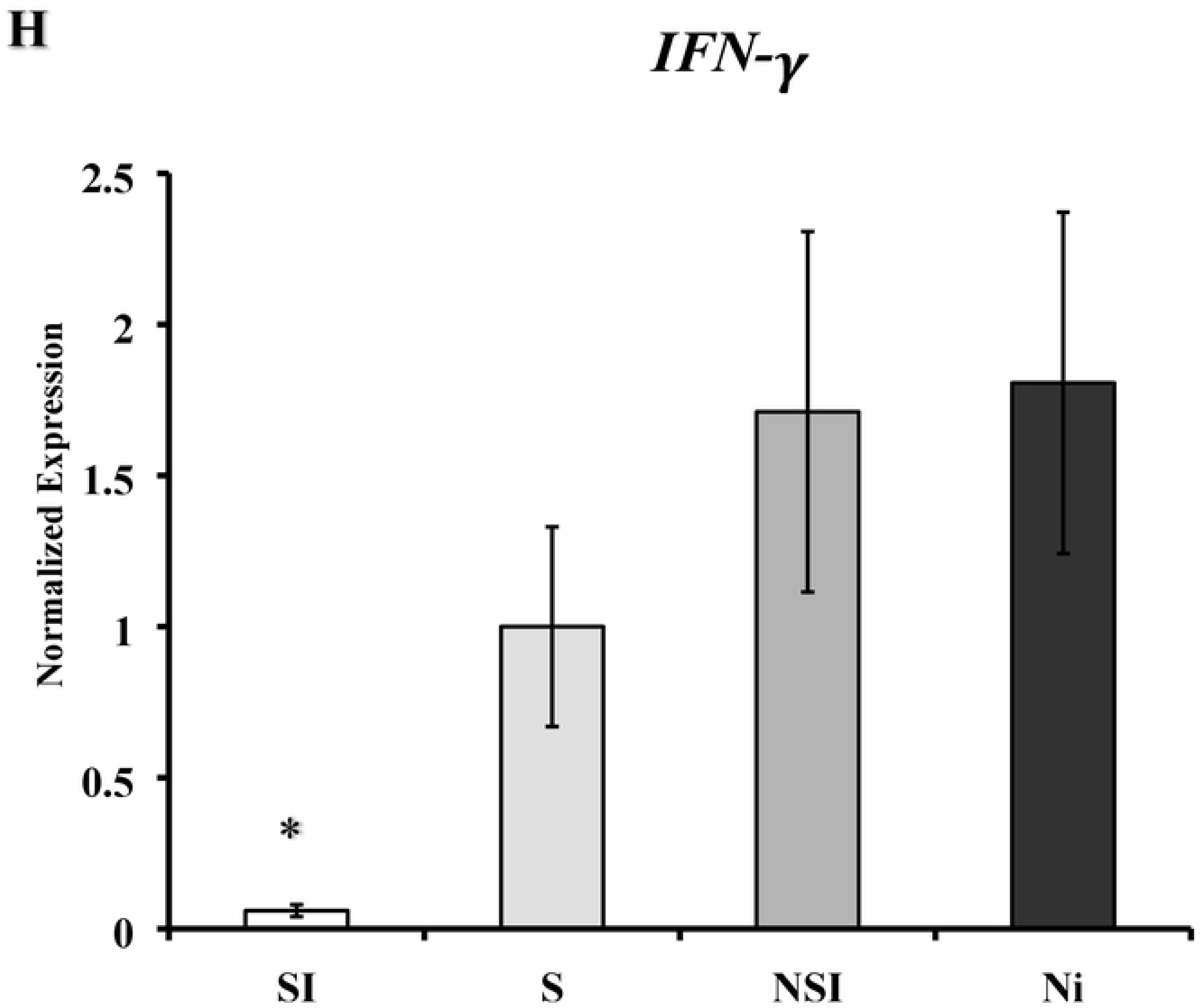
Gene expression profiles of (**A**) GATA-3, (**B**) T-bet, (**C**) Interleukin-4 (IL-4, and (**D**) gamma-interferon (IFN-γ) in naïve CD4+ T cells isolated from spleen cells. Expression fold changes of (**E**), GATA-3, T-bet (**F**), (**G**) IL-4 and (**H**) IFN-γ in splenic T cells normalized to the internal control gene (GAPDH) and relative to the non-stress control are shown. Data shown are a representative or a mean +/_ SEM of two or more independent experiments ran on different dates. *Denotes significant statistical differences between treatment groups at the level of (p < 0.05).

The normalized mRNA levels of GATA-3 and IL-4 in CD+ T cells displayed significantly increased expression in stressed and infected mice, while T-bet and IFN-γ in CD+ T cells displayed significantly decreased expression in stressed and infected mice. Our results show a direct correlation between the Ct values and fold-changes of the transcription factors and the selected cytokines. The gene expressions of GATA-3, T-bet, IL-4, and IFN-γ in naïve CD4+T cells isolated from stressed and non-stressed mice normalized to the internal control gene GAPDH are shown in (**Figure 5E, 5F, 5G**, and **5H**). The results show a positive correlation in gene expression patterns of GATA-3 and IL-4 or T-bet and IFN-γ in stressed and non-stressed mice.

### Persistence of stress effect on modulation of gene expression of transcription factors and secretion of Cytokines

To determine how long the effect of stress persists, we compared the gene expression of T cell transcription factors or secretion of signature cytokines by CD4+ T cells at two- or 11-days post infection. Glyceraldehyde 3-phosphate dehydrogenase (GAPDH) was used as an internal control to normalize target gene expression levels, which were determined using the 2-ΔΔCT method. As shown in **Figure 6A**, a marked increase in expression of mRNA encoding GATA-3 was demonstrated at day two but expression was significantly decreased at day 11 in stressed and infected mice compared to non-stressed and infected mice (P<0.05). As shown in **Figure 6B**, a marked decrease in expression of mRNA encoding T-bet demonstrated at day two remained constant through day 11 in stressed and infected mice compared to non-stressed and infected mice (p< 0.05). In contrast, normalized gene expression of IL-4 at day 2 was high and further increased at day 11 compared to the non-stressed and infected group (**Figure 6C**). Similarly, the T-bet gene expression profile of IFN-γ remained significantly low days 2 through 11 in stressed mice compared to the non-stressed mice (**Figure 6D**). The results presented here strongly suggest the cold-induced stress effect of enhanced IL-4 production and decreased IFN-γ persists for days. Moreover, our findings suggest that the upregulation of GATA-3 and IL-4 is associated with the induction of Th2 and suppression of Th1 CD4+ T cells in response to prolonged chronic stress and chlamydia genital infection.

**Figure 6.**
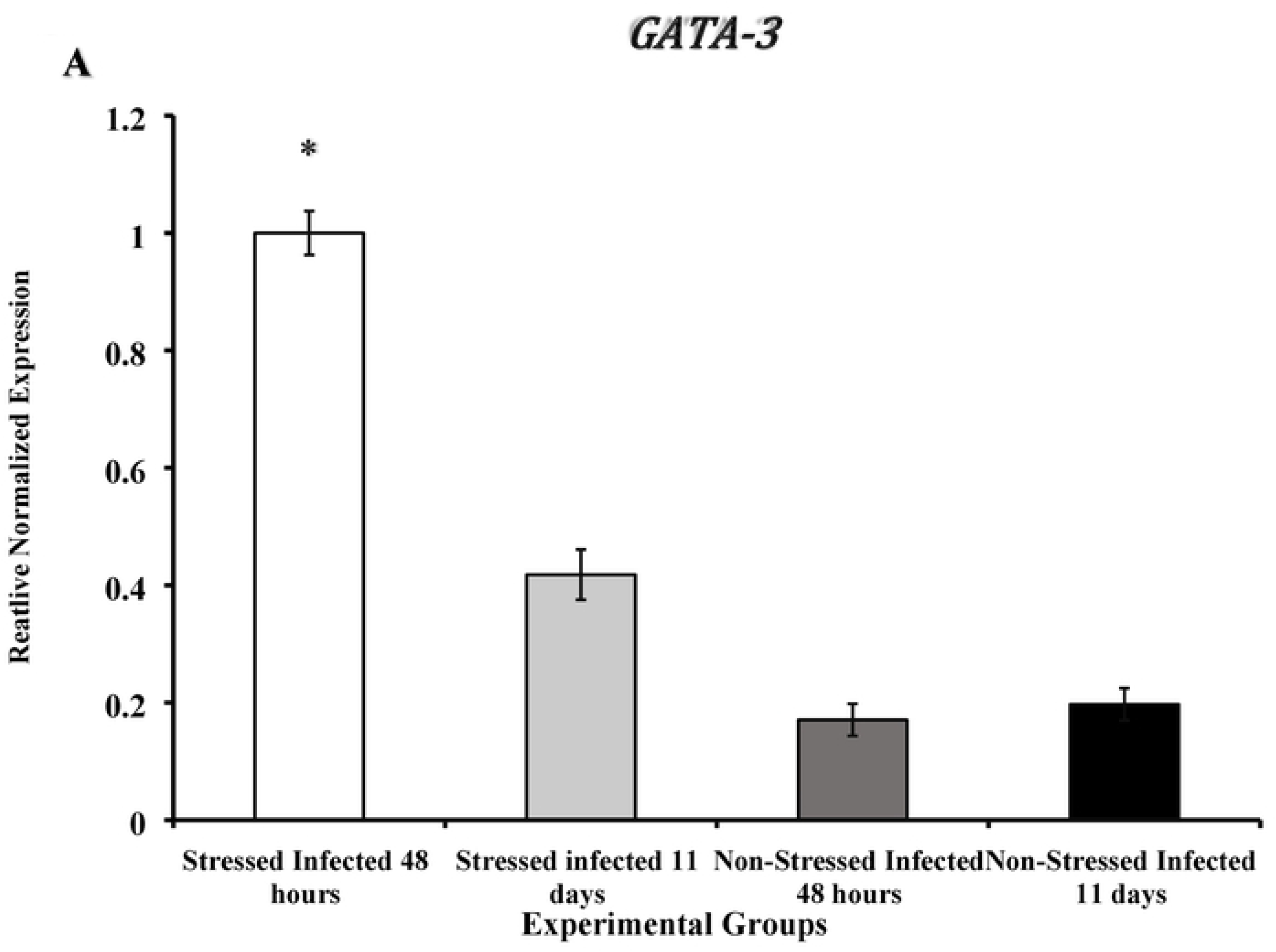
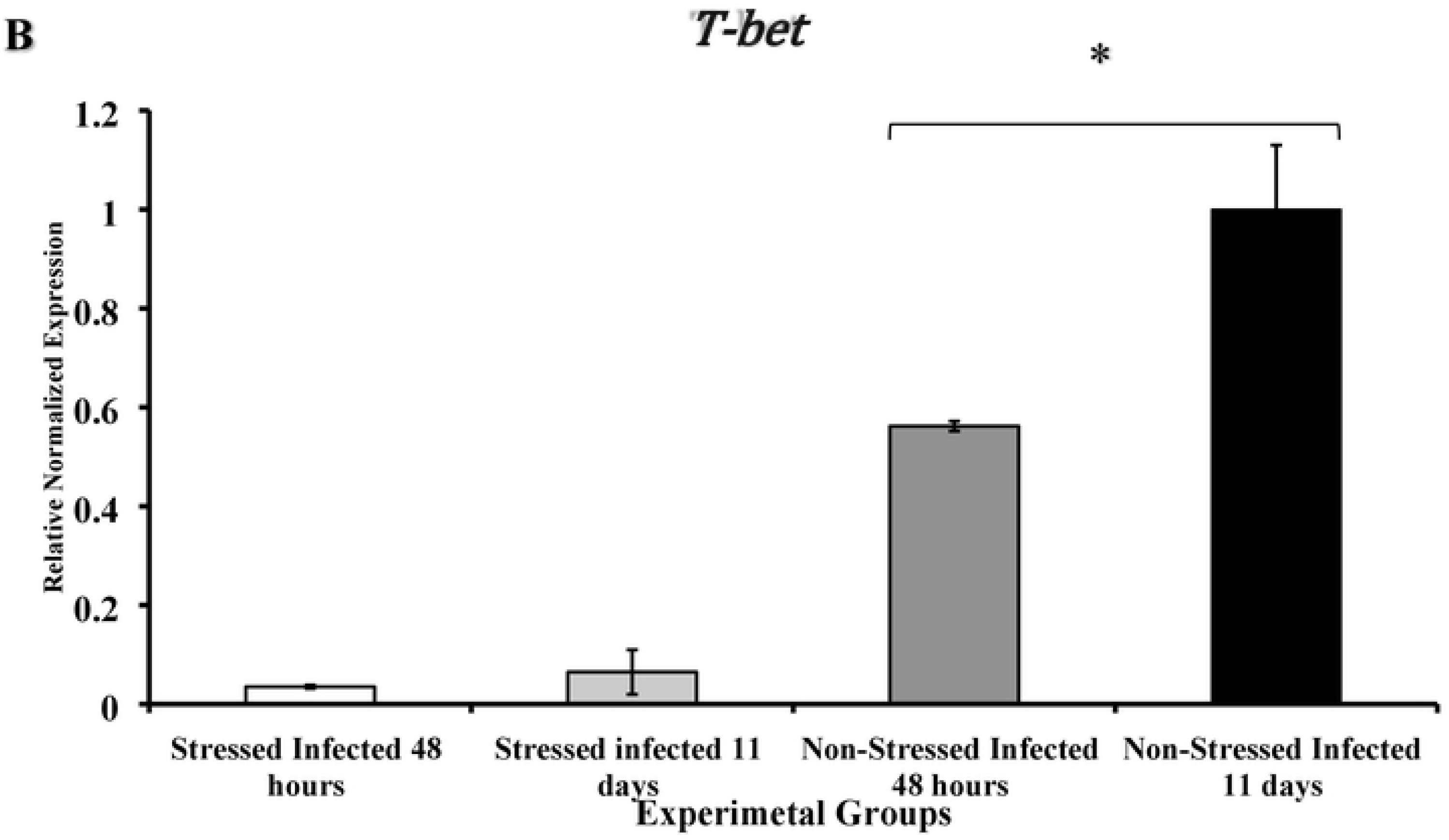
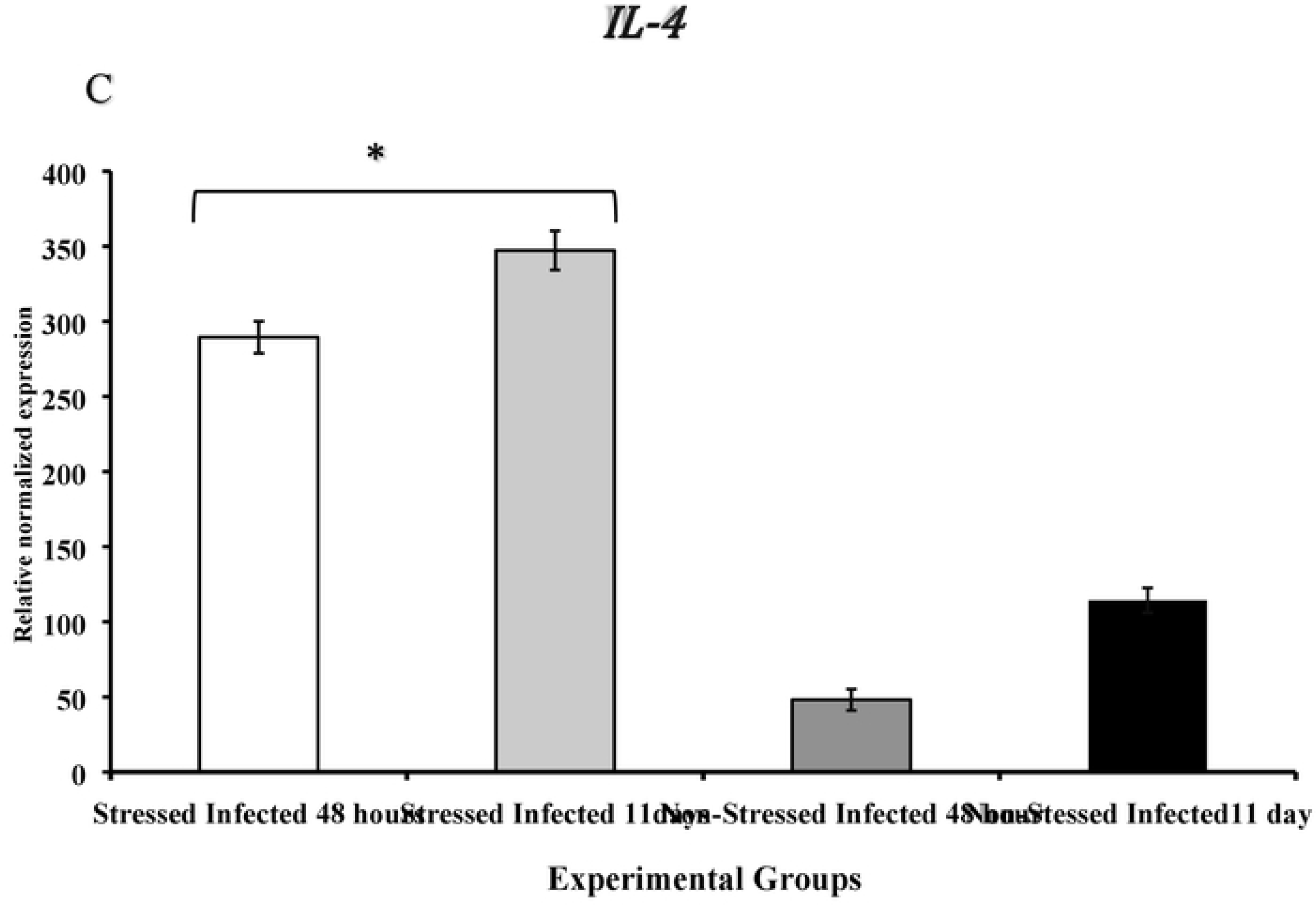
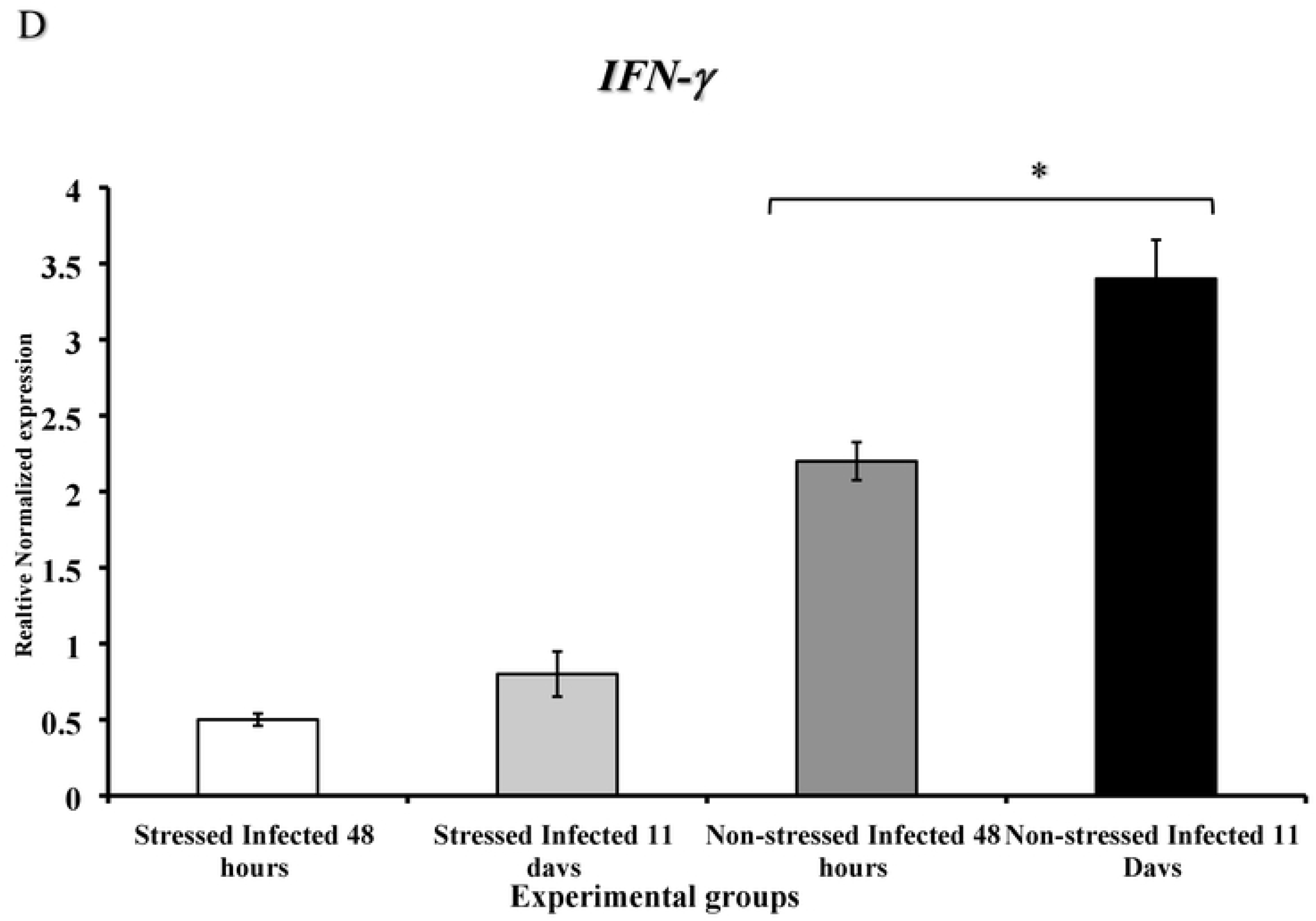
The persistence of the effect of cold-induced stress on modulation of transcription factors and cytokines in the genital tract of stressed/non-stressed mice. Gene expression of (**A**), GATA-3, (**B**) T-bet IL-4 (**C**), and (**D**) interferon-gamma during 48- or 11-days post *Chlamydia muridarum* genital infection. Results are either a representative or a mean +/- SEM of two independently performed experiments. *Denotes significant statistical differences between treatment groups at the level of (p < 0.05).

### Pre-exposure of splenic mouse naïve CD4+ T cells to β2-AR agonist or antagonist alters gene expression patterns of transcription factors and signature cytokines

Whether the gene expression of signature cytokines of splenic naïve CD4+ T cells or transcription factors and are influenced by exposure to β2-AR agonist or antagonist were tested. As shown in **Figure 7A**, treatment of naïve CD+ T cells with feroterol resulted in a marked increase of IL-4 gene expression compared to cells only. In contrast, exposure of naïve CD+ T cells to the β2-AR antagonist, ICI118, 551 resulted in elevated IL-12 gene expression (**Figure 7B**). On the other hand, treatment of naïve CD+ T cells with feroterol resulted in a marked decrease of T-bet gene expression compared to control cells, whereas exposure of naïve CD+ T cells to the β2-AR antagonist ICI.118, 551 resulted in restoring gene expression of T-bet in stressed mice. Similarly, in the presence of feroterol, the threshold cycle (Ct) value of IL-4 of stressed mice had the lowest value, which is directly correlated with up-regulated gene expression compared to all the treatment groups (**Figure 7C**), whereas, IFN-γ had the highest Ct value in the group (**Figure 7D**). A direct correlation between the Ct values and fold-changes was observed in all the treatment groups.

**Figure 7:**
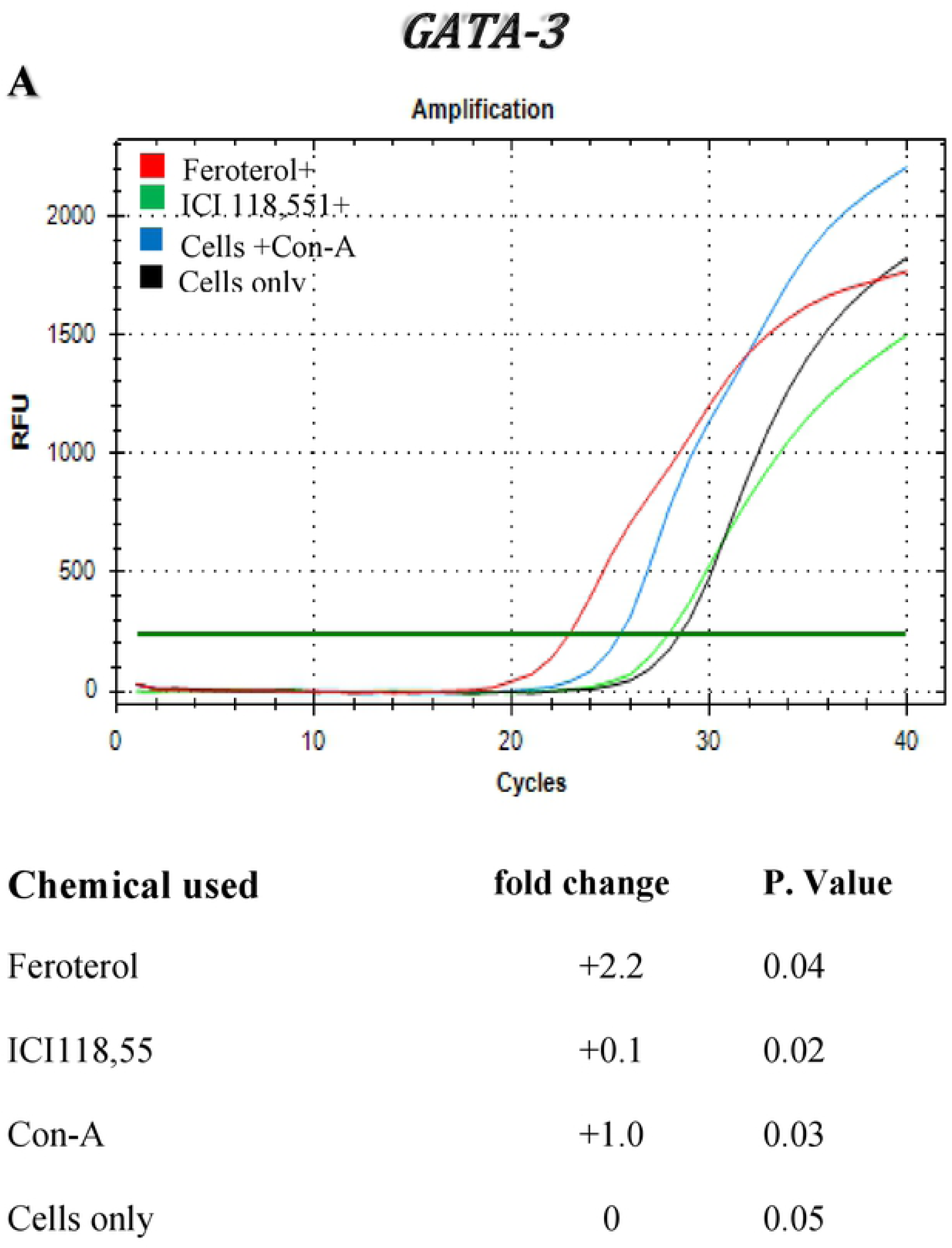
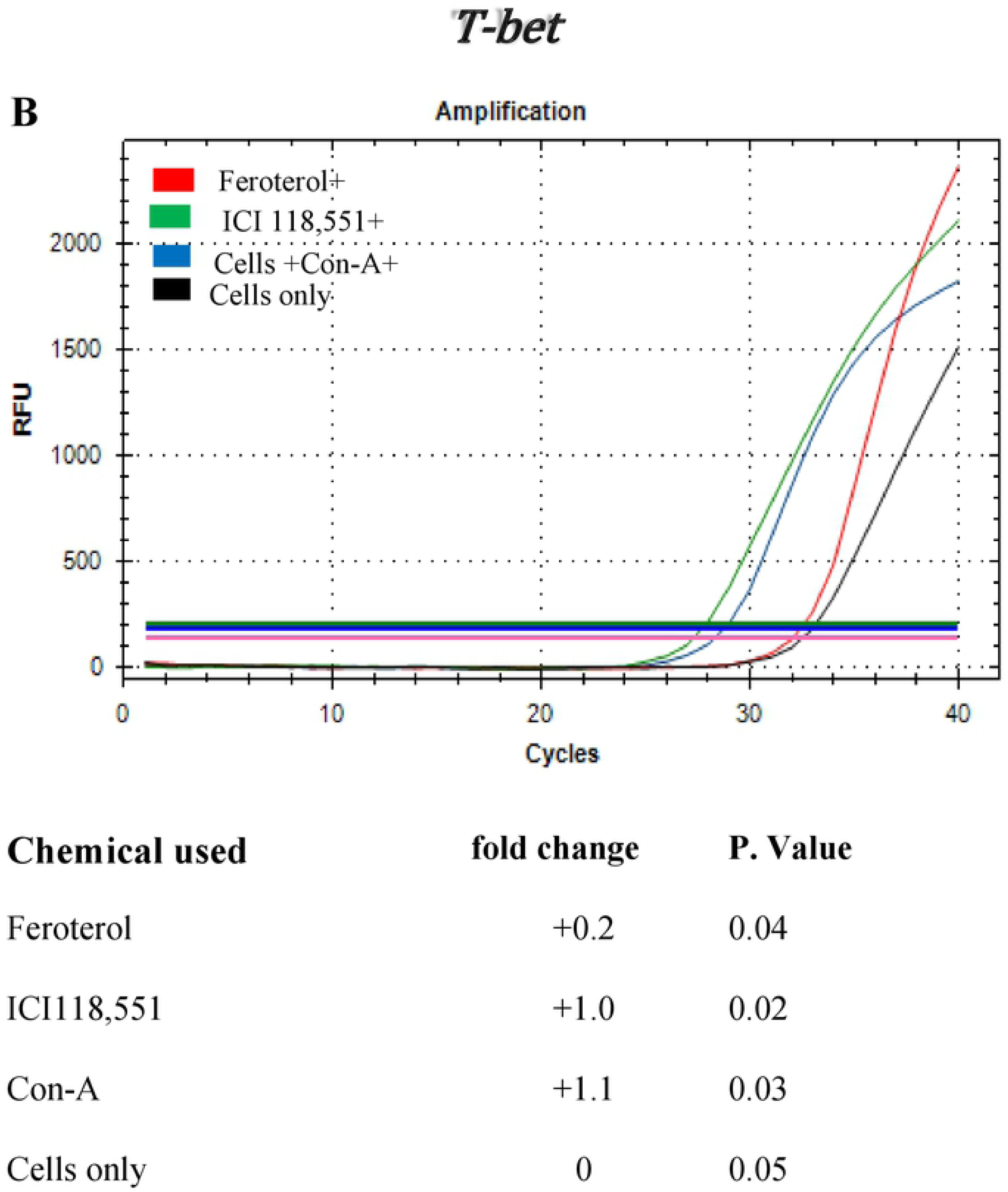
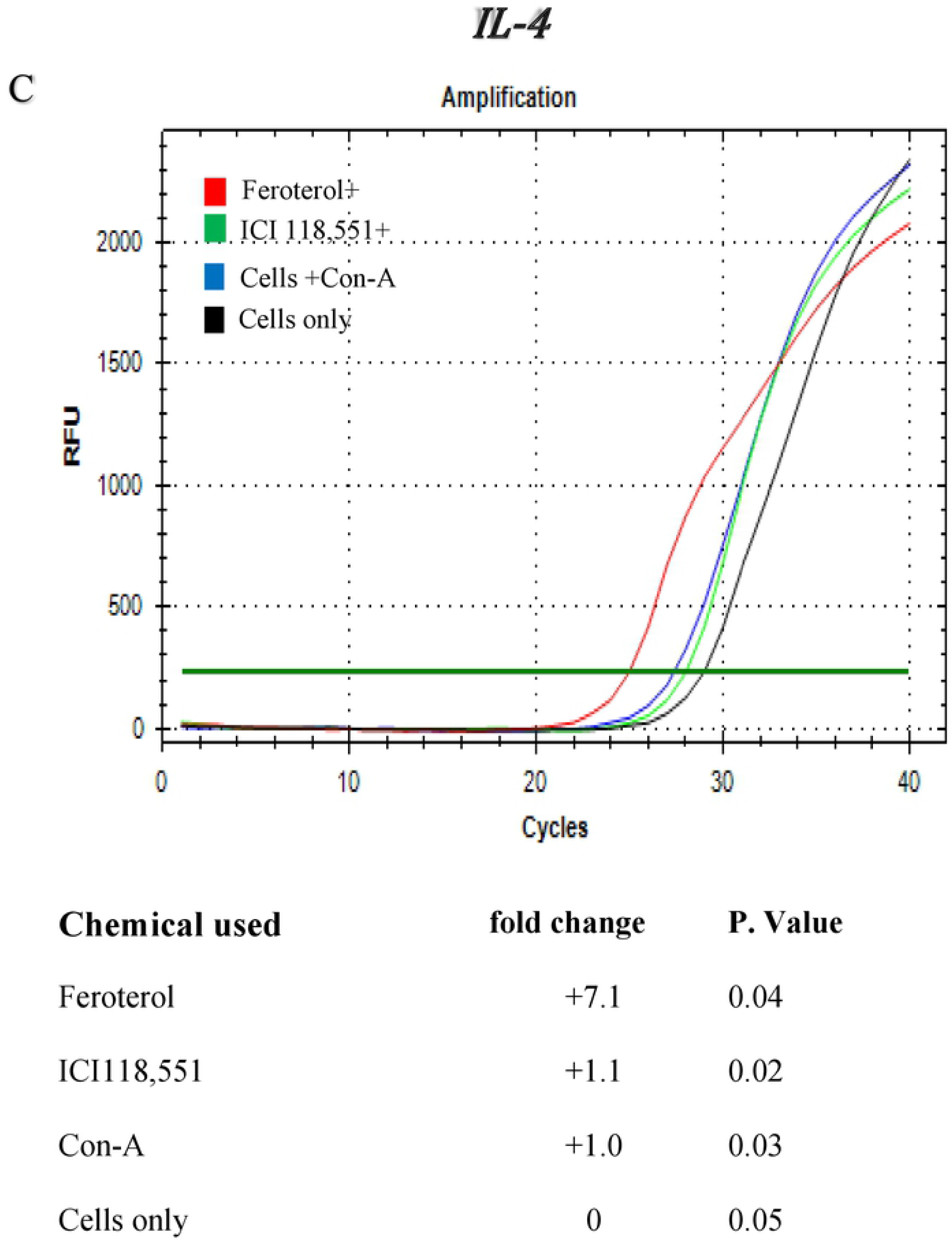
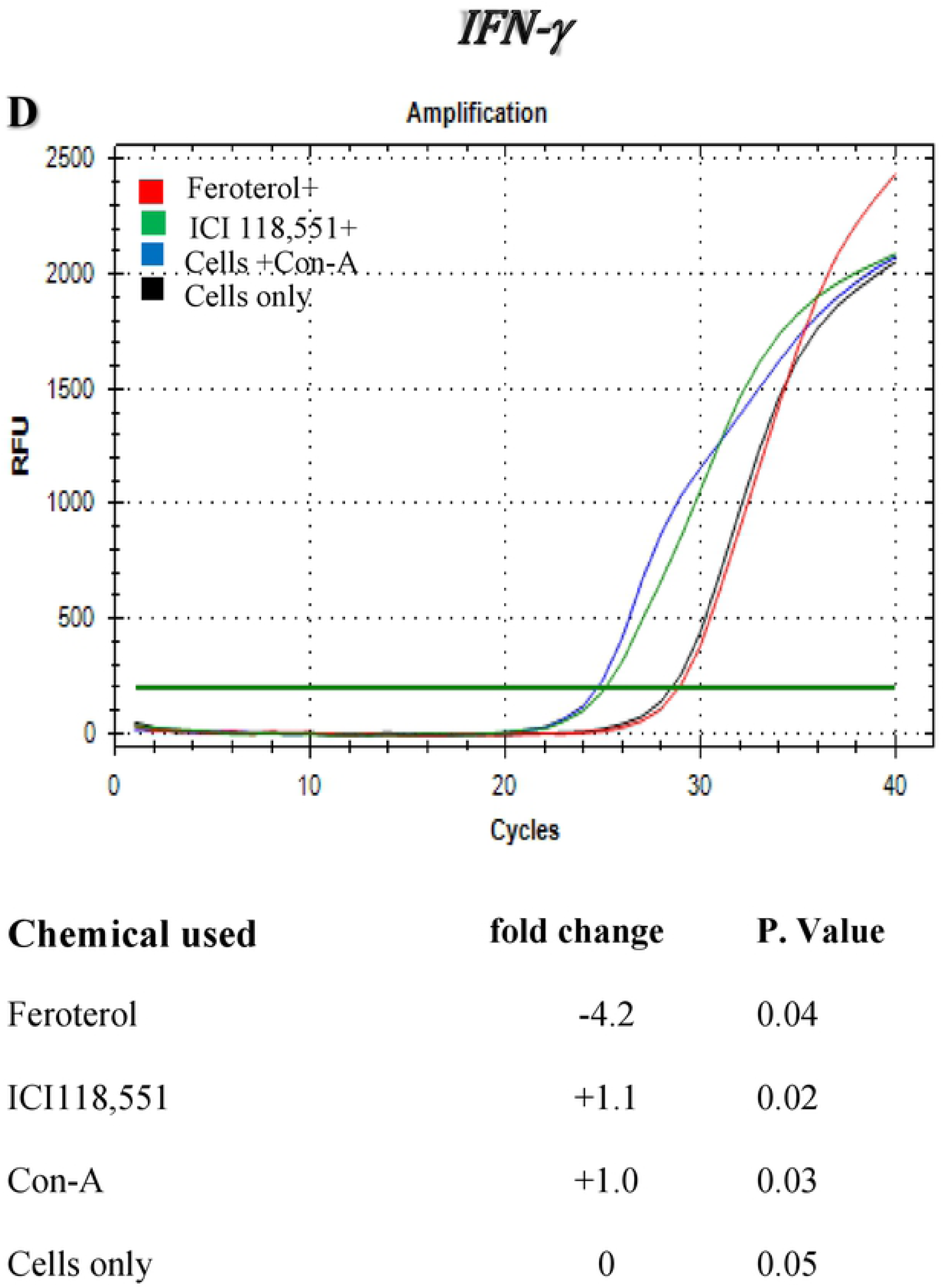
Influence of beta2-adrenergic receptor agonist and antagonist on expression pattern of (**A**) IL-4, (**B**) IL-12, (**C**) GATA-3, and (**D**) T-bet during *in vitro* proliferation of splenic naïve CD4+ T cells along fold-changes of mRNA levels are shown. Immature dendritic cell (iDC) culture was pre-exposed to NE (1 μM), fenoterol (β2-AR antagonist) (1 μM) and ICI 118,551 (β2-AR antagonist) (1 μM) for 1 h before stimulation with LPS (5 μg/mL) for 24 h before RNA isolation. Data shown are a representative of two or more independent experiments ran on different dates.

### Preexposure of splenic naïve CD4+ T cells to β2-AR agonist or antagonist affects the production of signature cytokines

In this study, production of IL-4 and IFN-γ after exposure of CD4+ T cells to a β2-AR agonist was evaluated. As shown in **Figure 8A**, the production of IL-4 in naïve CD4+ T cells was substantially increased with exposure to the β2-AR agonist Feroterol, whereas pre-exposure to β2-AR antagonist ICI118,551 resulted in a significantly reduced production of IL-4 (P<0.05). As shown in **Figure 8B**, the production of interleukin-12 (IL-12) in naïve CD+ T cells was considerably decreased due to pre-exposure to the β2-AR agonist Feroterol, whereas pre-exposure with ICI118,551 resulted in a significantly increased production of IL-12 (p <0.05). Production of IL-23 in naïve CD4+ T cells was substantially increased in the presence of Feroterol, but decreased in the presence of ICI118,551 (**Figure 8C**). This data suggests that β2-AR signaling may play an important role in modulating DC function as an essential regulator of the immune system during chlamydia genital infection.

**Figure 8:**
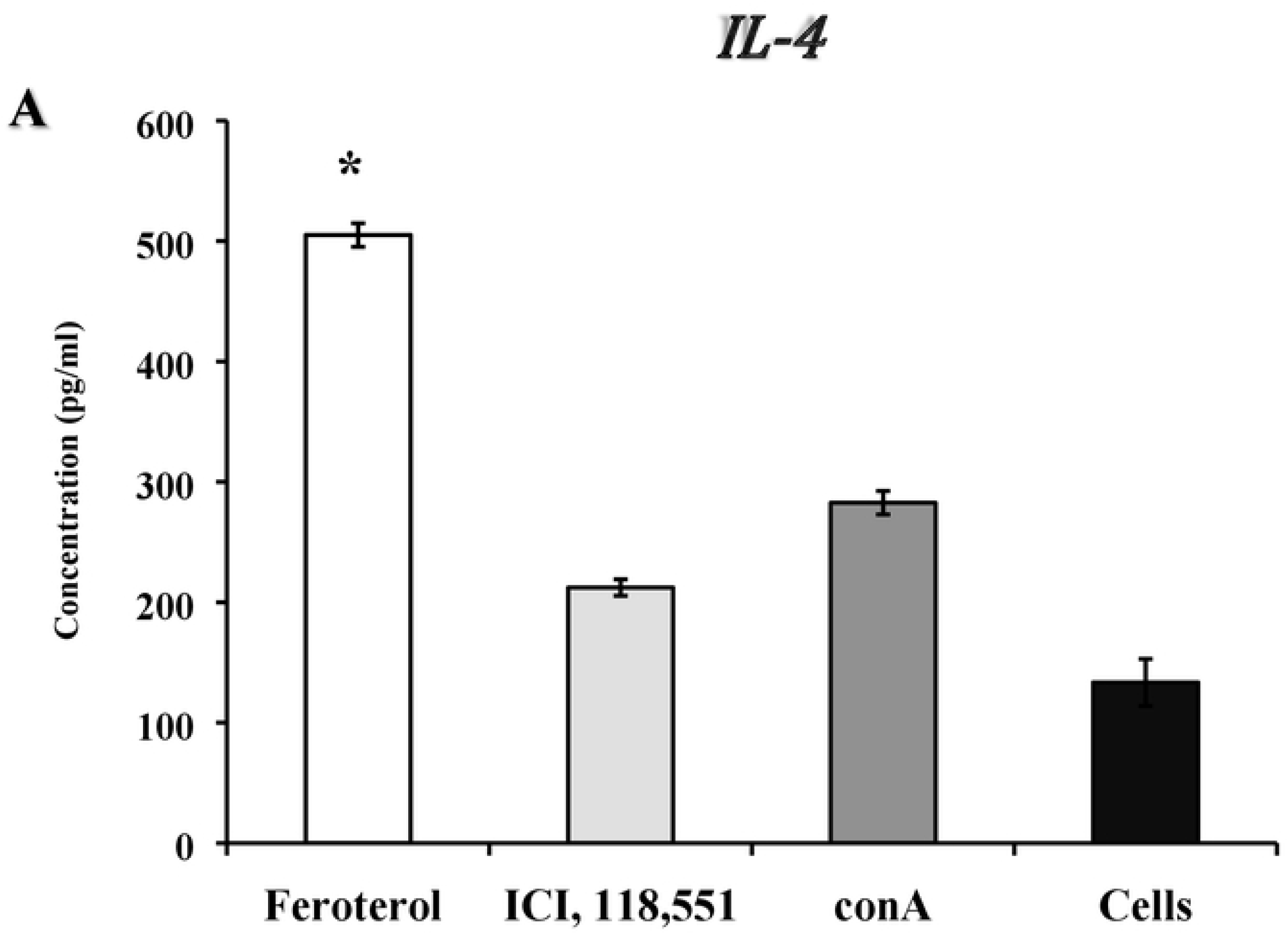
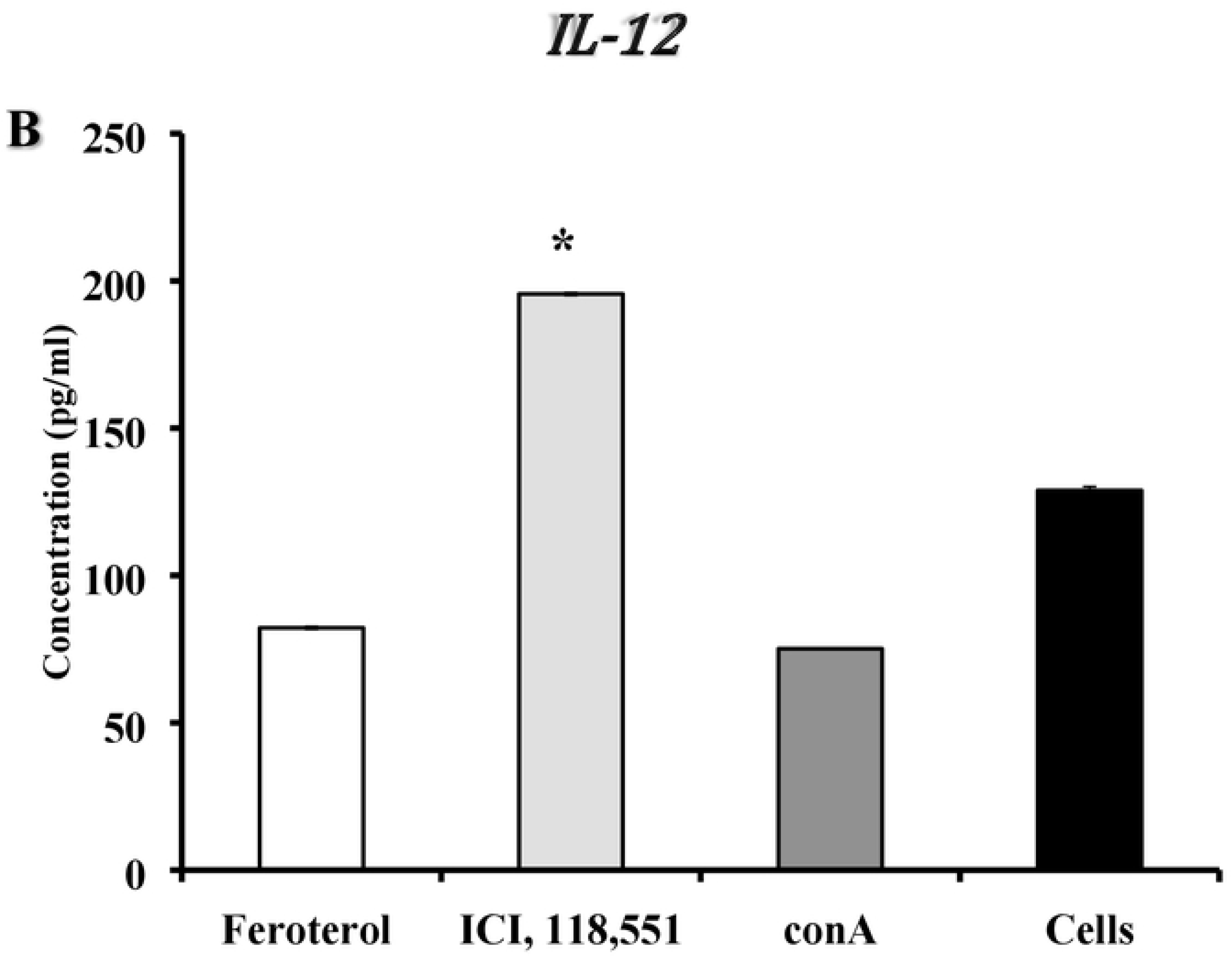
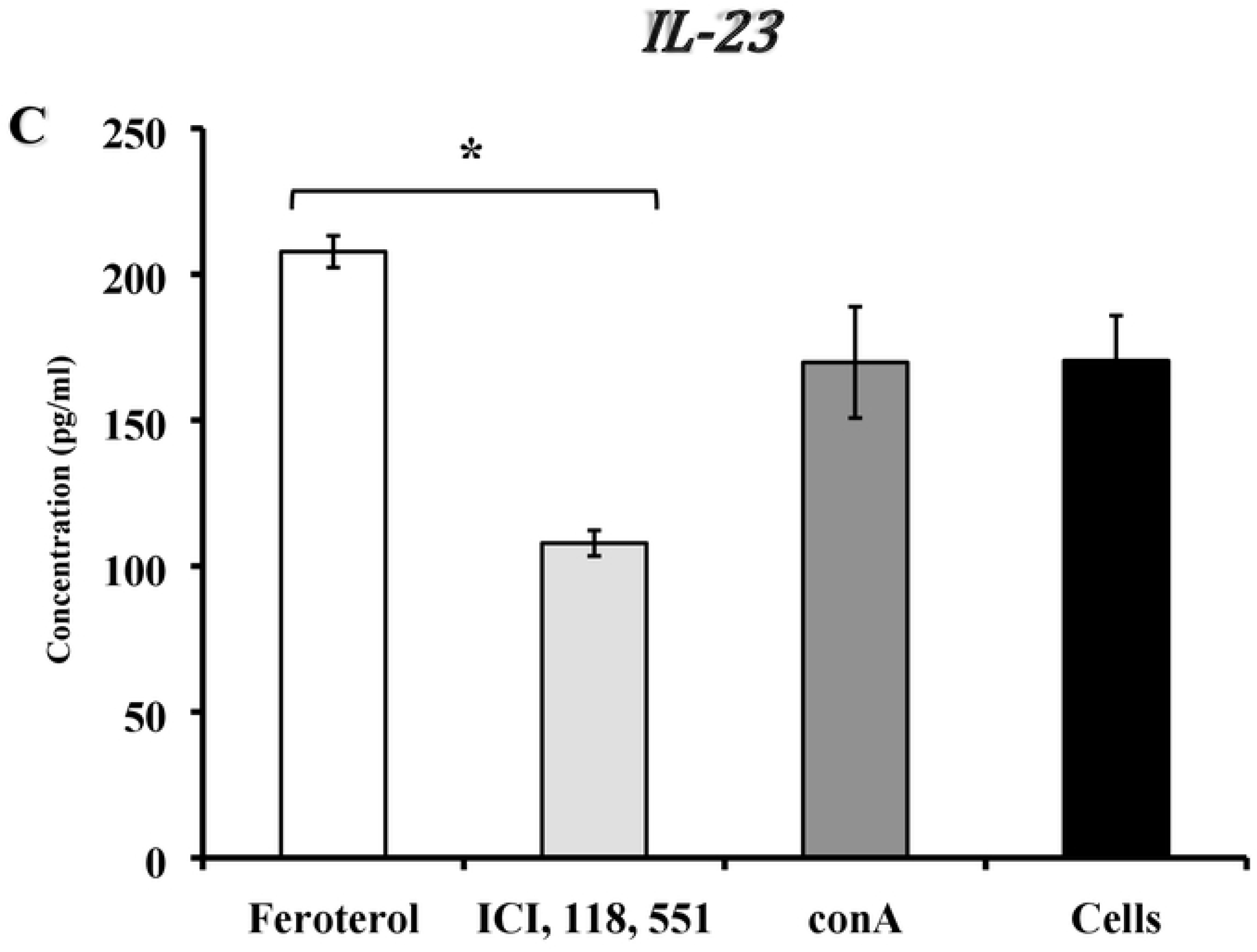
Presence of β2-AR agonist or antagonist on *in vitro* production of (**A**) IL-4, (**B**) IL-12, and (**C**) IL-23 in naïve CD4 T-cells isolated from stressed non-infected mice. Immature dendritic cell (iDC) culture was pre-exposed to NE (1 μM), fenoterol (β2-AR antagonist) (1 μM) and ICI 118,551 (β2-AR antagonist) (1 μM) for 1 h before stimulation with LPS (5 μg/mL) for 24 h before RNA isolation. Data shown are a representative or a mean +/_ SEM of two or more independent experiments ran on different dates. *Denotes significant statistical differences between treatment groups at the level of (p < 0.05).

### Norepinephrine, β2-AR agonist, or β2-AR antagonist modulates the differentiation of bone marrow derived dendritic cells (BMDCs) cultured *in vitro*

Culturing of immature BMDC in the presence or absence of norepinephrine, β2-AR and β2-AR *in vitro* prior to stimulation with LPS was performed for 24 h and culture supernatants were collected for the detection of cytokine production using ELISA. As shown in **Table2**, Unexpectedly, preexposure to feroterol, β2-AR agonist resulted in a marked increase of IL-12 in non-stressed-non-infected mice compared to that of non-stressed-infected mice. However, β2-AR antagonist, ICI 118,551, treatment resulted in increasing production of IL-12 by BMDCs of stressed mice compared to that of non-stressed mice. Pretreatment with NE, feroterol, showed a significant decrease in IL-12 production between non-stressed-infected and non-stressed-non-infected (**Table 3**). However, ICI 118,551 resulted in a significant increase of IL-12 production particularly in stressed-infected mice (**Table 3**). Pretreating of BMDCs obtained from stressed mice with NE, Feroterol, or ICI118,551 resulted in significantly decreased production of IL-10 compared to that of BMDCs obtained from non-stressed-non-infected mice (**Table 4**).

**Table 2:** Preexposure of immature bone marrow derived dendritic cells (BMDCS) of non-stressed mice to norepinephrine, β2-AR agonist, or β2-AR antagonist resulted in a variable IL-12 production with/without *Chlamydia muridarum* genital infection.

| **<u>Treatment Groups<sup>a</sup></u>** | **<u>Concentrations (pg/mL)</u>** |
| --- | --- |
| Non-stressed-non-infected +norepinephrine | 476.4 +/-108.6 |
| Non-stressed-non-infected + Ferterol<sup>b</sup> | 855.0 +/-26.7 |
| Non-stressed-non-infected + ICI 118,551 | 565.3 +/-168.3 |
| Non-stressed-infected + norepinephrine | 602.6 +/-102.81 |
| Non-stressed-infected + Feroterol | 494.4 +/-138.4 |
| Non-stressed-infected + ICI 118,551<sup>c</sup> | 735.5 +/-91.6 |
| Non-stressed-infected only | 450.5 +/-127.3 |

<sup>a</sup> Immature BMDCs of non-stressed mice were pretreated with norepinephrine, β2-AR agonist, or β2-AR antagonist for 15 minutes prior proliferation for 24 hours. Culture supernatants were tested for cytokine production using ELISA.
<sup>b</sup> Pretreatment with feroterol resulted in a significant increase in production of IL-12 in non-infected mice ( $P < 0.05$ ).
<sup>c</sup> Pretreatment with ICI 118, 551 resulted in a significant increase in production of IL-12 compared to non-stressed and infected.
Immature dendritic cell (iDC) culture was pre-exposed to NE (1 μM), fenoterol ( β2-AR antagonist) (1 μM) and ICI 118,551 ( β2-AR antagonist) (1 μM) for 1 h before stimulation with LPS (5 μg/mL) for 24 h before RNA isolation. Data shown are a representative or a mean ± SEM of two or more independent experiments ran on different dates. \*Denotes significant statistical differences between treatment groups at the level of ( $p < 0.05$ ).

**Table 3:** Preexposure of immature bone marrow derived dendritic cells (BMDCS) of stressed mice to norepinephrine, β2-AR agonist, or β2-AR antagonist resulted in a variable IL-12 production with/without *Chlamydia muridarum* genital infection.

| **<u>Treatment Groups<sup>a</sup></u>** | **<u>Concentrations (pg/mL)</u>** |
| --- | --- |
| Stressed-non-infected + norepinephrine | 93.6 ± 40.5 |
| Stressed-non-infected + Forterol | 131.2 ± 126.7 |
| Stressed-non-infected + ICI 118,551 | 396.5 ± 181.0 |
| Stressed-infected + norepinephrine | 120.8 ± 10 |
| Stressed-infected + Forterol<sup>b</sup> | 157.6 ± 38.3 |
| Stressed-infected + ICI, 118551<sup>c</sup> | 901.6 ± 80.4 |
| Stressed-infected only | 300.3 ± 12.5 |

<sup>a</sup> Immature BMDCs of stressed mice were pretreated with norepinephrine, β2-AR agonist, or β2-AR antagonist for 15 minutes prior proliferation for 24 hours. Culture supernatants were tested for cytokine production using ELISA.
<sup>b</sup> Pretreatment with ICI, 118 551 resulted in a significant increase in production of IL-12 in stressed and infected mice ( $P < 0.05$ ).
<sup>c</sup> Pretreatment with exogenous chemicals norepinephrine or forterol resulted in a significant decrease in production of IL-12 compared to the stressed and infected only.
Immature dendritic cell (iDC) culture was pre-exposed to NE (1 μM), fenoterol ( β2-AR antagonist) (1 μM) and ICI 118,551 ( β2-AR antagonist) (1 μM) for 1 h before stimulation with LPS (5 μg/mL) for 24 h before RNA isolation. Data shown are a representative or a mean ± SEM of two or more independent experiments ran on different dates. \*Denotes significant statistical differences between treatment groups at the level of ( $p < 0.05$ ).

**Table 4:** Preexposure of immature bone marrow derived dendritic cells (BMDCS) of stressed mice to norepinephrine, β2-AR agonist, or β2-AR antagonist resulted in a variable IL-10 production with/without *Chlamydia muridarum* genital infection.

| **<u>Treatment Groups<sup>a</sup></u>** | **<u>Concentrations (pg/ml)</u>** |
| --- | --- |
| Non-stressed-non-infected +norepinephrine | 321.6 +/-266.0 |
| Non-stressed-infected + Formoterol<sup>b</sup> | 379.8 +/-126.0 |
| Non-stressed- infected + ICI 118,551 | 246.5 +/-82.3 |
| Stressed-non-infected + norepinephrine | 34.7 +/-8.6 |
| Stressed-non-infected + Formoterol | 50.7 +/-0 |
| Stress only | 429 +/-3 |

<sup>a</sup> Immature BMDCs of non-stressed mice were pretreated with norepinephrine, β2-AR agonist, or β2-AR antagonist for 15 minutes prior proliferation for 24 hours. Culture supernatants were tested for cytokine production using ELISA.
<sup>b</sup> Pretreatment with feroterol resulted in a significant increase in production of IL-10 in non-infected mice compared to stressed mice ( $P < 0.05$ ).
<sup>c</sup> Pretreatment with ICI 118, 551 resulted in a significant decrease in production of IL-10 compared to non-stressed and infected ( $p < 0.05$ ).

### Cold-induced stress results in differential gene expression of β-AR subtypes in splenic dendritic cells

BMDCs harvested from the femurs and tibias of stressed and non-stressed mice were allowed to differentiate in complete RMPI for 8 days in the presence of 20 ng/mL GM-CSF. Immature BMDCs were pre-exposed to norepinephrine, Feroterol or ICI118,551 for 20 minutes then stimulated with 1 μg/mL LPS for 24 h to obtain mature BMDCs. We hypothesized cold induced-stress results in increased gene expression of β2-AR and decreased gene expression of β1-AR and β3-ARs in splenic DCs. Purified splenic DCs from stressed and non-stressed mice were proliferated with/without LPS for RNA isolation and real-time PCR analysis. As shown in **Figure 9A**, gene expression β2-AR BMDCs was slightly up-regulated in stressed mice in the presence of LPS. On the other hand, stress and stimulation with LPS resulted in little or no difference in gene expression of β1-AR and β3-AR in splenic DCs (**Figures 9B, 9C**). Gene expression β2-AR BMDCs was up regulated whereas little or no difference in gene expression of β1-AR and β3-AR in BMDCs (**data not shown**). The relative expression of β2-AR in BMDC of stressed mice was the most highly expressed compared to the relative expression of β1 or β3 in DCs.

**Figure 9:**
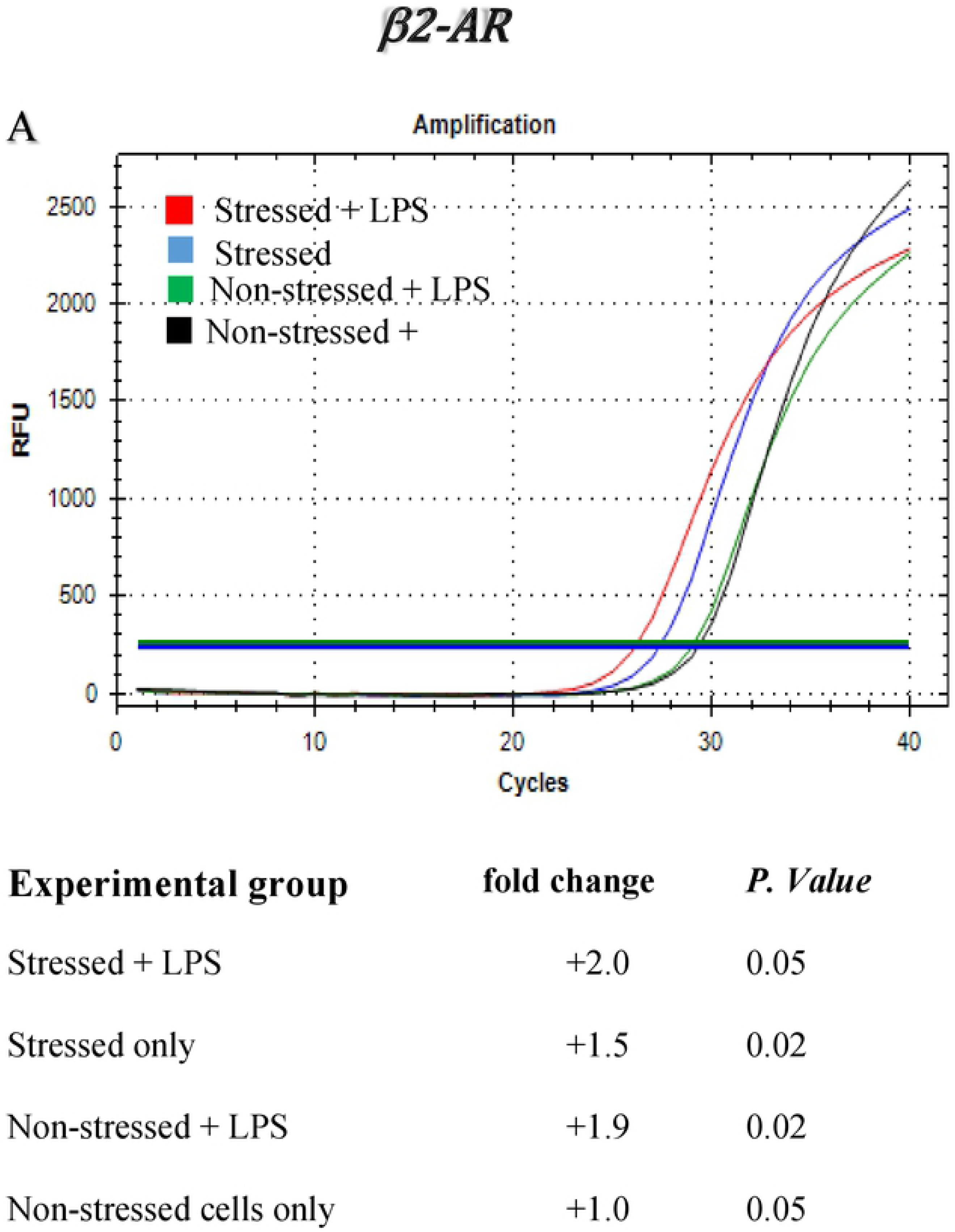
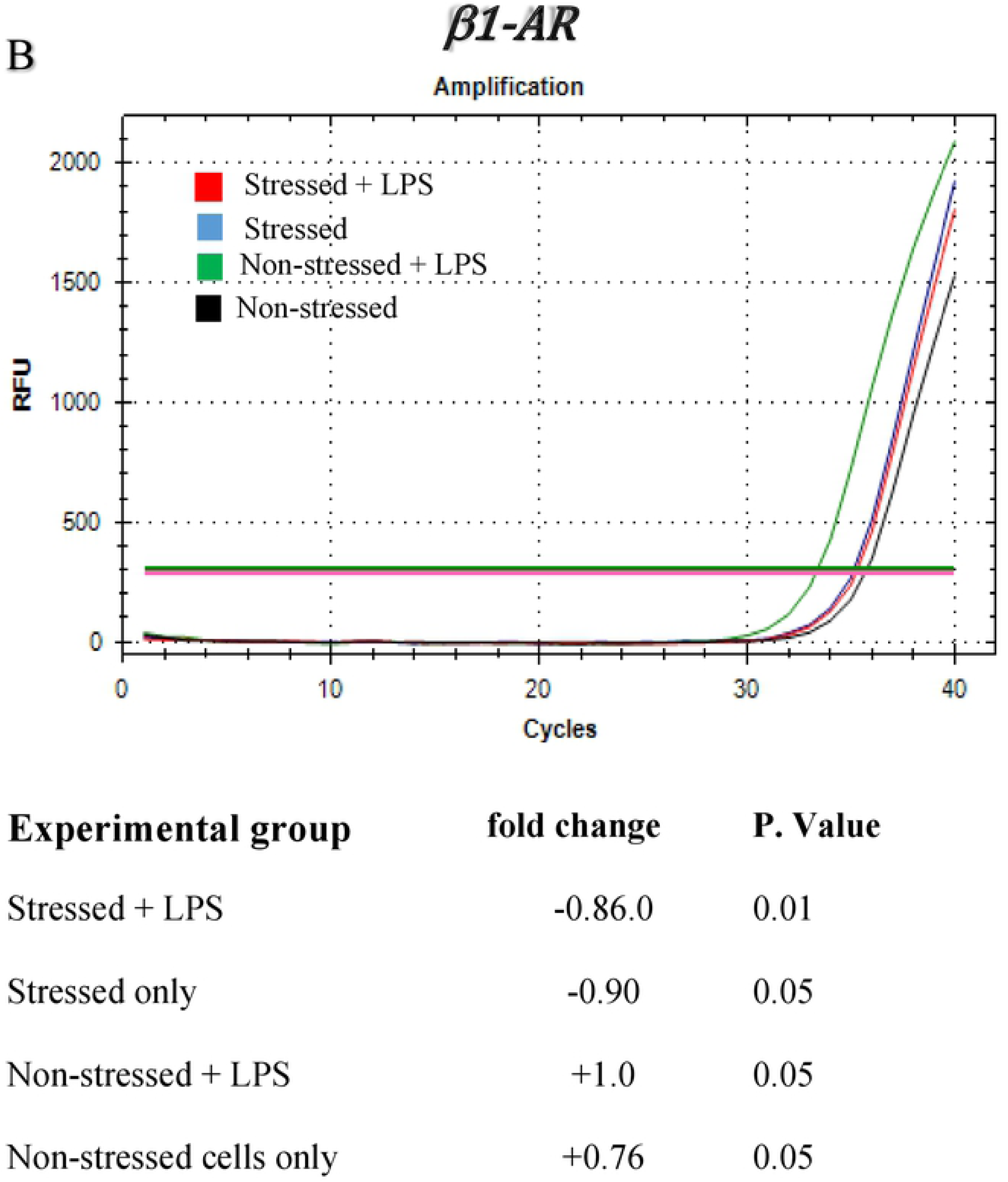
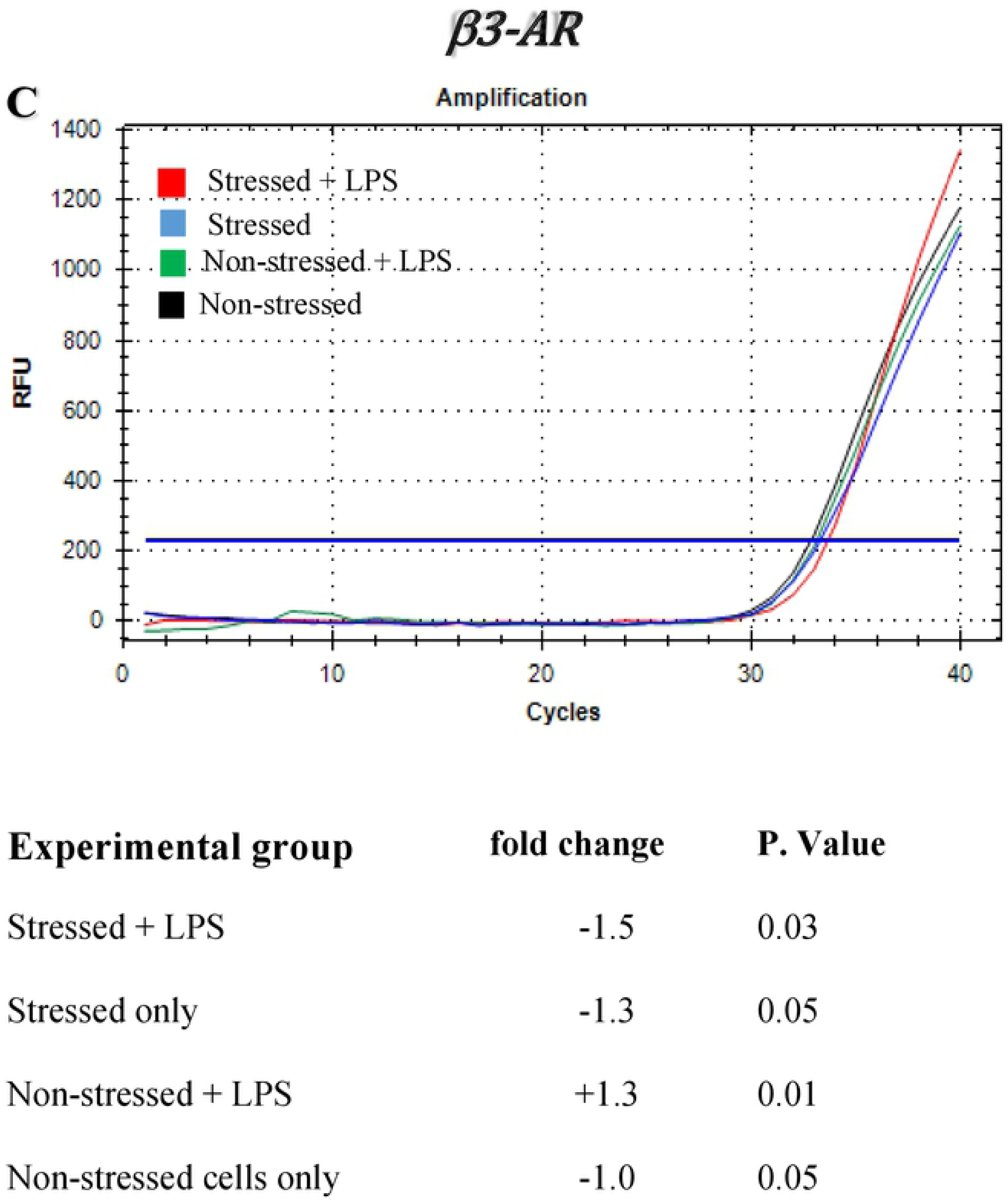
Gene expression profiles and fold change mRNA levels of adrenergic receptor subsets in splenic dendritic cells isolated stressed and non-stressed mice with/without *Chlamydia muridarum* genital infection. (**A**) β2 adrenergic receptor, (**B**) β1 adrenergic receptor, and (**C**) β3 adrenergic receptor. Shown data are representatives of two or more independent experiments ran on different dates.

### Naïve T cells co-cultured with BMDCs pretreated with β2-AR agonists and antagonists resulted in production of different levels of Th1 and Th2 cytokine production

We first determined the levels of Th1 and Th2 cytokine production without BMDC co-culturing in the absence of Anti-CD3 stimulation. The ability of DCs to induce/derive T helper cell differentiation and cytokine production was determined. We tested the activity of DCs on stimulation of CD4+ T at non-polarizing conditions *in vitro*. To clarify how cold-induced stress affects the function of DCs in our mouse model, CD4+ T cells from stressed and non-stressed mice with/without *C. muridarum* genital infection was co-cultured with BMDCs. Our results show that BMDCs pre-treated with NE or Feroterol showed decreased IL-12 production compared to non-treated T cells (**Figure 10**). Co-culturing of mature BMDCs and naïve CD4+ T cells resulted in increased production of IL-4, IL-10, and IL-17 in culture supernatants, suggesting that stimulation of β2-AR receptor leads to the increased production of Th2 cytokines (**Figure 10**). Our results demonstrate that cold-induced stress influences the activation and differentiation of BMDCs stimulated by LPS, shifting the Th1/Th2 ratio and thus impairing the production of protective Th1 cytokines.

**Figure 10:**
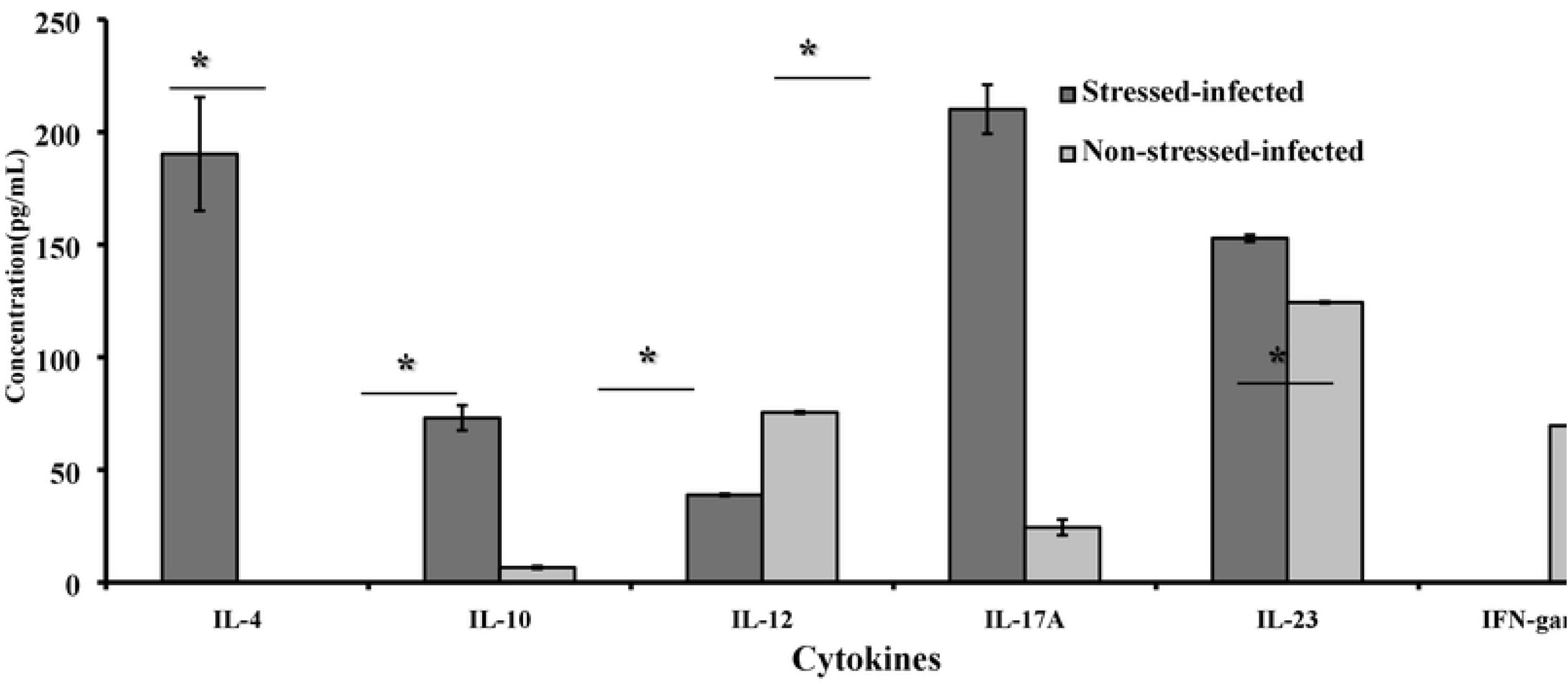
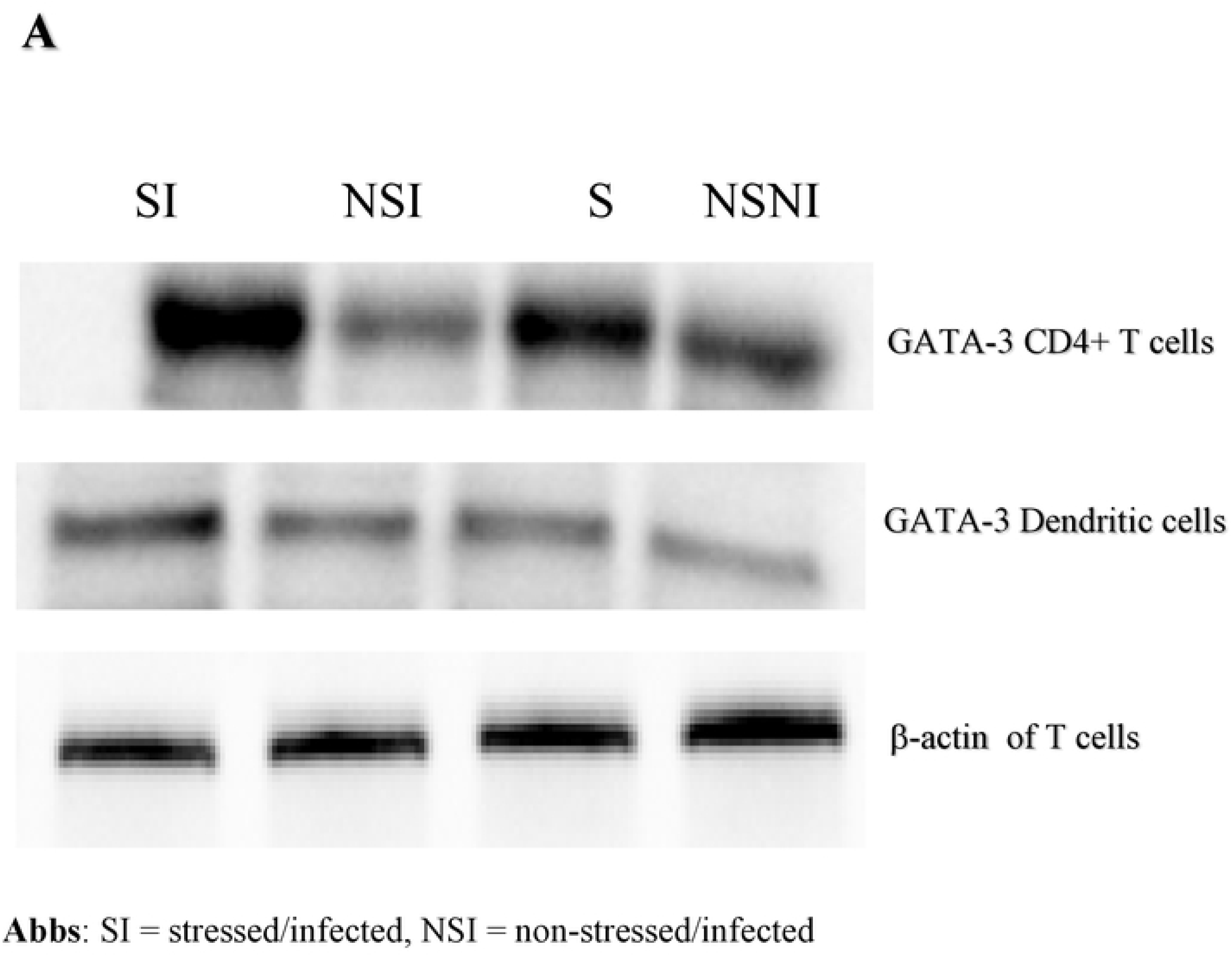
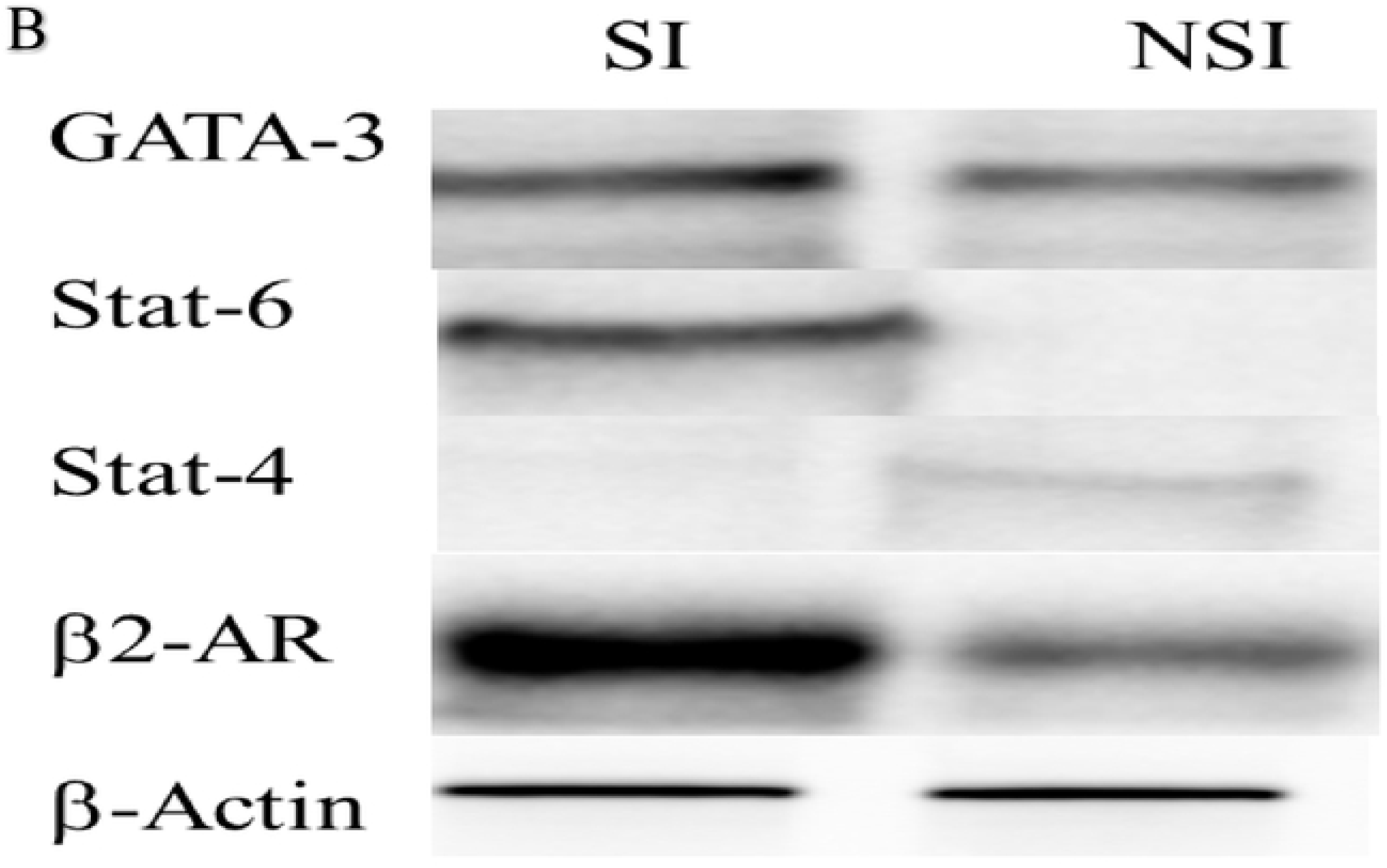
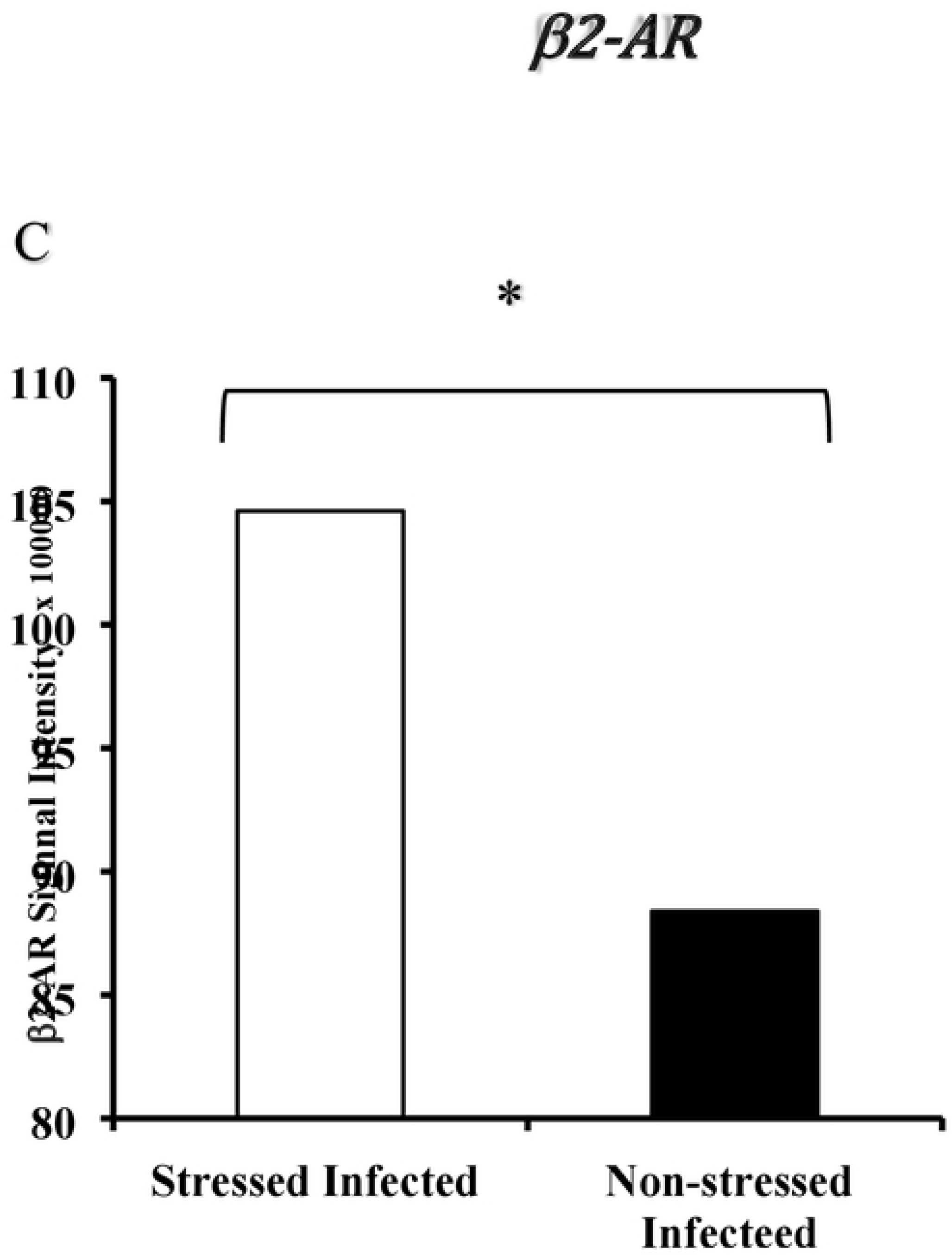
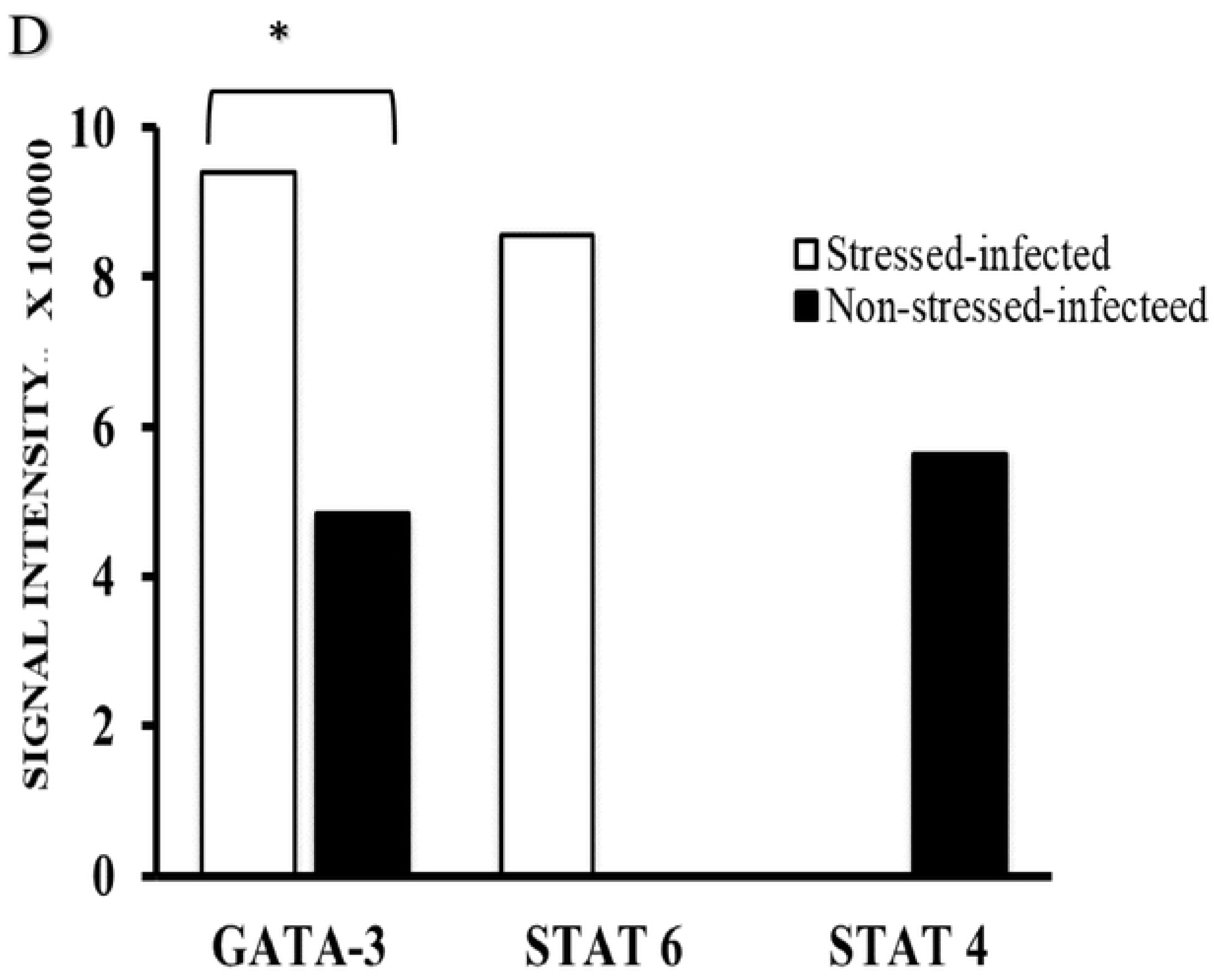
Cytokine production kinetics of CD4+T cells and dendritic cells co-cultured *in vitro.* Following maturation of BMDC using LPS, they were pre-primed for 1 h by β2-AR agonist or antagonist. CD4+ T cells from stressed and non-stressed mice isolated by negative selection were co-cultured for 48 h. Shown data are representatives of two or more independent experiments ran on separate dates. *Denotes statistically significant differences between stressed and non-stress mice.

Whether CD4+ T activation is T cell receptor (TCR)-independent or dependent, BMDC-CD4+ T cell co-culturing was in plates coated with anti-CD3 and anti-CD28 antibodies with/without feroterol or ICI 188,551(**Table 5**). Culture supernatants were collected after 48 h of proliferation for ELISA analysis. The β2-AR agonist resulted in complete blockage of IL-12 production and a significant blockage of IL-4 production, regardless of stress treatment and plate coating with anti-CD3 coating (p<0.5). β2-AR agonist feroterol resulted in a significant increase of IL-4 production in stressed mice but in plates without anti-CD3 coating (p<0.5). The production of IL-10 in stressed mice was the maximum in plates coated with antiCD3 (p<0.5). Production of IL-23 was high regardless of stress/no stress conditions and plate coating with ani-CD3 antibodies.

**Table 5:** Effect of β2-AR agonist or β2-AR antagonist on production of Th1 and Th2 cytokines by CD4+ T cells of stressed/non-stressed mice co-cultured with BMDCs on plates with/without anti-CD3 antibodies. All data represent a mean of duplicates of two experiments performed on separate dates. One-way ANOVA was used to determine group differences.

| BMDC-<br>CD4+ T<br>culture | (-) CD3 |  |  |  | (+) CD3 |  |  |  |  |  |  |
| --- | --- | --- | --- | --- | --- | --- | --- | --- | --- | --- | --- |
|  | Stressed |  |  |  | Stressed |  |  |  |  |  |  |
|  | IL-4 | IL-10 | IL-12 | IL-23 | IL-4 | IL-10 | IL-12 | IL-23 | IL-4 | IL-10 | IL-12 |
| ICI,188,551<br>+<sup>a</sup> | 37 | 123 | ND<sup>c</sup> | 191 | 26 | 74 | ND | 172 | ND | 49 | ND |
| Forterol | 192 | 104 | 74 | 160 | 36 | 577<sup>d</sup> | ND | 248 | 44 | 60 | 48 |
| CD4+ T<br>cells only | 64 | 70 | ND | 88 | 38 | 101 | ND | 74 | ND | 64 | 75 |

<sup>a</sup>The β2-AR antagonist, ICI, 188,551, resulted in complete blockage of IL-12 and a significant blockage of IL-4 production irrespective of stress treatment and plate coating with anti-CD3 coating ( $p < 0.5$ )
<sup>b</sup>The β2-AR agonist, forterol, resulted in a significant increase of IL-4 production in stressed mice in plates without anti-CD3 antibodies ( $p < 0.5$ ). Samples from non-stressed mice showed no significant stimulation or inhibition (**data not shown**).
<sup>c</sup>ND indicates non-detectable level of cytokines by ELISA
<sup>d</sup>The production of IL-10 in stressed mice was the maximum in plates coated with anti-CD3 ( $p < 0.5$ )
Immature dendritic cell (iDC) culture was pre-exposed to NE (1 μM), fenoterol ( β2-AR antagonist) (1 μM) and ICI 118,551 ( β2-AR antagonist) (1 μM) for 1 h before stimulation with LPS (5 μg/mL) for 24 h before RNA isolation. Data shown are a representative or a mean ± \_
SEM of two or more independent experiments ran on different dates. \*Denotes significant statistical differences between treatment groups at the level of ( $p < 0.05$ ).

### Western blot analysis on secretion of transcription factors and signal transducer and activator of transcriptions (STATs)

Western blot analysis confirmed that cold-induced stress treatment results in increased secretion of GATA-3 in the genital tract, suggesting that cold induced stress is associated with dysregulation of transcription factor of T-bet while favoring increased signaling of GATA-3 (**Figure 11**). Elevated expression of GATA3 transcription factor was detected in CD4+ T cells obtained from stressed mice compared to that of non-stressed mice suggesting that GATA3 is crucial for inducing key attributes of Th2 cells essential for Th2 cytokine production including IL-4. In addition to GATA-3 and T-bet expression patterns, the production of Th1 and Th2 CD4+ T cells, changes in signal transducer and activator of transcription (STAT) 4 and 6 were examined by Western bot analysis. Our data show that that CD4+ cells from stressed mice showed high intensity of STAT 6 and GATA-3 compared to non-detectable level STAT 6 of T-bet suggesting that CD4+ development is skewed toward Th2 differentiation by high GATA-3 STAT 6. We also showed up-regulation of β2-AR and these findings lead to a model of T cell differentiation that holds that naive T cells tend toward Th2 differentiation through induction of GATA-3, IL-4, STAT-6 and subsequent down-regulation of STAT4, IL-12, T-bet and IFN-γ levels.

**Figure 11.**
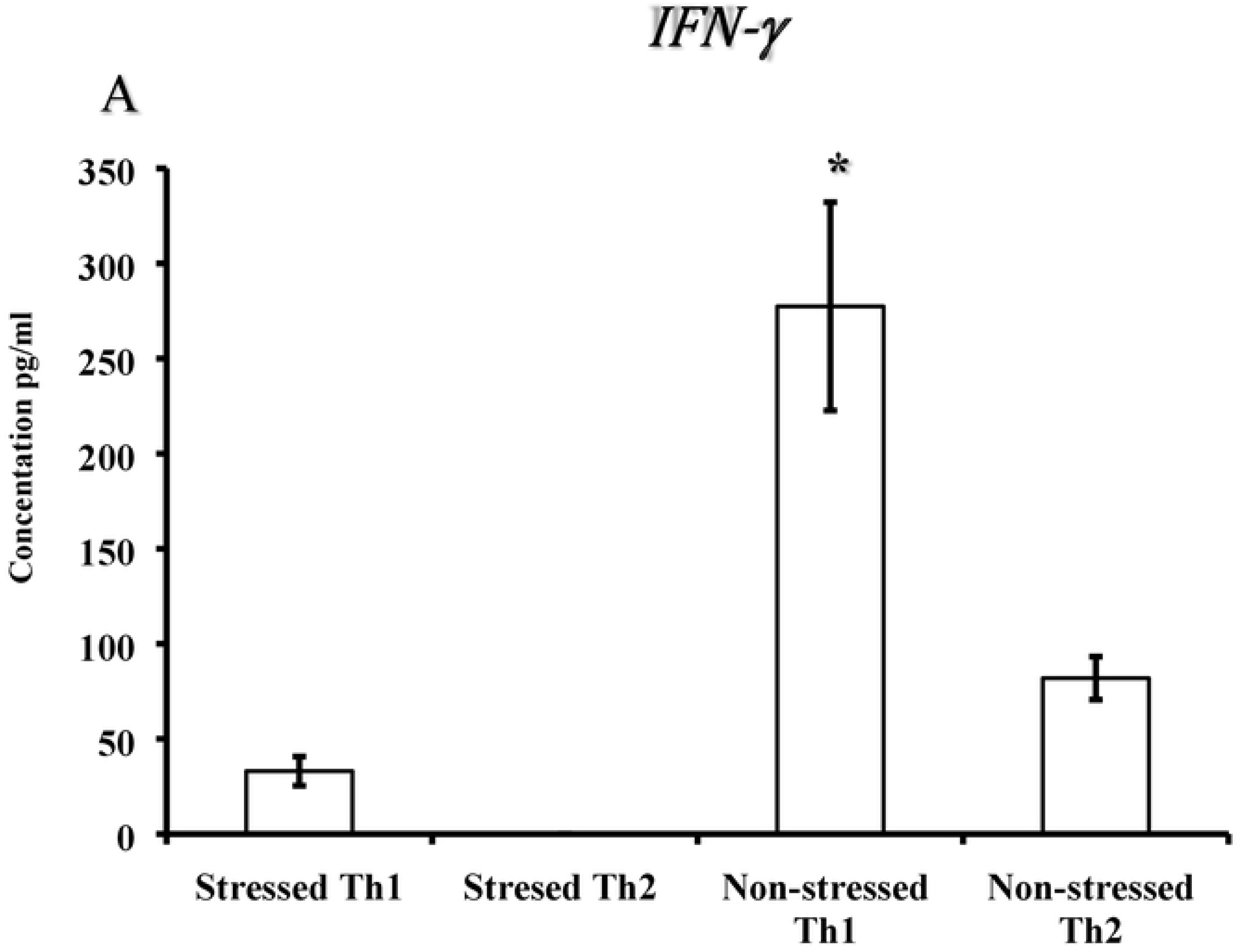
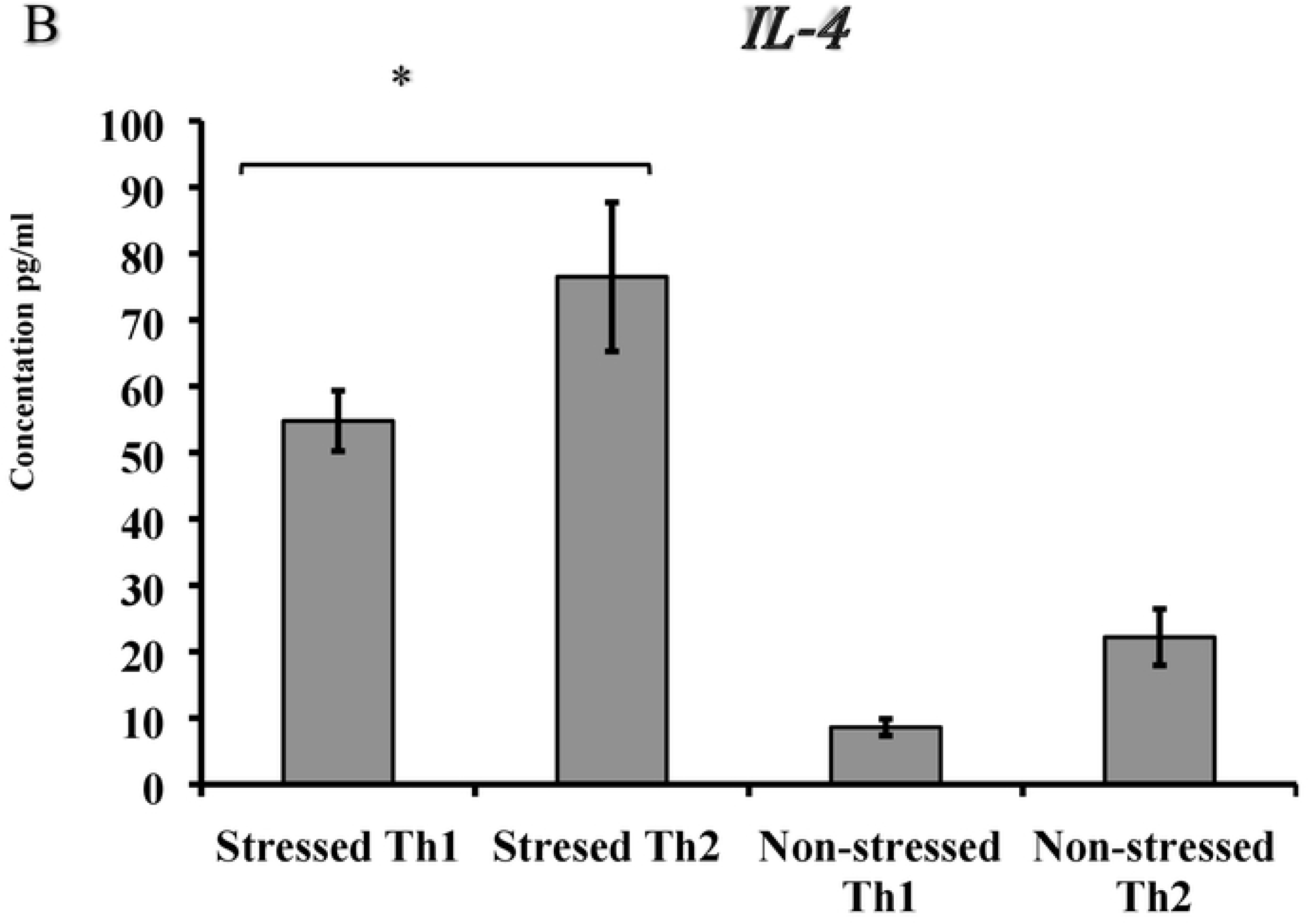
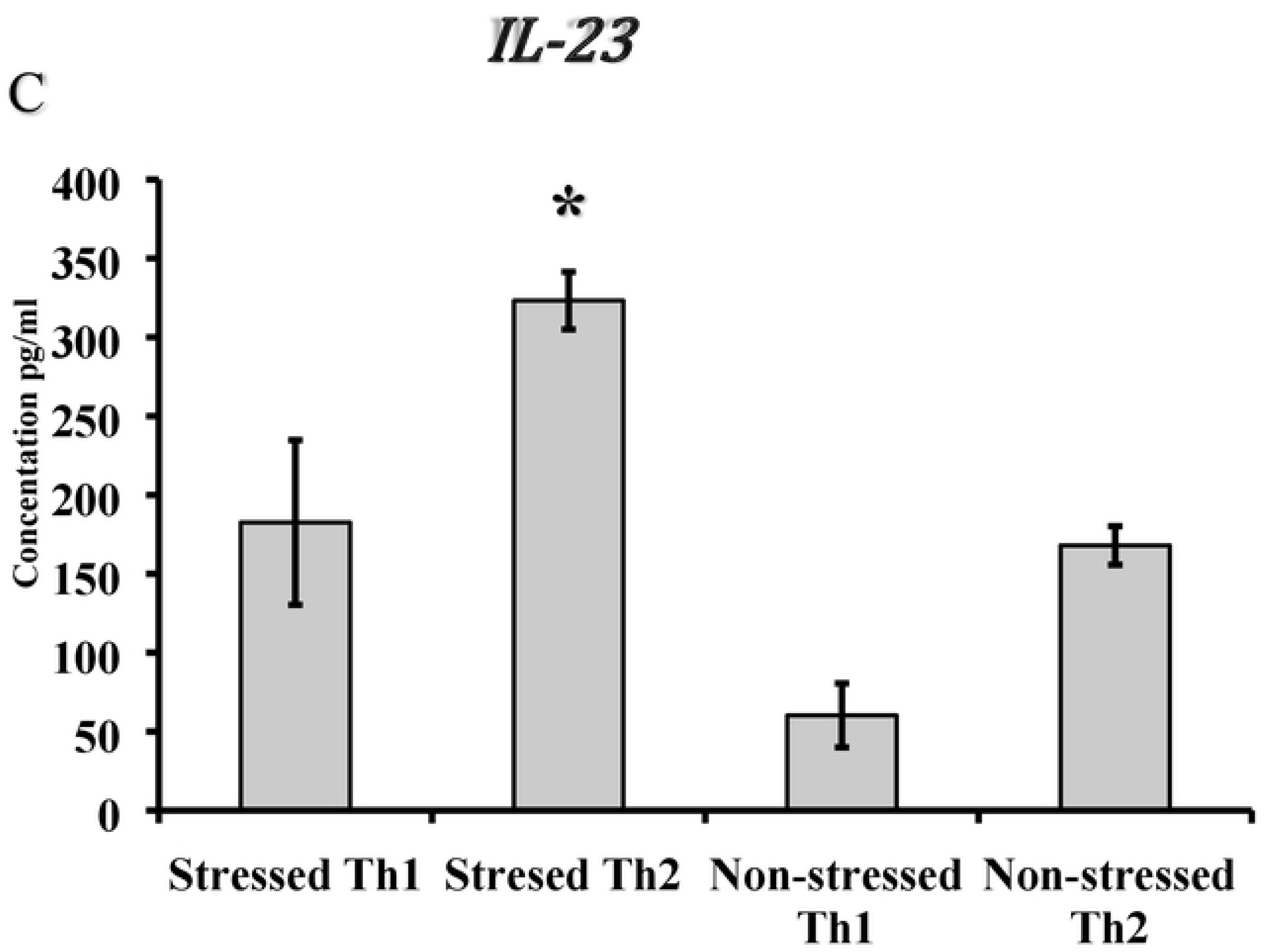
Western blot analysis of Type 1 and 2 T helper transcription factor GATA-3 and β-2 adrenergic receptor expression in CD4+T cells and Dendritic cells of stressed and non-stressed mice. Protein band was detected by Amplified Opti-4CN Substrate Kit fallowing manufacturer’s instructions purchased from Bio-Rad. Relative expression was calculated by the ΔΔCt method to compare expression levels of the same transcript in different samples or by the ΔCt method to compare expression levels of several transcripts in the same sample. Data shown are a representative or a mean +/_ SEM of two or more independent experiments ran on different dates.

### Polarization of T helper 1 and T helper 2 cytokine production in a cold-induced stress mouse model during *Chlamydia muridarum* genital infection

The purpose of this study was to evaluate the effect of cold-stress in altering the dynamics of Th1 and Th2 cytokine during *C. muridarum* genital infection. We hypothesized that cold stress alters the differentiation and variation in switching of cytokine productions in CD4+T cells. T cells isolated from the genital tract were proliferated as follows. For differentiation into Th1 cells, either Th1 cytokines and anti-IL-4 antibodies or Th2 cytokines and ani-IFN-γ antibodies were added to cell cultures in plates and stimulated by coating with mouse anti- CD3/CD28 antibodies. Production of cytokines were determined by ELISA. Western Blot analysis of Type 1 and 2 T helper transcription factors and β2-AR expression in CD4+T cells and dendritic cells. Our results showed no polarization of IFN-γ where increased IL-4 production was obtained which was associated with production of IL-4 and Stat-6, as evidenced by Western Blot analysis. These results indicate that cold-induced stress impairs the stimulation that drives polarization of T cells toward IFN-γ production but not IL-4 production. Cytokine IL-4 along with Stat-6 production is well known to support the Th2 arm of development. The study suggests that Th 1/ Th 2 cytokine imbalance may lead to severe chlamydia genital infection in the stress model.

**Figure 12.** Polarization of Naïve CD4+ cells in production of signature cytokines determined by ELISA. **(A)** IFN-γ, **(B)** IL-4, (**C**) IL-23. For differentiation of Th1 cells, mouse IL-12, mouse IFN-γ, human IL-2 and mouse anti-IL-4 were added to cell cultures in plates precoated with mouse anti- CD3/CD28 for stimulation. For Th2 cells, mouse IL-4, mouse anti-IFN-γ and mouse and anti-IL-12 were added in plates precoated with mouse anti-CD3/CD28 for stimulation. At day 3 cells were proliferated for additional 4 days in 15 ml of fresh culture medium containing the above Th1 and Th2 polarizing mouse and human cytokines and mouse antibodies. Production of IFN-γ, IL-4 and IL-23 cytokines were determined by ELISA Data shown are a representative or a mean +/_ SEM of two or more independent experiments ran on different dates.

### Interconnection of genes and their products that may be essential during stress and genital infection

Up-regulated or down-regulated gene expression profiles from stressed and non-stressed mice during *Chlamydia muridarum* genital infection were used to generate an interaction network to identify any other gene that could also be affected by their expression. As shown in **Figure 13**, gene nodes that are colored blue were down-regulated and up-regulated during stress and non-stressed conditions, respectively. Gene nodes in green were up regulated during stress and down regulated during non-stress conditions. The gene node that is colored red was down-regulated during both in stressed and non-stressed mice. The node shape shows the type of function that the genes play during stressful conditions. Nodes that are square are transcription factors, nodes that are circles are cytokines and nodes that are triangles are cells signaling receptors. Line color and shape indicates the regulation that their expression could play on other genes during stressful conditions. Green dashed lines indicate the genes could be down regulated and up regulated during stress and non-stressed respectively. Red arrowed lines show genes that could be up regulated and down regulated during stress and non-stressed respectfully.

**Figure 13:**
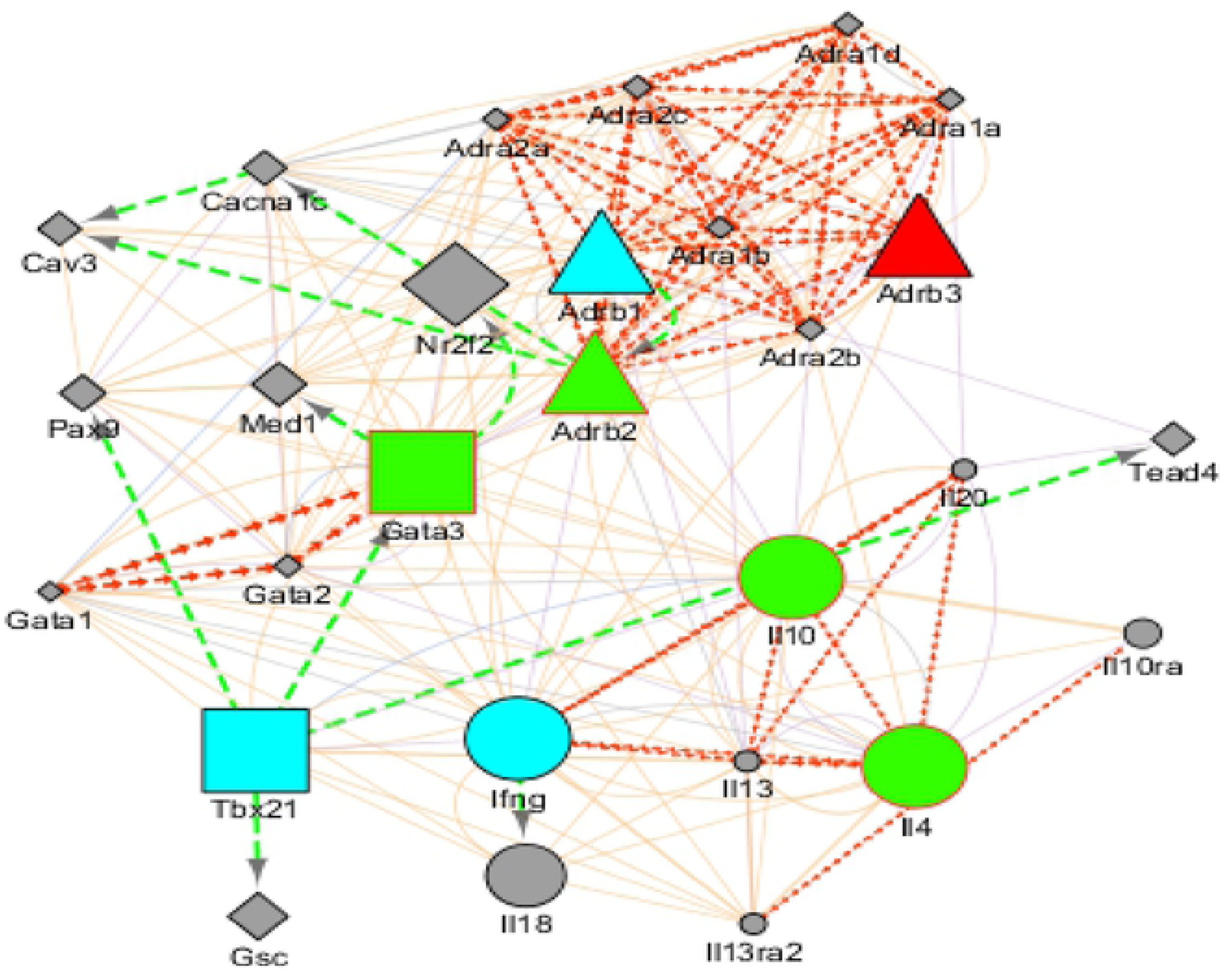
Cystoscope image of functional gene expression and potential regulation during *Chlamydia muridarum* genital infection in mice. The color of the nodes shows up- or down-regulation of genes in stressed mice during *C. muridarum* genital infection. Nodes shapes are made to show transcription factor, cell signaling receptors and cytokines. The edges are colored to show nodes that are affected by up-and down-regulation in stressed mice during *C. muridarum* genital infection.

## DISCUSSION

We showed previously that repeated exposure of mice to cold water leads to increased norepinephrine and epinephrine production in plasma, spleen, and genital tract lysates that are speculated in increasing the intensity of *C. muridarum* genital infection [59, 60]. Several studies have shown that CD4+ T helper (Th) cells have important protective roles during chlamydia genital infection, however, whether cold-induced stress contributes to imbalance of Th1 and Th2 cells in the mouse model is unknown. In the present study, the following investigations including gene expression profiles of beta-adrenergic receptor (β-AR) subtypes and differentiation of mixed T helper 1 (Th1) and T helper 2 (Th2) cell populations by analyzing the expression of T-bet and GATA-3 transcription factors were comparatively performed. Further, differentiation of bone marrow-derived dendritic cells (BMDCs) and their impact on production of Th1 type cytokine interferon gamma (IFN-γ)) and Th2-type cytokine (interleukin-4 (IL-4)), stimulation of *β*2-anadrenergic receptor (β2-AR) by pre-exposure to its agonist and antagonist and *in vitro* polarization of naïve CD4+ T cells toward the Th1 or Th2 subset in the presence of respective favoring cytokines and ant-neutralizing antibodies was examined.

Real-time quantitative PCR was used in gene expression profiling based on real-time fluorescence detection and QPCR target concentration measurements using threshold cycle (Ct). In most cases of our findings, the Ct is inversely proportional to the initial copy number and a direct relationship of a higher initial copy number to a lower threshold cycle was obtained. However, as shown in our amplification curves, reaction endpoint is not uniform even though the same amounts of cDNA from mRNA were being amplified. This observation is consistent with the fact PCR reaction efficiency can decrease during later amplification cycles as reagents are consumed and accumulation of inhibitors to the reaction do increase. We feel that these differences in final fluorescence values of samples are not related to the starting template concentrations.

Several studies indicate that the stress hormone norepinephrine (NE) suppresses the immune system through the stimulation of β2-AR expressed on various cell surfaces [39, 61, 62]. In this study, we determined the mRNA levels of *β*-anadrenergic receptor (β-AR) subsets in mixed-cell populations of CD4+ T cells isolated from the spleen or genital tract of stressed and non-stressed mice with/without *C. muridarum* genital infection. We have observed a dominant expression of β2-AR on CD4+ T cells isolated from the spleen and genital tract of stressed mice compared to that of non-stressed mice. However, we are not certain whether Th1 or Th2 cells is dominantly expressing β2-AR during the cold-induced stress application. On the other hand, stress treatment resulted in little or no gene expression of β1-AR and β3-AR subtypes in splenic CD4+ T cells, suggesting a differential expression of β2-AR subtypes by murine CD4+ T cells during stress and chlamydia genital infection. Numerous reports have shown that β2-AR is commonly expressed on the surface of immune cells, including Th1 cells, and when stimulated, it plays a big role in the inhibition of interleukin-12 (IL-12) and IFN-γ production [50]. However, researchers have shown that naïve CD4+ T cells and effector Th1 cells are responsible for expression of β2-AR but not Th2 cells [45, 46, 62]. Those researchers showed that Th2-promoting conditions at the early stage of proliferation increases the expression level of β2-AR mRNA, which decreased with increased IL-4 mRNA level in the system. Our results further show that β2-AR expression is significantly higher in stressed and infected mice when compared to stressed mice only, but the mechanisms that contribute to the stress-infection synergy which results in increased gene expression of the β2-AR subtype are unknown. These observations demonstrate that stress increases specifically the expression of β2-AR that is correlated significantly with increased C. *muridarum* infection, as evidenced by greater shedding in stressed mice than the control mice (60).

Because β2-AR is the most commonly expressed receptor, we tested whether a β2-AR agonist or β2-AR antagonist affects cytokine production. Our results show that β2-AR agonist decreases Th1 cytokine production probably by blocking T cell receptor (TCR)-mediated signaling, whereas β2-AR antagonist restores Th1 cytokine production. However, controversy exists on the effect of β2-AR agonist or β2-AR antagonist on Th2 cells, but studies have shown that β2-AR agonist modulates T cell functions via direct actions on both Th1 and Th2 cells (63). Overall, the results of β2-AR and NE interaction suggest that the differential receptor expression may alter the functional responsiveness of CD4+ T cells during cold-stress treatment and C. *muridarum* genital infection. This stress-infection model may provide a system to begin investigating the sequential relationship between stress or stress-infection combined synergy and to identify mechanisms by which cold-induced stress alters susceptibility to *C. muridarum* genital infection. Furthermore, understanding the mechanisms that regulate β2-AR expression may lead to development of methods for limiting Th1 or Th2 responsiveness to NE or β2-AR agonists that may lead to the development of CD4+T cells-mediated pathologies during chlamydia genital infection. From our study, we speculate that cold-induced stress increases expression of β2-AR in Th cells of stressed mice, which may play a significant role in alteration of the immune response. In contrast our findings suggest that col-induced stress has little effect on the pattern of gene expression of β1-AR and β3-AR subtypes in splenic CD4+ T cells. Exploring the expression of β2-AR in Th1 and Th2 under Th1- or Th2-deriving conditions may aid in finding the decreased IFN-γ and increased IL-4 production during cold stress treatment and *C. muridarum* genital infection.

Transcription factors expressed on T cells (T-bet) and GATA-binding protein-3 (GATA-3) are known to play key roles in the differentiation of naive Th1 cells towards Th1 or Th2 cells in human subjects and animal models [53, 54]. One of the aims of this study was to measure the gene expression of T-bet and GATA3 in CD4+ T cells using real time PCR. Our results show that T-bet gene expression was low in CD4+ T cells of stressed mice, whereas GATA-3 gene expression was markedly high in in CD4+ T cells, indicating cold-induced stressing results in restricted pattern of gene expression of transcription factors of Th1 and Th2 cells. However, other researchers reported that effector/memory CD4+ T cells that can produce either Th1 or Th2 cytokines commonly co-express T-bet and GATA-3 [64, 65].

It is well documented that naïve CD4+ T cells are able to produce different patterns of cytokines on their stages of differentiation, but how cold-induced stress affects their differentiation into Th cell subsets is not well defined. The effect of cold-induced stress on mRNA expression and secretion levels of IFN-γ, signature cytokine of Th1, or IL-4, signature cytokine of Th2, in CD4+ T cells of stressed and non-stressed mice with/without *C muridarum* was determined. Our study was based on non-polarizing conditions of CD4+ T cell differentiation that express their Th1 or Th2 cytokines and revealed that increased GATA-3 expression correlated with increased IL-4 expression/secretion was observed. In contrast, decreased expression of T-bet, correlated with decreased IFN-γ expression or secretion by splenic or genital tract CD4+ T cells is the hallmark findings of the study. The increased expression of GATA-3 and IL-4 secretion of our study favors the dominance of Th2 cells reflecting the switching of IFN-γ production by Th1 to IL-4 production by Th2 that can subsequently resulting in pathological development of chlamydia genital infection [13, 18]. These findings further confirm and extend previous reports from other research labs that stress treatment leads to up-regulated expression of GATA-3 and down-regulation of T-bet [17, 39, 54]. Although our findings show a high level of GATA-3 expression associated with IL-4 secretion and a low level of T-bet expression associated with IFN-γ production during cold-induced stress, cytokine production profiles of either Th1 or Th2 are not determined yet. Furthermore, the CD4+ T cell populations from the spleen and genital tract were not branded as naïve, resting memory or effector, but we speculate that IFN-γ and IL-4 are produced more during the effector stage of T cell differentiation than in the naïve and resting memory T cells.

We evaluated the relationship of gene expression and production of IL-4 and signal transducer and activator of transcription 6 (STAT6) protein expression in CD4+ T cells in the situation of cold-induced stress and chlamydia genital infection. Interestingly, significant secretion of IL-4 and STAT6 were shown in stressed mice compared to non-stressed mice. This positive correlation indicates that STAT6 - IL-4 pathway activation is perhaps involved in activating and regulating of Th2 immune response during cold-induced stress and chlamydia genital infection [39, 62].

Chronic stress generation adopted in our lab has demonstrated to exert a significant suppressive effect on Th1 cell cytokine production. We tested how long cold-induced stress can persist in our mouse model and results show that cold-induced stress has lasting effect on suppression of the immune system after the cessation of stressing process. This observation suggests to consider how the timing of stress is critical in promoting immune suppression in the implicated host. This slow immune response recovery shown in our study is in line with several studies demonstrating stress has long-lasting effects on the immune system that subsequently leads to infection [27, 28, 36, 40, 58].

The functional differences in Th1 and Th2 responses as protective and immunopathology, respectively during chlamydia infection was reported [18, 30]. In line with those reports, our data implicate that cold induced stress application changes the trends of Th1 and Th2 function that may play important role in reducing the host defense and promoting pathology during chlamydia genital infection. Differences found in Th1 and Th2 subset cytokine secretion in response to cold-stress treatment may provide insight onto the potential intrinsic immune equilibrium established between normal and stressful conditions. This study suggests the balance of Th1 and Th2 cytokines doubtless is important in resolving the immunopathogenesis of chlamydia genital infection during stressful conditions. The revealed changes in expression of transcription factors may indicate the imbalance between T-helper cells of the Th1 and Th2 type cells, which can be one of the causes for the development of pathology and infertility associated with chlamydia genital infection that was previous reported [60]. In contrast, recent studies revealed that *C. trachomatous* genital infection is mostly controlled by Th1 immunity, recent report claim that Th2 immunity has a major in controlling chlamydia genital infection in females [40].

Dendritic cells are significantly important in antigen presention activities to T cells for activation and differentiation of T cells to different subsets. However, the influence of cold-induced stress on differentiation and maturation of BMDCs and its activity in T cell function is not well defined. Moreover, little is known about how NE, β2-AR agonist, feroterol and the antagonist, ICI, 118 551 influences on the function of DCs in T cell differentiation. Because DCs and T cells express the β2-AR, which is the receptor of NE, we decided to examine the influence of feroterol and ICI, 118 551 on cytokine production in BMDCs and T cells. Further, we tested the function of matured BMDCs on CD4+ T cell activation in combination by co-culturing *in vitro*. Our hypothesis was that cold-stress treatment alters profiles of cytokine mRNA levels and cytokine secretion profiles during the differentiation and maturation of BMDCs or the function of BMDCs associated with CD4+ T cell activation. Our results revealed that pretreatment with feroterol resulted in a significant in stimulation of β2-AR in stressed mice, whereas ICI,118551, has played an inhibition role of β2-AR stimulation in stressed mice. We observed cold stress treatment results high secretion of IL-10 while reducing IL-12 suggesting a Th2 skewing potential of BMDCs pretreated with β2-AR agonist. Our results show agreement with previous studies that have shown NE plays important role in modulating T cell function through β2-AR activation, which is a key receptor expressed in many immune cells involved in mediating the immune system as reported previously [42, 44–48].

The co-culturing of DCs and CD4+ T cells showed decreased IL-12 and IFN-γ production but increased Il-17 and IL-10 secretion. However, we do not know whether matured BMDCs, T cells, or both are more responsible for the increased production of IL-12. Our findings confirm and extend previous observations of increased β2-AR expression Lipopolysaccharide (LPS)-stimulated DC indicated the production of lower amounts of IFN-γ and higher levels of IL-17 by CD4 T cells that decrease 12P70 section leading to a shift in the IL-12p70/IL-23 ratio [51, 61]. Based on our data, we feel that cold-stress leads to dominant Th2 cytokine production that probably is relevant to exacerbation of pathology during chlamydia genital infection. Our findings are in line with previous reports that have shown epinephrine-primed murine BMDCs facilitate production of IL-17A and IL-4 but not IFN-γ by CD4+ T cells (64) suggesting that the release of epinephrine and norepinephrine regulates the function of immune cells (66). Our findings are in line with previous reports that have shown feroterol inhibits the expression and production of IL-12 but promoted IL-10 production suggesting the stimulation of T lymphocytes by DCs [51, 61].

Because of the dichotomy of Th1 and Th2 cells remain as a major area to investigate in our model, polarizing cytokines were used for Th subset differentiator *in vitro*. To determine polarization of naïve CD4+ T cells IFN-γ and IL-4 are important sources for their own differentiation to Th1 and Th2, polarization of Th1 and Th2 toward their respective subsets. Our results showed no polarization of Th1 cells toward IFN-γ production of stressed mice was obtained where moderate Th2 polarization was obtained indicating L-4-producing CD4+ lymphocytes recovered from suppression from stress whereas Th1 function remained dampened. The conditions that trigger polarization Th1 maybe is not the only mechanism requiring IL-12 and STAT4 signaling, whereas for Th2 polarization, the major source is IL-4. Under the exogenous cytokine milieu used in this study, IL-4 was sufficient to derive Th2 differentiation possibly through STAT 6 activation, whereas poor polarization of IL-12 production associated with less activation of STAT 4 and ultimately less Th1 differentiation. To determine the involvement of T cell receptor (TCR) CD4+ T cells were stimulated by proliferation on ant- CD3/CD28 coated wells and observed differential of Th1 and Th2 cytokine production through indicating the mechanisms of stimulation are signals through the T cell receptor complex (signal 1) and other additional costimulatory (signal 2) as shown previously (66). Our findings implicate that stress effect selectively persists in the differentiation of Th1 compared to that of Th2 development but further study is warranted about polarization of Th1 and Th2 cytokines.

Studies in our and other labs have demonstrated that cold-stress treatment to mice brings changes that impact the activity of the immune system. However, it is not yet known how cold stress exposure suppresses or enhances the immune system activities. It is suggested that the sympathetic nervous system activation leads to the release of norepinephrine and epinephrine that interact with the immune system cells through binding to adrenergic receptors on their surfaces. Although there are multiple mechanisms that sympathetic nervous induced system of regulation, β2-AR signaling may contribute to increased immunosuppression during stress chlamydia genital infection. We feel that the major stress response is norepinephrine production which plays a key role in regulation CD4+ T cells and DCs function. To clarify the role of β2-AR signaling on the modulation of DC function, we compared the effect of NE, forterol and IC 118,551 on maturation, cytokine production assess the ability of these BMDCs to induce/derive T helper differentiations. We observed cold stress treatment results high secretion of IL-10 while reducing IL-12. This phenomenon corresponded with a Th2 skewing potential of BMDCs pre-exposed to β2-AR agonist. We were able to demonstrate the dichotomy between IL-12 and IL-4 in the presence of β2-AR agonist and antagonist. Although *C. trachomatis* serovar D infection in mice was used, it was reported the Th1 subset with T-bet and IFN-γ production dominated the upper genital tract compartment female mice whereas the lower genital tract has IL-10 and GATA-3 suggesting the dominance of Th2 [40, 63]. Based on our present findings of th1 and Th2 cytokine production profiles in our stress model, we feel that cold-induced stress leads to Th2 dominated conditions which favor enhanced development of immunopathogenesis during *C. muridarum*. Although previous studies showed that b2-AR is absent in Th2 cells (47), other studies demonstrated that β2-AR agonists modulate T cell functions via direct actions on Th1 and Th2 cells, where CD3 + CD28-stimulatd IL-13 (Th2 cytokine) and IFN-g and IL-2 (Th1 cytokines) production were inhibited by β2-AR agonist (40,63). Our study on mixed population of Th1 and Th2 resulted in the inhibition of Th1 cytokines but not Th2 cytokines thus our findings underscore the importance of further study on the action of β2-AR agonists during stressful conditions.

Nowadays, datasets are available in public archives such as the Gene Expression Omnibus (GEO) and cystoscope [69]. Although the scope of this study was not to undertake detailed analysis of biological processes and signaling pathways of genes or their products, we inserted our findings into global datasets and constructed a network and identified the genes shown in Figure 13. We identified a group of proteins that have interconnection during chlamydia infection of stressed model identified several candidate targets for further fundamental experimental research.

In summary, we undertook a detailed assessment of how cold-induced stress modulates a mixed population of CD4+ T and BMDC cells by examining expressions of β-AR subsets, T cell transcription factors, mRNA levels and secretion of hallmark cytokines of Th1 and Th2 cells, stimulation of β2-AR *in vitro* setting using a physiological applicable concentration of NE, β2-AR agonist, or antagonist. Further, polarization toward Th1 or Th2 of a mixed population of naïve CD4+ T cells from stressed and non-stressed in the presence of anti-CD3 stimulation and polarizing milieu. A marked increase in expression of GATA-3 associated with elevated expression and section of IL-4 compared to decreased expression of T-bet and expression, IL-12, and secretion of IFN-γ in CD4+ T cells of stressed mice compared to that of non-stressed mice was obtained. Further, co-culturing of mature BMDCs and naïve CD4+ T cells of stressed mice resulted in decreased expression and secretion of IL-12 whereas increased production of IL-4, IL-10, and IL-17 in culture supernatants of the co-culture suggests that stimulation of β2-AR receptor leads to the increased production of Th2 cytokines and down-regulated CD4+ T cells of stressed mice compared to that of non-stressed mice was obtained. The revealed changes in expression of transcription factors and cytokine production dynamics may indicate cold-induced stress treatment leads to the imbalance between Th1 and Th2 type cells, which could be one of the causes of development of pathology and infertility associated with chlamydia genital infection.

For the first time we were able to present evidence that cold-induced stress leads to the overexpression of GATA-3 in contrast to T-bet increased mRNA level and secretion of IL-4 in contrast to diminished secretion of IFN-γ in CD4+ T cells and BMDCs. Based on our results revealed that there maybe is a shift from a Th1 proinflammatory response toward anti-inflammatory th2 response as characterized by increased levels of GATA-3, IL-4 and STAT-6. This observation suggests that while it is known that CD4+ Th1 are essential in the clearance of chlamydia, the increased production of IL-10 and IL-17 in our stress model may lead to the establishment of chronic chlamydia infection (66-68). We believe that this study may offer important direction for further investigation of the mechanisms of stress on chlamydia genital infection and immunity in humans. We feel that this study is important because psychological or physical stressors may lead to increased stress hormone production and have profound effects on health, including prevalence of chlamydia genital infection. Evidence shows that these stressors generally are greater in populations of lower socioeconomic status, in which there are increased health concerns. The present finding, taken together within previous observations [59, 60] indicate that cold-water induced stress increases the intensity of chlamydia genital infection in mice. The long-term goal of our study is to employ the cold-water induced stress in a β2-AR KO mouse model, and the hypothesis to be tested is: cold water-induced stress leads to NE modulation of the immune response against *C. muridarum* genital infection by enhancing the production of immunopathogenic cytokines that result in disease sequelae.

## ACKNOWLEDGEMETS

We are grateful to Dr. Joseph Igiesteme; at the Centers for Disease Control and Prevention (CDC), for his expertise and consultation in the Chlamydia project and for providing us Chlamydia stock cultures throughout the study. We are also thankful to Dr. Guy Sims’s Title III Office at Bluefield State College for the financial assistance to prepare the manuscript. We also are very thankful to Dr. Julie Kalk, for editing our complete manuscript.

